# A physical theory of organismal metabolism across life

**DOI:** 10.64898/2026.09.03.749078

**Authors:** Peter F. Pelz, Tobias Meck

**Affiliations:** Technische Universität Darmstadt, 64287 Darmstadt, Germany

## Abstract

A *“*fundamental *but unanswered biological question”* is how much energy life forms expend per unit mass and time to remain alive [1]. The West–Brown–Enquist (WBE) theory explains Kleiber’s 3*/*4-power law [2] through complex vascular networks [3–14], but assumes *“size-invariant terminal units”* and predicts neither absolute rates, small-organism linearity, nor group-specific levels. Yet capillary dimensions vary with body mass [15] and obey Murray’s optimization principle [16]. Furthermore, 3*/*4-power scaling occurs in planarians with a simple tubular metabolic system [17]. Here we extend Krogh’s capillary–tissue unit [18] to an optimized mesoscale oxygen-transport system, the Metabolic Holon (MH). Krogh’s geometry, Murray’s minimum-work principle, and optimal oxygen utilization determine its geometry and operating state. Kleiber’s asymptote thus emerges from an optimized Krogh unit, without requiring complex vascular-network architecture; repetition and within-group body-mass-invariant topology determine organismal metabolism. Without fitting metabolic-rate data, MH predicts absolute rates across 18 orders of magnitude (*R*^2^ = 97.7 %), including linear scaling and group-specific Kleiber asymptotes. Independent dimensional analysis recovers these asymptotes. The theory also predicts absolute mammalian heart rates without calibration; absolute lifespan additionally requires an empirical lifetime-heartbeat constraint. By representing evolutionary selection and adaptation through mesoscale optimization, the theory explains why *“mass-specific metabolic rates are strikingly similar across life’s major domains”* [1].

---

From bacteria to whales, aerobic organisms must supply oxygen at the rate required by cellular metabolism. Since Kleiber identified the three-quarter-power relation between metabolic rate and body mass in 1932 [2], its physical basis has remained the subject of continuing debate [1, 3–14, 19–21].

Organismal metabolism undergoes a fundamental transport transition: metabolic rate scales linearly (*m*^1^) in small, diffusion-dominated organisms such as protists, but continuously approaches Kleiber’s 3/4-power law as convective transport becomes necessary. At the same time, absolute metabolic levels shift systematically across organizational groups, from simple tubular planarians to vascularized ectotherms and endotherms. Across the observed 18 orders of magnitude in body mass, no single physical framework currently accounts simultaneously for small-organism linearity, the transition to the 3/4-power regime, and the group-specific absolute metabolic levels (Fig. 1; [1, 17, 22, 23]).

**Figure 1:**
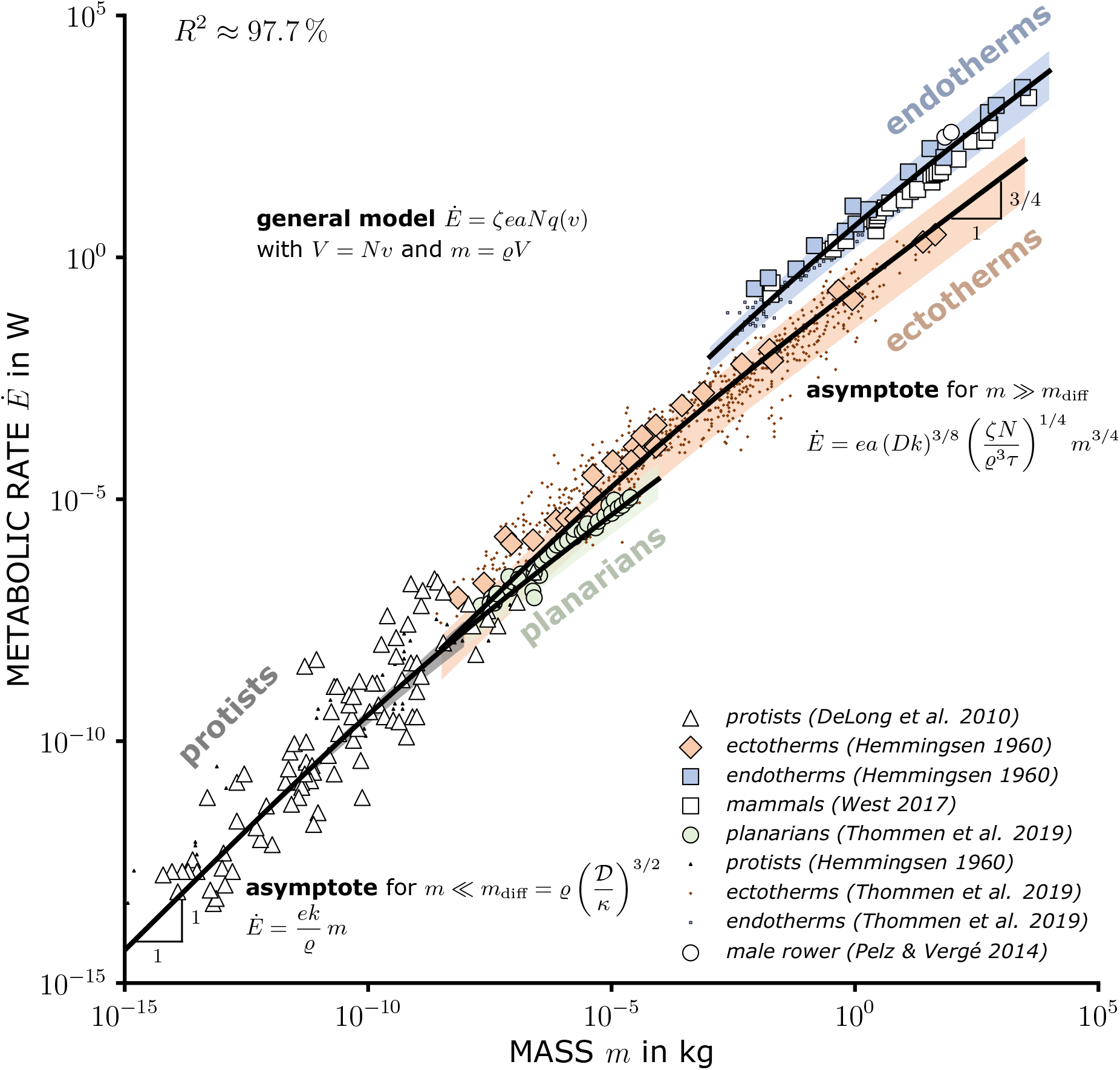
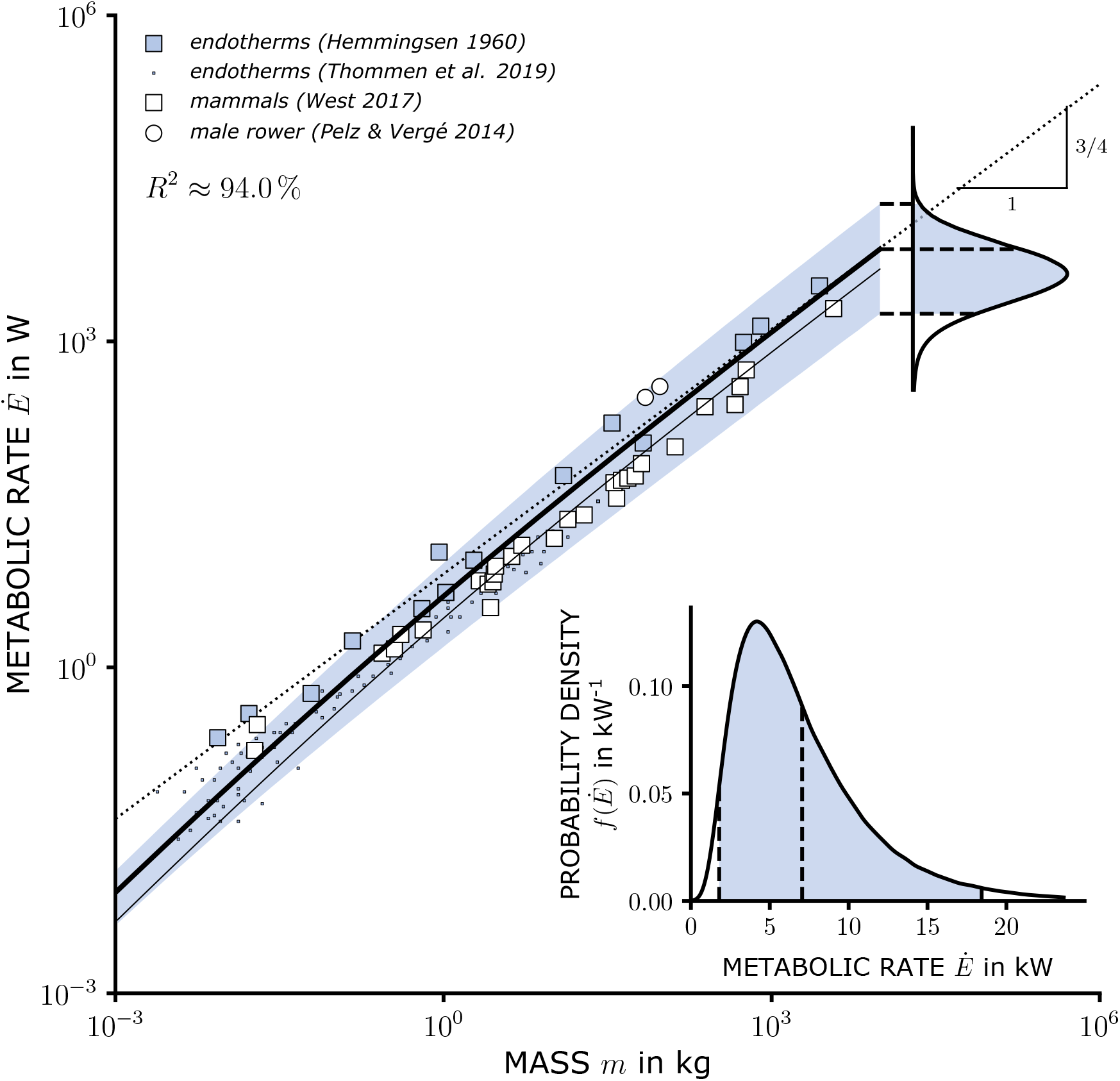
Predicted metabolic rates across metabolic regimes. **a**, Thick solid line (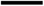): predictive mean of the general MH model (Eq. 8); shaded region: central 95 % parameter-uncertainty interval. No parameter is fitted to metabolic-rate observations (pooled logarithmic *R*^2^ = 97.7 %; 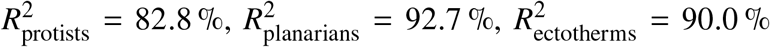 and 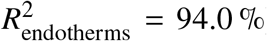).Symbols denote observations from [17, 22, 23, 56, 57]. **b**, Thick solid line (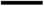): predictive mean of Eq. 8; thin solid line (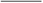): deterministic solution of Eq. 8; dotted line (····): MH Kleiber asymptote (Eq. 9). The departure at smaller body mass shows the transition toward the asymptotic regime.

The missing link lies between cellular respiration and whole-organism metabolism. We extend Krogh’s capillary–tissue unit [18] through Murray’s minimum-work principle [16] and optimal oxygen utilization into an optimized mesoscale system, the Metabolic Holon (MH). Figure 2 illustrates its geometry and operating state. Repetition connects the MH to organismal metabolism while separating mesoscale transport physics from organismal topology.

**Figure 2:**
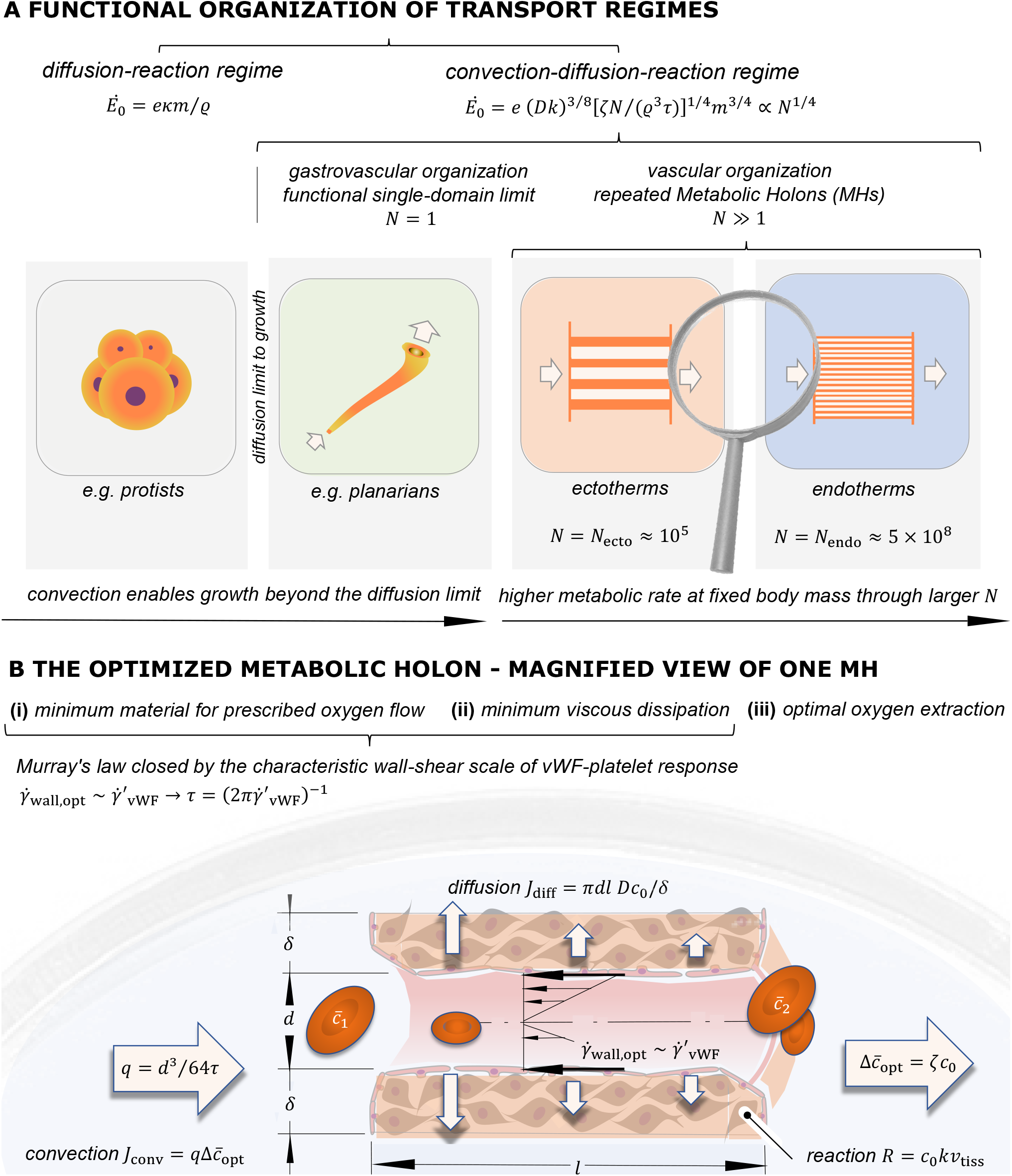
Cross-scale organization of metabolic transport and the optimized Metabolic Holon. **(A)**Cellular demand, the MH mesoscale, and organismal repetition define the cross-scale structure; *N* = 1 is the single-domain limit and *N* ≫ 1 the repeated topology of vascularized organisms. **(B)** Vascular realization of one optimized MH, combining Murray’s law, optimal oxygen utilization, and convection–diffusion–reaction balances (Eqs. 1–7; Murray: Eq. 4). Colors correspond to Fig. 1.

## Metabolic scaling theories and the MH framework

Cellular models explain diffusion-dominated linear scaling (*m*^1^), whereas network theories, most notably West–Brown–Enquist (WBE), derive Kleiber’s 3/4-power law from complex vascular-network architecture [3–5, 7–9, 13]. WBE assumes *“size-invariant terminal units”* [3, 10, 11, 13], but predicts neither absolute metabolic rates, small-organism linearity, nor group-specific levels. Yet capillary dimensions vary with body mass [15] and are governed by Murray’s optimization principle [16]. Furthermore, 3/4-power scaling occurs in planarians with a simple tubular metabolic system [17]. There is a striking irony in the search for the physical origin of Kleiber’s law: WBE seeks its explanation at the *organismal macroscale*—in large, complex vascular networks—while assuming a body-mass-invariant capillary diameter. At the opposite end, Thommen *et al*. [17] explain its occurrence in small, simple planarians at the *cellular microscale*, through size-dependent changes in cellular energy storage. The MH framework instead identifies the decisive optimization pressures at the *mesoscale*, where Krogh’s tissue cylinder, optimized through Murray’s minimum-work principle [16] and optimal oxygen utilization, yields the 3/4 exponent within a single functional unit. Crucially, the MH framework represents evolutionary selection and adaptation through mesoscale optimization while coupling all three scales: cellular physiology determines transport and reaction properties, whereas organismal repetition and topology determine group-specific metabolic levels. The resulting theory predicts absolute metabolic rates, the transition from linear scaling to Kleiber’s law, group-specific metabolic levels, and heart rates, while lifespan additionally requires one empirical lifetime-heartbeat closure. The final irony is historical: the geometric foundation of this cross-scale physical theory has been before us since Krogh’s seminal work in 1919 [18]—precisely at the level where the original WBE framework assumes *“size-invariant terminal units”*.

Thermodynamic surface-to-volume approaches instead relate scaling to heat production and surface heat loss [20]. At the tissue scale, however, thermal diffusivity exceeds oxygen diffusivity by roughly two orders of magnitude, 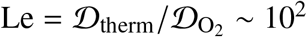 [24], and blood further redistributes metabolic heat convectively. Oxygen transport therefore provides the slower tissue-scale transport constraint considered here; Galileo’s 2/3 surface-to-volume scaling remains a physical constraint but need not be the dominant one.

The seven physical relations of the MH model connect cellular oxygen demand with optimal oxygen transport and geometry. Its general solution predicts the linear regime, the transition to convection, and group-specific Kleiber asymptotes in absolute units. Transformation invariance and complete similarity constrain the body-mass dependence of organism-level parameters A and *N*; independent anatomical and physiological observations constrain their group-specific magnitudes. Neither metabolic-rate nor heart-rate data are used for calibration; only absolute lifespan requires an empirical lifetime-heartbeat constraint. By extending Murray’s minimum-work principle, the MH framework inherently incorporates its optimal flow–diameter relation and predicts the observed capillary scaling (Supp. S1; [15, 16, 25, 26]). By representing evolutionary selection and adaptation through constrained model-based optimization, the framework provides a physical basis for understanding oxygen consumption, respiratory symmorphosis (Supp. S2), and evolutionary innovation.

### What happens when organisms grow?

Small organisms can supply their cells by diffusion alone. Their oxygen transport and cellular oxygen consumption are governed, respectively, by the molecular diffusion coefficient D of oxygen in tissue and the cellular oxygen-consumption rate constant *κ*. Cellular biochemistry enters the theory only through this coarse-grained oxygen demand: at leading order, *κ* phenomenologically represents an effective first-order oxygen-consumption rate constant rather than an intracellular reaction network. Together with tissue density *ϱ*, these quantities define the organism-scale diffusion mass *m*_diff_ := *ϱ* (*D*/*κ*)^3/2^.

For *m* ≪ *m*_diff_, the organism-scale Damköhler number represents the ratio of diffusion and reaction timescales and is

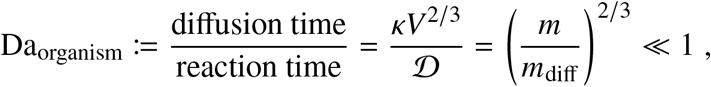

where *V* denotes organism volume and *m* = *ϱV* body mass. Diffusion therefore equilibrates oxygen faster than cellular reaction, leaving the concentration field nearly homogeneous and metabolism proportional to body mass (*m*^1^). As *m*/*m*_diff_ approaches unity, Da_organism_ approaches one; for *m*/*m*_diff_ > 1, diffusion becomes too slow to prevent anoxia, and further growth requires convection coupled to diffusion and cellular reaction within a bounded mesoscale domain. Repetition of these mesoscale domains builds the organismal macroscale (Fig. 2).

## The Metabolic Holon—the missing link

Krogh’s tissue cylinder provides the geometric substrate for the vascular realization of the Metabolic Holon [18], but anatomy alone does not define the functional unit. More generally, a tubular convective–diffusive supply domain becomes an MH only when, at a prescribed MH volume *v* := *V*/*N*, its optimal aspect ratios and operating state are determined by the transport optimization developed below (Fig. 2). In planarians, the gastrovascular tubular system provides a special single-domain realization (*N* = 1); in vascularized ectotherms and endotherms, the corresponding tubular element is a capillary, and repeated MHs form the microvascular supply system. Recognizing that real tissues exhibit spatial variations in structure and capillary density, these spatial heterogeneities are coarse-grained into an effective homogeneous representation characterized by the mean MH number density *n* := *N*/*V*.

### Why does Krogh’s tissue cylinder become a holon?

Arthur Koestler coined the term *holon* for an entity that is simultaneously a whole and a part [27]. The MH realizes this duality directly: a single domain can constitute the complete functional supply system in planarians (*N* = 1), whereas repeated MHs form the microvascular supply system of ectotherms and endotherms. The relation *V* = *Nv* maps the unit to the whole organism (Fig. 2).

### Geometry

In its vascular realization, the MH contains a capillary of diameter *d* and an effective arteriole-to-venule path length *l*. More generally, *d* and *l* denote the characteristic transverse and axial dimensions of the tubular convective–diffusive domain. For vascular MHs, *l* represents the functional path length of the unit rather than a simple geometric multiple of a single capillary segment; direct three-dimensional microvascular reconstructions reveal interconnected pathways, loops, and substantial tissue- and species-dependent variation (Fig. 2B; [28–30]). In the vascular geometry, adjacent capillaries have a center-to-center spacing of *d* + 2*δ*, where *δ* denotes the effective tissue diffusion distance. Together, the central capillary and its surrounding tissue volume *v*_tiss_ define the total MH volume *v*.

## Similarity constrains growth and topology

As shown below, the optimization of a single MH already yields Kleiber’s 3/4-power asymptote, independently of network architecture—whether simple, as in planarians, or complex, as in mammals. The decisive physical mechanism therefore lies not in the detailed architecture of the distribution network, but in the functional units where oxygen supply meets cellular demand. Yet the metabolic rate of a single MH determines that of the entire organism only in the single-domain case (*N* = 1), as in planarians. For organisms with repeated MHs, absolute metabolic rates additionally depend on their functional topology *N*.

We therefore first constrain organismal topology through physical similarity, using a coarse-grained, effectively homogeneous representation, before deriving the optimized MH and its absolute metabolic predictions. *The central biological question is whether growth requires organisms to create progressively more functional oxygen-supply domains, or whether an established set of domains simply enlarges with the organism*.

Cross-scale coupling connects the cellular microscale, MH mesoscale, and organismal macroscale through the transfer of physical information.

First, information from two directions converges at the MH mesoscale. From the cellular microscale, physicochemical and physiological properties provide the constitutive quantities *D, κ*, the characteristic timescale *τ* entering Murray’s flow–diameter relation, and the dimensionless relative oxygen capacity of blood *ζ*. From the organismal macroscale, body mass *m* and functional topology *N* determine the MH volume *v* = *m*/(*N ϱ*). Together, these quantities determine the geometry and operating state of the MH.

Second, the MH framework takes a methodological approach in which *evolutionary selection and adaptation are represented through model-based optimization*. An optimization problem can be approached by heuristic search or exact analytical methods. Natural selection and adaptation are interpreted as evolutionary heuristics, whereas the MH framework determines the optimum analytically. The MH optimization has a unique, robust solution, independent of the path taken to reach it. Our hypothesis is therefore simple: *one optimization problem, two solution approaches, one physical optimum*.

Crucially, the Pareto parameters weighting the competing optimization pressures are not freely adjustable. At the mesoscale, Murray’s optimization (Supp. S1) balances vascular material costs against viscous dissipation through *ε*_MH_. At the organismal macroscale, respiratory optimization (Supp. S2) balances respiratory and vascular material costs against operational transport costs through *ε*_RS_. Cross-scale coupling determines their physiological meaning and absolute values hierarchically: *ε*_MH_ through microscale blood rheology, and *ε*_RS_ through the MH turnover time.

From the organismal macroscale, the dimensionless quantities A, *B*, and *N* characterize the stroke-volume ratio, relative blood volume, and functional topology, respectively. In mammals, for example, *V*_heart_/*V* ≈ 6 × 10^−3^ and *V*_stroke_/*V* ≈ 10^−3^, yielding A := *V*/*V*_stroke_ ≈ 10^3^ [31, 32]. Similarly, the relative blood volume *B* := *V*_blood_/*V* ≈ 7–9 % in endotherms exhibits no systematic body-mass dependence [14, 32].

To distinguish between these possibilities, dimensional analysis determines how cellular-microscale properties and organism-level topology constrain the MH mesoscale under the fundamental requirement that physical laws remain invariant under transformations of base units [33], prior to specifying vascular architecture. The biological consequence is simple: if the dimensionless organization defining a metabolic similarity group is preserved during growth, increasing body size need not require an increasing number of functional oxygen-supply domains.

We first apply this axiomatic method to the well-established body-mass invariance of the stroke-volume ratio A and the relative blood volume *B* (Supp. S2), and then extend the same formal reasoning to functional topology. This yields the central prediction that the group-specific number *N* of repeated MHs is likewise body-mass invariant, *N* ∝ *m*^0^.

We establish this result in three systematic steps: (1) physical similarity provides the axiomatic basis, (2) morphogenesis offers a possible mechanistic realization, and (3) independent observations provide empirical evaluation (Table 1). The 11 empirical comparisons explicitly distinguish inference (I) of the group-specific magnitude of *N*, consistency tests (C) of its mass invariance, and prediction tests (P) of downstream physical consequences.

**Table 1:**
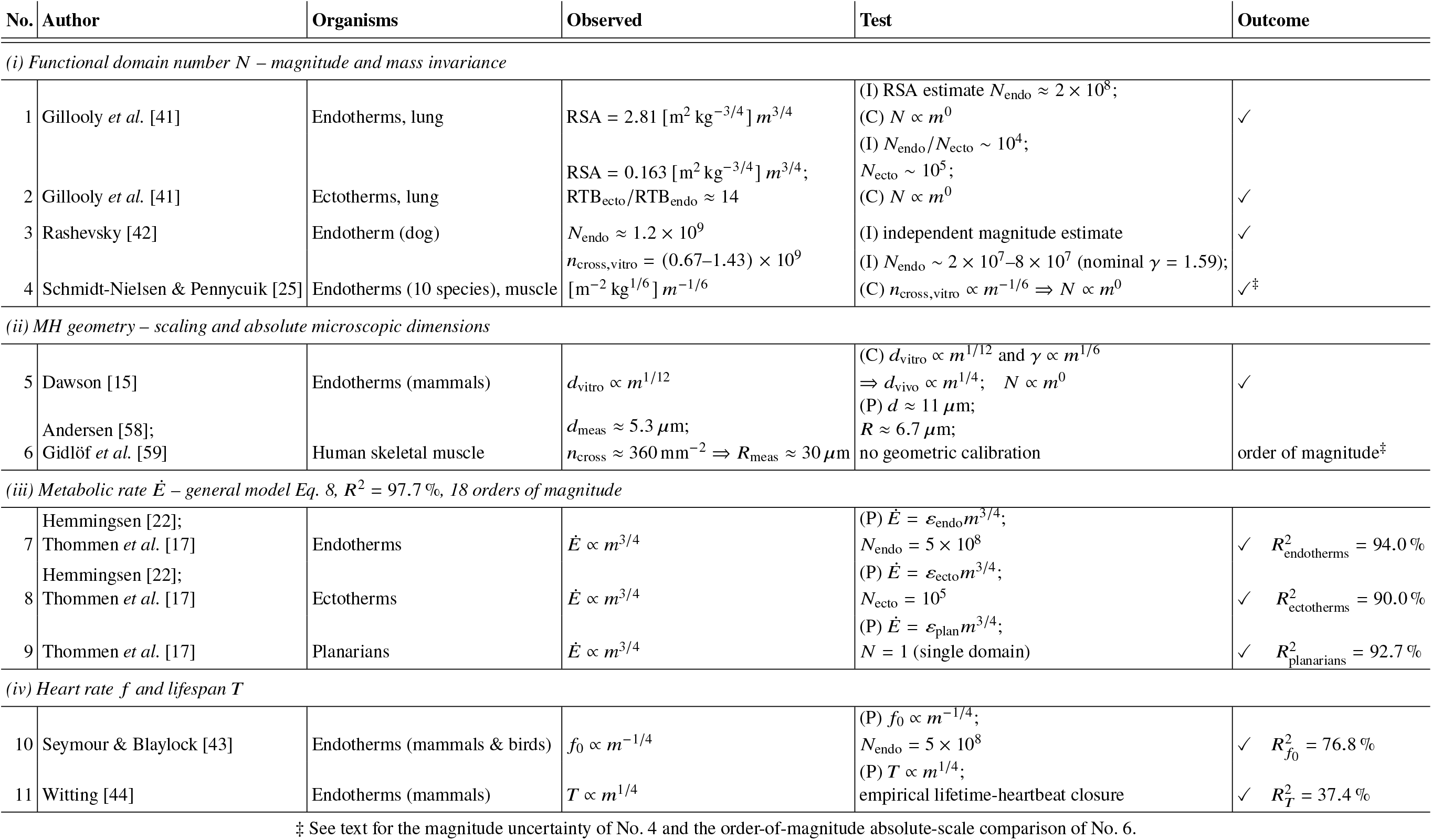
Convergent quantitative tests of the MH framework and the mass-invariant functional oxygen-supply domain number *N*. Eleven empirical comparisons across geometric, physiological, and organism-level observations probe complementary levels of the theory: model-based inference (I) of the group-specific magnitude of *N*, consistency tests (C) of *N* ∝ *m*^0^, and prediction tests (P) of downstream consequences once *N* has been established; lifespan uses one empirical lifetime-heartbeat closure for its absolute scale.

### (1)Axiomatic basis: similarity within metabolic groups [33–36]

Physical relations must be invariant under transformations of the base-unit system. Bridgman’s postulate of the *absolute significance of relative magnitude* expresses that relative measures based on scales intrinsic to the problem, such as *m*/*m*_diff_, are independent of any arbitrary choice of units and therefore have absolute physical meaning [34]. Buckingham’s Π theorem gives the mathematical consequence of this invariance: the number of independent dimensionless products of powers of the dimensional quantities equals the number of dimensional quantities in the relation minus the rank of their dimensional matrix.

Before specifying vascular geometry, we ask which combinations of the biological and physicochemical variables can carry a body-mass dependence and, more importantly, which must remain body-mass invariant. For the stroke-volume ratio as for the relative blood volume, Buckingham’s Π theorem reduces

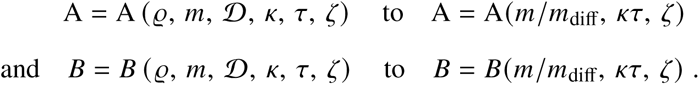

Two entities are *physically similar* when the dimensionless products relevant to the problem are equal [35]. In the order-of-magnitude theory used here, this condition is applied in coarse-grained form: these dimensionless products remain of the same order of magnitude and exhibit no systematic scaling with body mass. Applied to organisms, entities satisfying this condition belong to the same *metabolic similarity group*. For the data considered here, we distinguish four such groups: diffusion-dominated protists, single-domain planarians (*N* = 1), ectotherms, and endotherms; these groups characterize coarse-grained transport and topology rather than taxonomic rank.

Physical similarity may also be attained asymptotically. A dependent dimensionless quantity may initially depend on other independent dimensionless products, Π_0_ = Π_0_(Π_1_, Π_2_, …). If, as one of these products tends to zero or infinity, Π_0_ approaches a finite non-zero limit independent of that product, the latter drops out of the leading-order relation. This asymptotic independence is referred to as *complete similarity* [35, 36].

Within a metabolic similarity group, *κτ* and *ζ* remain body-mass invariant. Consequently, the body-mass dependence of both A and *B* is confined to the similarity variable *m*/*m*_diff_. Assuming complete similarity in the sense of Barenblatt [35, 36] for *m*/*m*_diff_ → ∞ gives

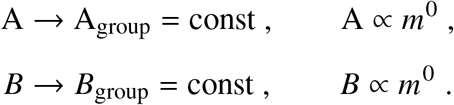

The constant may differ among metabolic similarity groups (endotherms and ectotherms; Figs. 2A and 1) because each group represents a distinct similarity branch. The observed order-of-magnitude invariance of A and *B*, without systematic body-mass dependence within metabolic similarity groups, is therefore consistent with asymptotic complete similarity [14, 31, 32].

The same similarity argument that provides an axiomatic basis for the well-established relations A ∝ *m*^0^ and *B* ∝ *m*^0^ applies to the MH number *N*. Applying the invariance requirement through Buckingham’s Π theorem reduces the relation from

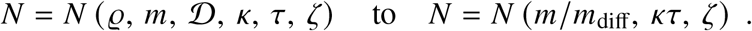

Because *κτ* and *ζ* are body-mass-invariant physiological quantities within a metabolic similarity group, the body-mass dependence of *N* is likewise confined to *m*/*m*_diff_. For *m*/*m*_diff_ → ∞, complete similarity with respect to *m*/*m*_diff_ gives

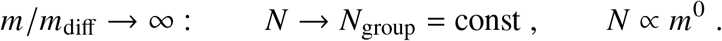

Although counterintuitive from organismal size alone, this prediction does not imply invariant vascular geometry: within a similarity group, increasing body size is accommodated by rescaling the geometry and volume of the repeated MHs rather than by a systematic increase in their number.

For A, body-mass invariance is empirically established; for *N*, it is a theoretical prediction. The group-specific magnitude *N*_group_ is inferred from geometry and physiology, not metabolic rates. Growth changes MH geometry while preserving *N*; evolutionary innovation can establish a different topology (Fig. 2A).

### (2) Possible mechanistic realization: morphogenetic fixation of topology

Two distinct phases of organismal development provide a possible biological mechanism by which this topological mass invariance can be realized. During early embryogenesis, a Turing-type reaction–diffusion system establishes a spatial pattern within the growing developmental domain *V* [37]. In the classic uncoupled Turing mechanism, the selected intrinsic wavelength *λ* is dictated by intrinsic reaction– diffusion kinetics rather than by domain boundary conditions; if active pattern formation persisted throughout all stages of organismal growth, the total number of pattern elements would scale continuously with organism size.

However, studies on morphogen scaling and growth–pattern coupling demonstrate that active pattern generation can be followed by geometric scaling of an established template [38–40]. Once developmental topology becomes morphogenetically fixed, subsequent growth expands the pre-existing domains rather than creating additional units de novo. Consequently, their characteristic spatial spacing scales directly with overall organismal length, *λ* ∝ *L* ∝ *V* ^1/3^, yielding

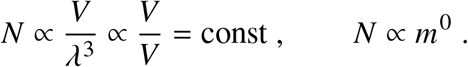

Early morphogenetic fixation could thus establish *N*_group_, with subsequent growth enlarging individual MHs. This developmental hypothesis is not required by the similarity derivation.

### (3) Empirical evaluation: inference (I), consistency (C), and prediction (P) [15, 17, 18, 25, 41–44]

Table 1 and Supplement S5 evaluate the predicted mass invariance and independently constrain the group-specific magnitude of *N*. Respiratory surface area (RSA) [41] is linked to the aggregate microvascular blood–tissue surface by the *respiratory-surface matching relation*, derived from organism-level optimization (Supp. S2; Extended Data Fig. 2):

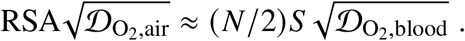

Observations give RSA ∝ *m*^3/4^ [41], while the MH model gives *S* ∝ *v*^3/4^ = (*V*/*N*)^3/4^ (Extended Data Fig. 3). Therefore, their combination yields *N* ∝ *m*^0^, consistent with the similarity prediction. The RSA prefactor then constrains the absolute magnitude of endotherm topology to the 10^8^ scale.

Measured endotherm–ectotherm surface and barrier ratios similarly constrain ectotherm topology to the 10^5^ scale (Extended Data Fig. 3). Independent observations of capillary density, capillary-diameter scaling, and Rashevsky’s whole-organism estimate provide complementary constraints on the magnitude and mass dependence of *N* (Extended Data Fig. 4; [15, 25, 42]). The RSA-based inference gives *N*_endo_ ≈ 2 × 10^8^, while Rashevsky’s independent whole-organism estimate gives approximately 1.2 × 10^9^. Their geometric mean, *N*_endo_ = 5 × 10^8^, together with *N*_ecto_ = 10^5^, provides the group-specific inputs at the order-of-magnitude resolution of the theory. The remaining datasets test MH geometry, metabolic rate, heart rate and lifespan.

Across all 11 inference, consistency, and prediction comparisons (Table 1), the evidence converges on *N* → *N*_group_ ∝ *m*^0^, making a substantial residual body-mass dependence of *N* very unlikely.

The MH framework is open to *falsification at two levels*: through its *consistency with physical principles*, including Murray’s law, blood rheology, and cellular oxygen transport and consumption; and through *empirical tests of its predictions* for microvascular morphology, organismal metabolic rates, scaling asymptotes, group-specific metabolic levels, and heart rate. Despite its coarse-grained, order-of-magnitude description, the strength of the framework lies in connecting these diverse observations through a common physical mechanism rather than fitting metabolic-rate or heart-rate data. Lifespan provides a further test of the predicted scaling, while its absolute magnitude requires an empirical lifetime-heartbeat constraint.

## Mesoscale optimization of the MH determines metabolism

For a prescribed MH volume *v* = *V*/*N*, oxygen transport, cellular reaction, and geometry define a model-based optimization at the mesoscale. Three competing physical objectives determine MH shape and operation:

i. **minimizing vascular material volume** for a prescribed oxygen flux,
ii. **minimizing viscous dissipation**, and
iii. **maximizing oxygen extraction** while maintaining distal tissue supply.

The first two objectives yield Murray’s Pareto-optimal flow–diameter relation

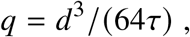

where *τ* is the characteristic transverse hydrodynamic timescale (Supp. S1; [16]). Murray’s law gives the wall shear rate 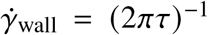. Independent observations of vWF-mediated platelet adhesion identify a characteristic high-shear regime at 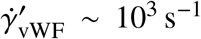 (Extended Data Fig. 1; [45, 46]). Identifying this physiological shear scale with Murray’s optimal wall shear rate gives

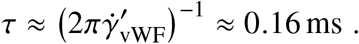

Thus, *τ* is independently constrained by physiological observations rather than fitted to metabolic data. Shear-sensitive endothelial adaptation provides a possible mechanism for maintaining this operating state (Supp. S1; [46, 47]).

### Convection and optimal oxygen utilization

Oxygen-binding pigments such as hemoglobin or hemocyanin increase the oxygen available for transport. We write the inlet oxygen concentration available for extraction as 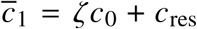, where *c*_0_ is the saturation concentration of physically dissolved oxygen in blood plasma [48], *ζ* the dimensionless relative oxygen capacity of blood, and *c*_res_ ≪ *ζ c*_0_ the residual oxygen concentration. For mammalian blood, *ζ* ≈ 30 [49]. A short capillary leaves extractable oxygen unused; a long capillary exhausts it before the distal tissue is supplied.

The oxygen-utilization optimum therefore satisfies

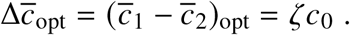

The net convective molar oxygen flow rate through the MH (inflow minus outflow) at this optimum is

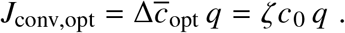

Integral and differential formulations of oxygen conservation are given in Supplements S7 and S8, respectively.

### Diffusion and reaction

Within the MH, the spatially resolved three-dimensional (3D) convection– diffusion–reaction (CDR) solution is reduced to the effective coefficients *D* = Ψ*D* and *k* = Θ*κ*(Extended Data Fig. 5). The coefficients Ψ and Θ are coupled through the same concentration field rather than independently prescribed. Over the physiological MH size range, the self-consistent reduction of the 3D CDR solution gives Ψ ≈ 10^−4^ and Θ ≈ 1, with only weak variation and no systematic dependence on *m* or *N* (Supp. S9; Extended Data Fig. 6). The representative values Ψ = 10^−4^ and Θ = 1 are used below.

Using these effective coefficients *D* and *k*, integration of Fick’s law over the fluid–tissue interface *S* = *πdl* gives the diffusive molar oxygen flow rate of the MH, *J*_diff_ = *c*_0_ *SD*/*δ*, whereas integration of cellular oxygen consumption over the supplied tissue gives the molar oxygen consumption rate within the MH, *R* = *c*_0_*k v*_tiss_.

### Governing equations of the optimized MH

At the MH mesoscale, convection, diffusion, cellular reaction, and geometry are coupled through seven governing equations. Each equation represents a physical transport process or a geometrical relation. Oxygen conservation requires

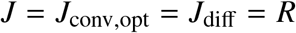

(Supp. S8). Together with Murray’s law and the three geometric relations, these balances form seven equations for *J, q, S, v*_tiss_, *d, l*, and *δ*:

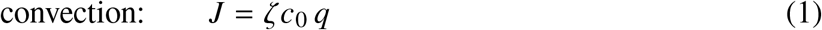

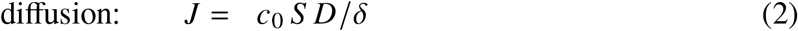

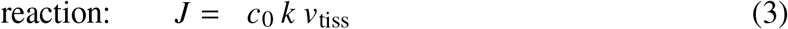

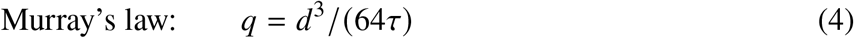

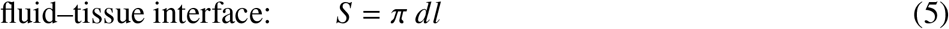

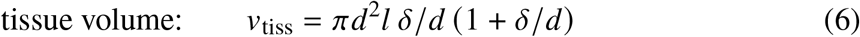

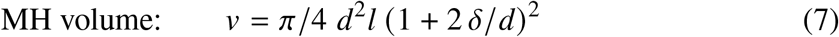

Each equation is physically interpretable and independently testable. Given *v, D, k, τ* and *ζ*, the closed system determines MH geometry (*d, l, δ*) and flow *q* = *q* (*v, D, k, τ, ζ*), with the following limits:

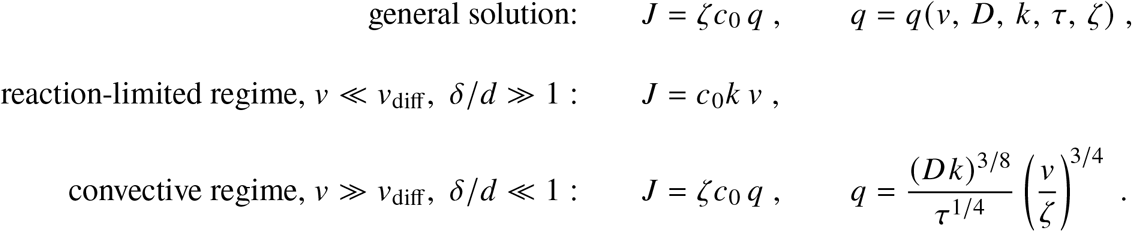

where the MH reaction–diffusion volume is *v*_diff_ := (*D*/*k*)^3/2^.

The first central result is that *one optimized MH already yields q* ∝ *v*^3/4^, continuously approached from the linear regime. Thus, Kleiber’s exponent follows from Krogh’s functional unit, extended by Murray’s principle and optimal oxygen utilization, without requiring complex vascular-network architecture.

Remarkably, dimensional analysis independently recovers both metabolic asymptotes, *m*^1^ and *m*^3/4^, without invoking the MH equations, prescribing vascular-network architecture, or assuming body-mass-invariant topology *N* (Supp. S4). When modular organization into MHs is additionally introduced, the same analysis further constrains the scaling of the individual functional units and their repetition at the organismal scale. Thus, dimensional analysis establishes an independent, architecture-free consistency result, while the MH equations provide the physical transport–reaction mechanism, absolute metabolic rates, and the continuous transition between asymptotes.

## From the MH mesoscale to organismal metabolism

Three distinct operations map the optimized MH to the whole organism. *Kinematic repetition* with topology *N* gives organism volume and basal flow, *V* = *Nv* and *Q*_0_ = *N q. Dynamic conversion* with mass density *ϱ* and metabolic energy density *e* gives body mass and basal metabolic power, *m* = *ϱV* and *Ė*_0_ = *ζ eQ*_0_. Finally, *physiological deployment a* := *Q*/*Q*_0_ describes operation away from the optimized basal state, *Q* = *aQ*_0_ and *Ė*= *aĖ*_0_. The distinction matters biologically: *N* specifies how many functional domains supply the organism, *ϱ* and *e* introduce mass and energy, and *a* describes how the organism uses an already established transport system. Neither *a* nor the deployment state enters MH optimization or the similarity argument for *N*.

The energy scale follows from oxygen utilization. Oxidation of one mole of glucose consumes *v* = 6 moles of oxygen and releases |Δ*H*^◦^| = 2850 kJ/mol [50]. With cellular efficiency *η* ≈ 0.4 [51, 52], the released heat per mole of oxygen is (1 − *η*) |Δ*H*^◦^|/*v*. Thus *e* = *c*_0_(1 − *η*) |Δ*H*^◦^|/*v* ≈ 47.3 kJ/m^3^, where *c*_0_ ≈ 0.17 mol/m^3^ denotes the characteristic saturation concentration of dissolved oxygen in blood plasma and tissue ([48]; Extended Data Table 1). The metabolic power of one MH is *eJ*/*c*_0_.

We hypothesize that physiological deployment is bounded by 0 < *a* ≤ 1 in ectotherms, whereas endotherms operate at *a* = 1 under basal conditions and may reach *a* > 1 under increased metabolic demand. In endotherms, thermoregulation provides an additional functional separation between metabolic demand and body temperature by adjusting cutaneous blood flow and heat transfer. This decoupling allows metabolic power to increase without a proportional rise in body temperature and thereby increases physiological robustness to changing performance demands.

### General MH model and its asymptotes

Repetition, conversion to metabolic power and physiological deployment give the general organism-level MH model:

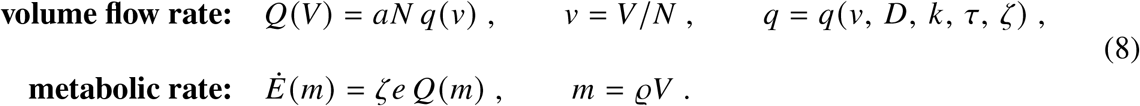

Equation 8 predicts *Q* and Ė in physical units without metabolic-rate normalization; numerical precision remains at the order-of-magnitude level of the coarse-grained model.

With the analytical optimization conditions already incorporated, the resulting nonlinear system is solved numerically (Supp. S10). Its analytical limits follow from the geometric factors 1 + *δ*/*d* and 1 + 2*δ*/*d* in Eqs. 6 and 7; the general prediction and Kleiber asymptote are compared in Fig. 1.

For *m* ≪ *m*_diff_, diffusion alone gives the linear small-organism asymptote *Ė*= *Ė*_0_ = *e κm*/*ϱ*. In this asymptote, *d*/*δ* → 0, physically realized by *d* → 0, so that the tubular convective system vanishes. Accordingly, topology drops out of the leading-order metabolic balance, whereas *N* = 1 marks the first finite tubular supply domain. The linear metabolism of protists is therefore contained as a limiting case of the general MH theory.

For *m*/*m*_diff_ → ∞, the general MH model approaches its Kleiber asymptote, for which

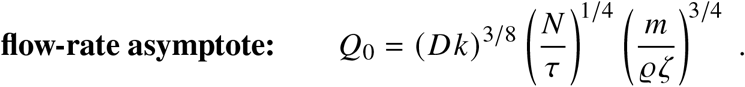

The corresponding Kleiber asymptote for basal metabolic rate is

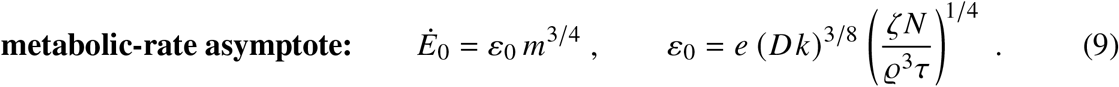

### Evolutionary shifts in metabolic levels

A striking consequence of the MH theory is that functional topology *N* determines the absolute metabolic level. Evolutionary innovation from a single tubular supply domain in planarians (*N* = 1) to repeated MHs in vascularized ectotherms (*N* ≈ 10^5^) and endotherms (*N* ≈ 5 × 10^8^) thus establishes distinct metabolic levels and increases mass-specific metabolic power at a given body mass (Fig. 1). The independently inferred endotherm–ectotherm topology ratio, *N*_endo_/*N*_ecto_ ~ 10^4^ (Supp. S5, Eq. ES29), gives

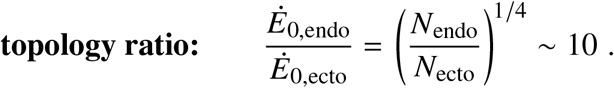

Thus, increasing functional modularity provides an evolutionary route to higher mass-specific metabolic power.

## Heart rate, heartbeats, and lifespan

For the heart-rate and lifespan comparisons, we consider again basal endothermic operation, *a* = 1. Blood volume flow rate is the product of heart rate and stroke volume, *Q*_0_ = *f*_0_ *V*_stroke_ = *f*_0_ *V*/A (Fig. 2). In the Kleiber asymptote of the general MH model, Equation 8 therefore gives the heart-rate asymptote

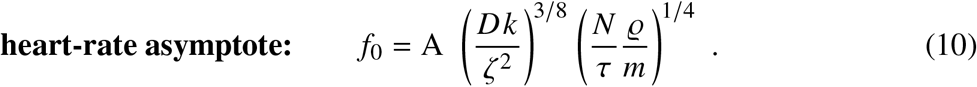

Because A and *N* are group-specific constants, Equation 10 gives *f*_0_ ∝ *m*^−1/4^. Physiological deployment gives *f* = *a f*_0_ without changing the mass exponent, with *a* = 1 at basal operation.

Lifespan requires an additional empirical closure, supplied by the lifetime-heartbeat relation. The characteristic MH turnover time is *t*_MH_ = *v*/*q*. It is the timescale associated with the blood volume flow rate through the complete supplied MH volume, rather than the capillary transit time *v*_cap_/*q*. The dimensionless number Ω := *T*/*t*_MH_ = *Tq*/*v* counts these MH turnover cycles during lifespan *T*. Published lifetime-heartbeat data determine the magnitude of Ω for the endothermic similarity group. Using *Q*_0_ = *N q, V* = *Nv*, and *Q*_0_ = *f*_0_ *V*/A gives

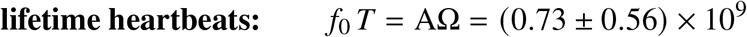

(Extended Data Table 1; [43, 44, 53]). The corresponding lifespan asymptote is therefore

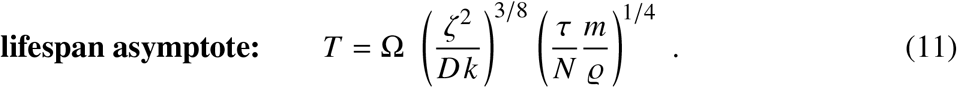

Extrinsic mortality contributes to the scatter of observed lifespans but does not alter the asymptotic prediction *T* ∝ *m*^1/4^.

### Parameter uncertainty

We propagated uncertainty in the independently specified parameters by solving the MH model for 5 × 10^5^ Monte Carlo samples drawn from the distributions in Extended Data Table 1 (Supp. S6; Extended Data Fig. 7). Figures 1–3 show predictive means and central 95 % parameter-uncertainty intervals. Thus, the general MH model (Eq. 8) is evaluated over an ensemble of physiological and structural parameter values sampled from their uncertainty distributions, using standard Monte Carlo uncertainty propagation [54].

## Comparison with observations

Statistical analyses show that metabolic scaling exponents are not uniquely determined by observations [6, 12]. In contrast, MH derives absolute metabolic rates from physical principles; metabolic-rate data are used only for subsequent validation, not parameter fitting. Parameter uncertainty is propagated by Monte Carlo simulation, whereas joint inference of parameter and model uncertainty, as in Bayesian analysis, is beyond the present scope. Importantly, the 3/4 exponent is an asymptotic consequence of the model structure, independent of parameter uncertainty, provided that the constitutive parameters and topology remain body-mass invariant within each metabolic similarity group.

Figure 1 compares these predictions with observations across 18 orders of magnitude in body mass. The general MH model reproduces the linear regime, the continuous transition to convective supply, the group-specific Kleiber asymptotes, and their absolute metabolic levels, with a pooled logarithmic coefficient of determination of *R*^2^ = 97.7 %. Within individual metabolic similarity groups, the logarithmic coefficients of determination are 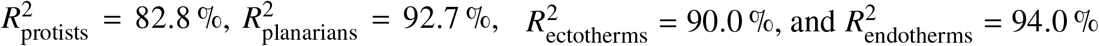.

For endotherms, the general solution reaches the Kleiber asymptote only at large body mass (Fig. 1b). Below 1 kg, the general MH model (Eq. 8) captures the systematic departure from a pure 3/4-power law because the asymptotic condition *δ*/*d* ≪ 1 is not yet fulfilled.

Figure 3 compares predicted and observed heart rates and lifespans in endotherms. Heart rate is predicted without calibration and gives 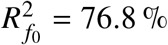. Lifespan uses the empirical lifetime-heartbeat range to determine Ω and gives 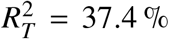; extrinsic mortality remains outside the metabolic model.

**Figure 3:**
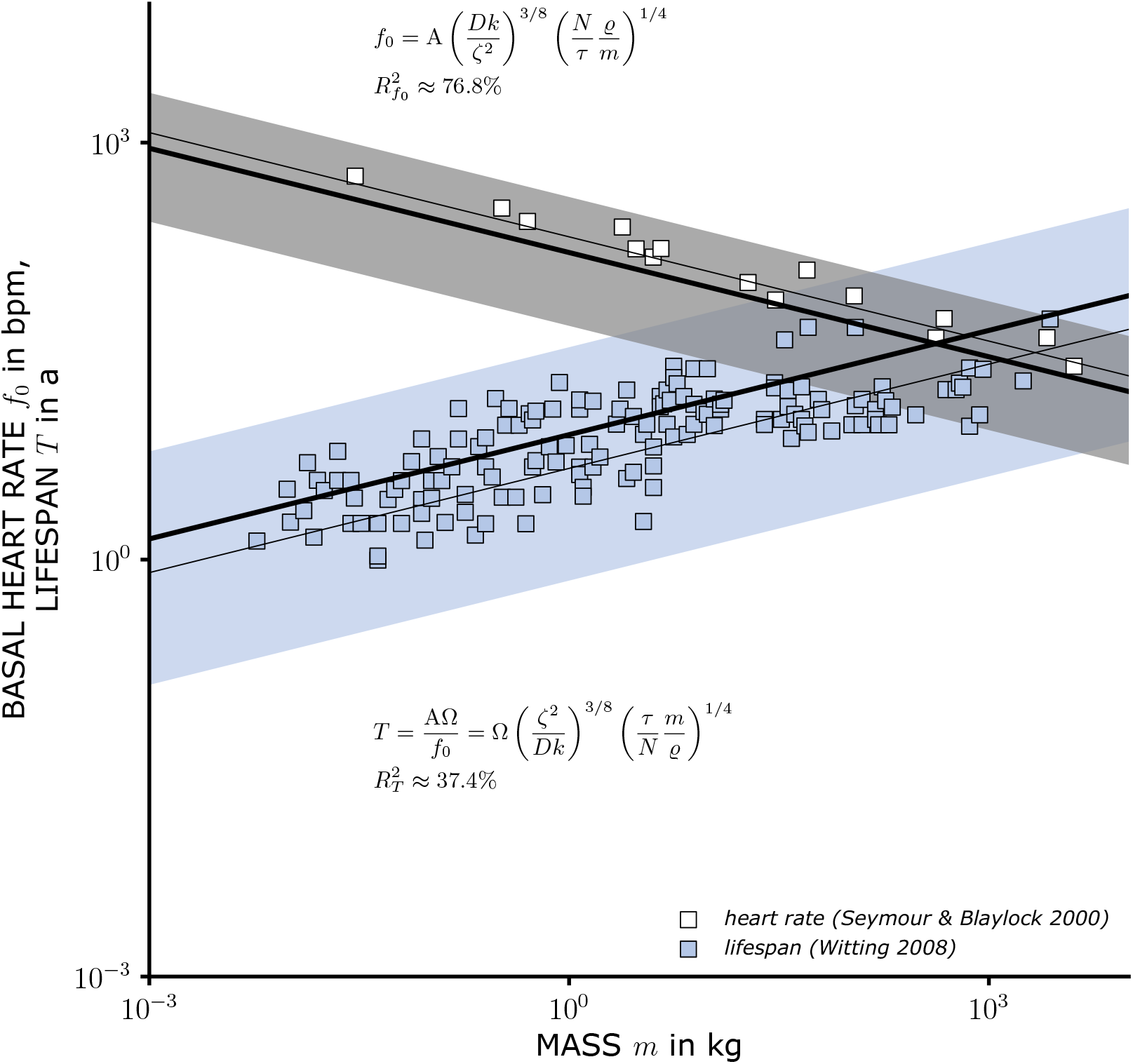
Heart rate and lifespan of endothermic animals. Thick solid lines (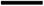): predictive means; shaded regions: central 95 % parameter-uncertainty intervals; thin solid lines (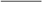): deterministic predictions. The curves represent the heart-rate and lifespan Kleiber asymptotes (Eqs. 10 and 11); the absolute lifespan scale additionally uses the empirical lifetime-heartbeat relation. Symbols denote observations from [43, 44]; 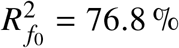 and 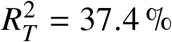.

## The MH theory: biological organization, physical constraints, and evolution

### From relative scaling to absolute prediction

At the MH mesoscale, Murray’s principle balances vascular material costs against viscous dissipation, while optimal oxygen utilization provides an additional constraint. Together, these constraints determine the optimized Krogh unit, from which Kleiber’s asymptote emerges without assuming complex vascular-network architecture. At the organismal macroscale, respiratory optimization matches oxygen supply to tissue demand (Supp. S2). This optimization-based representation of evolutionary processes couples the cellular microscale, MH mesoscale, and organismal macroscale, yielding absolute metabolic predictions consistent with cellular physiology, blood rheology, and Murray’s law. It therefore differs fundamentally from the West–Brown–Enquist (WBE) framework, which derives metabolic scaling exponents from eight assumptions specifying an idealized organism-level vascular-network architecture [3–5, 8–11, 13].

This methodological approach can be visualized in an evolutionary design space (Fig. 4). This space is bounded by (1) available resources, (2) required functions, (3) physicochemical laws, and (4) the repertoire of biological solutions. Resource scarcity and changing functional demands restrict the accessible region. Within this region, constrained optimization represents how selection and adaptation may favor functional transport structures, whereas innovation expands the range of feasible solutions.

**Figure 4:**
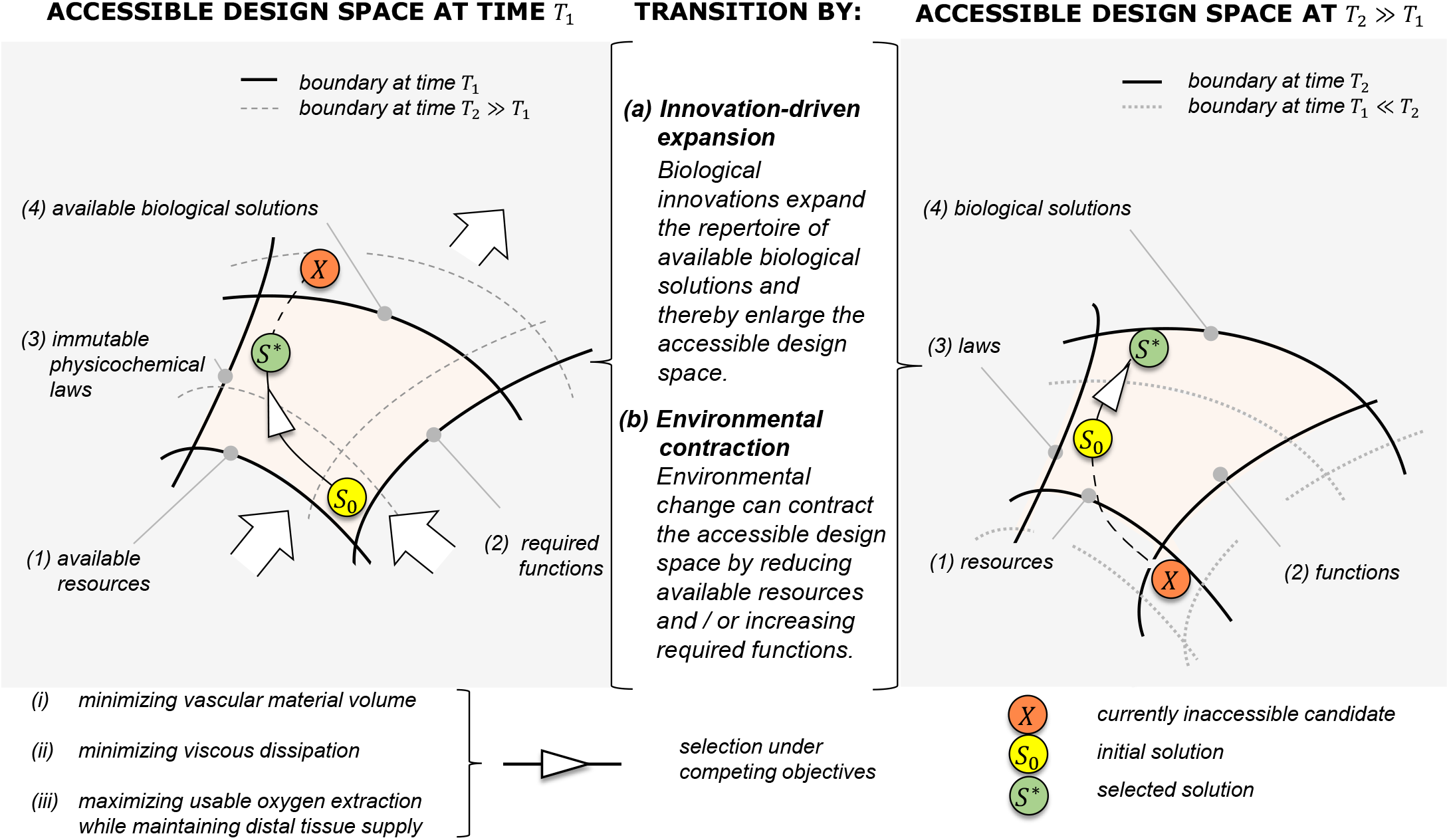
Transition between accessible evolutionary design spaces. Accessible solutions are bounded by available resources, required functions, physicochemical laws, and the repertoire of biological solutions. Selection acts within the accessible space, whereas innovation changes its boundary. *S*_0_ denotes an initial solution, *S*^*\**^ the selected solution, and *X* a solution inaccessible at *T*_1_ but potentially accessible at *T*_2_ ≫ *T*_1_.

### Overcoming the diffusion limit on growth

At the organismal scale, Da_organism_ ≈ 1 marks the diffusion limit on growth in the absence of convective oxygen distribution. For Da_organism_ > 1, molecular diffusion alone can no longer supply the organism. Tubular convection overcomes this limit by distributing oxygen to mesoscale transport domains. For each optimized MH, the mesoscale Damköhler number is Da_MH_ := *k δ*^2^/*D* = (1 + *δ*/*d*)^−1^ < 1. It approaches unity only in the Kleiber asymptote (Supp. S9), where oxygen is supplied *just in time* to meet cellular demand. Thus, tubular convection allows Da_organism_ > 1 and Da_MH_ < 1 to coexist: organismal growth exceeds the diffusion limit while local oxygen supply remains matched to cellular reaction.

### Evolutionary perspective on the MH framework

Metabolic-scaling theory has been framed as a contrast between *“Newtonian”* approaches based on physical constraints and *“Darwinian”* approaches emphasizing adaptive and evolutionary variation [21]. The MH framework connects these perspectives by representing possible outcomes of evolutionary selection and adaptation through constrained model-based optimization at two scales (Fig. 4). At the mesoscale, optimization determines the geometry and operating state of the MH as a functional oxygen-supply unit, yielding metabolic scaling. At the organismal macroscale, optimization matches respiratory oxygen supply to tissue demand, yielding respiratory symmorphosis and linking respiratory surface area to the aggregate microvascular exchange surface (Supp. S2; Extended Data Fig. 2).

The transition beyond diffusion provides a first example of innovation within the MH framework (Fig. 2). Directed tubular geometry introduces convection and thereby opens a region of design space in which growth beyond the organism-scale diffusion limit becomes possible. Planarians represent a single-domain realization, *N* = 1; repetition of such domains in vascularized ectotherms and endotherms constitutes a subsequent organizational innovation, *N* ≫ 1, establishing new feasible metabolic states and distinct metabolic similarity groups.

A developmental implication follows. Gastrulation establishes internal epithelial surfaces and a primitive gut, potentially enabling directed exchange beyond diffusion across an external body surface. The late-Ediacaran bilaterians described by Li *et al*., including numerous elongate worm-like fossils with a terminal mouth and an eversible anterior structure, provide a suggestive morphological context [55]. They are not evidence of an optimized MH: internal oxygen convection and transport–reaction coupling cannot be inferred from these fossils. Rather, they motivate a testable evolutionary question: *when did internal tubular organization begin to function as a single oxygen-supply domain, and when did such domains become repeated?*

Oxygen-binding pigments provide a second example of innovation within the MH framework (Fig. 2). By increasing oxygen-carrying capacity, they enlarge the set of metabolically feasible states and permit high metabolic demand without correspondingly extreme circulatory flow, frequency, or size. At fixed metabolic power and stroke volume,

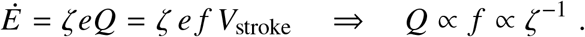

Without hemoglobin’s approximately 30-fold enhancement of oxygen-carrying capacity, maintaining the same metabolic rate would therefore require a 30-fold increase in both volume flow rate and heart rate.

More broadly, the MH framework offers a physical perspective on major evolutionary innovations. The emergence of internal exchange surfaces, oxygen-binding metal complexes, and the repetition of functional supply units can be understood as expansions of the accessible metabolic design space. Each innovation relaxes transport or resource constraints, enabling higher metabolic power at a given organismal mass or reducing the structural and operational costs of sustaining a prescribed metabolic rate. Thus, the MH framework not only predicts metabolic scaling and absolute rates but also provides a physical basis for understanding how evolutionary innovation expands the metabolic possibilities of life.

## Methods

The MH model is defined by Equations 1–7. The nonlinear system was solved either by Newton– Raphson iteration or by the spreadsheet procedure described in Supplement S10. The integral and differential oxygen balances are derived in Supplements S7 and S8, respectively; Supplement S8 also gives the three-dimensional CDR solution (Extended Data Fig. 5), which is mapped to the effective coefficients *D* and *k* in Supplement S9 (Extended Data Fig. 6).

For the numerical predictions, the representative CDR values Ψ = 10^−4^ and Θ = 1 are fixed. Topology is centered at the nominal group values *N*_endo_ = 5 × 10^8^ and *N*_ecto_ = 10^5^, with topology uncertainty propagated according to Extended Data Table 1. The independent dimensional analysis is given in Supplement S4. Parameter values and probability distributions were specified from independent literature sources (Extended Data Table 1), and uncertainty was propagated through 5 × 10^5^ Monte Carlo realizations (Supp. S6).

The coefficients of determination were evaluated on the log scale. For the predicted values, the logarithm of the mean model response from the Monte Carlo simulation was used. For the metabolic rate, the model was evaluated at discrete mass points that do not necessarily coincide with the observed masses; the corresponding predictions were obtained by linear interpolation in log–log space.

The geometric and physiological observations used for inference and testing are listed in Table 1 and analyzed in Supplement S5.

## Data availability

The complete reproducibility package for this study is publicly available on Zenodo in accordance with the FAIR principles (Findable, Accessible, Interoperable, and Reusable): https://doi.org/10.5281/zenodo.23472906. The package includes

i. all digitized experimental data underlying the empirical validation;
ii. all parameter files, including the self-consistent 3D CDR solutions used to determine Ψ and Θ;
iii. the complete Python implementations used for the numerical calculations, including the Monte Carlo uncertainty analysis and Newton–Raphson solution procedures;
iv. all scripts and notebooks required to generate the figures; and
v. a spreadsheet implementation of the MH theory, enabling its application without Python or dedicated numerical software.

Thus, all data, parameters, and computational procedures required to reproduce the quantitative results and figures reported in this study are publicly available.

## Code availability

All code is included in the same Zenodo reproducibility package: https://doi.org/10.5281/zenodo.23472906.

## Use of generative AI

Generative AI was used solely for editorial and language revision. All scientific content and conclusions were developed and verified by the authors.

## Acknowledgements

We thank Technische Universität Darmstadt for fostering an academic environment of trust, intellectual openness, and freedom in which this work could develop, and Katharina Henn and Kevin Logan for their careful comments on the manuscript. P.F.P. thanks his beloved wife Stephanie and his children Emma and Jakob for their patience and love, and for giving him the space and freedom to think.

## Author contributions

P.F.P.: conceptualization, theory, methodology, analysis, visualization, and writing. T.M.: uncertainty quantification, Monte Carlo simulation, numerical predictions, and writing of the uncertainty-analysis sections. Both authors reviewed and approved the manuscript.

## Additional information

Supplementary Information accompanies this paper. Correspondence and requests for materials should be addressed to P.F.P.

## Extended Data

**Extended Data Figure 1:**
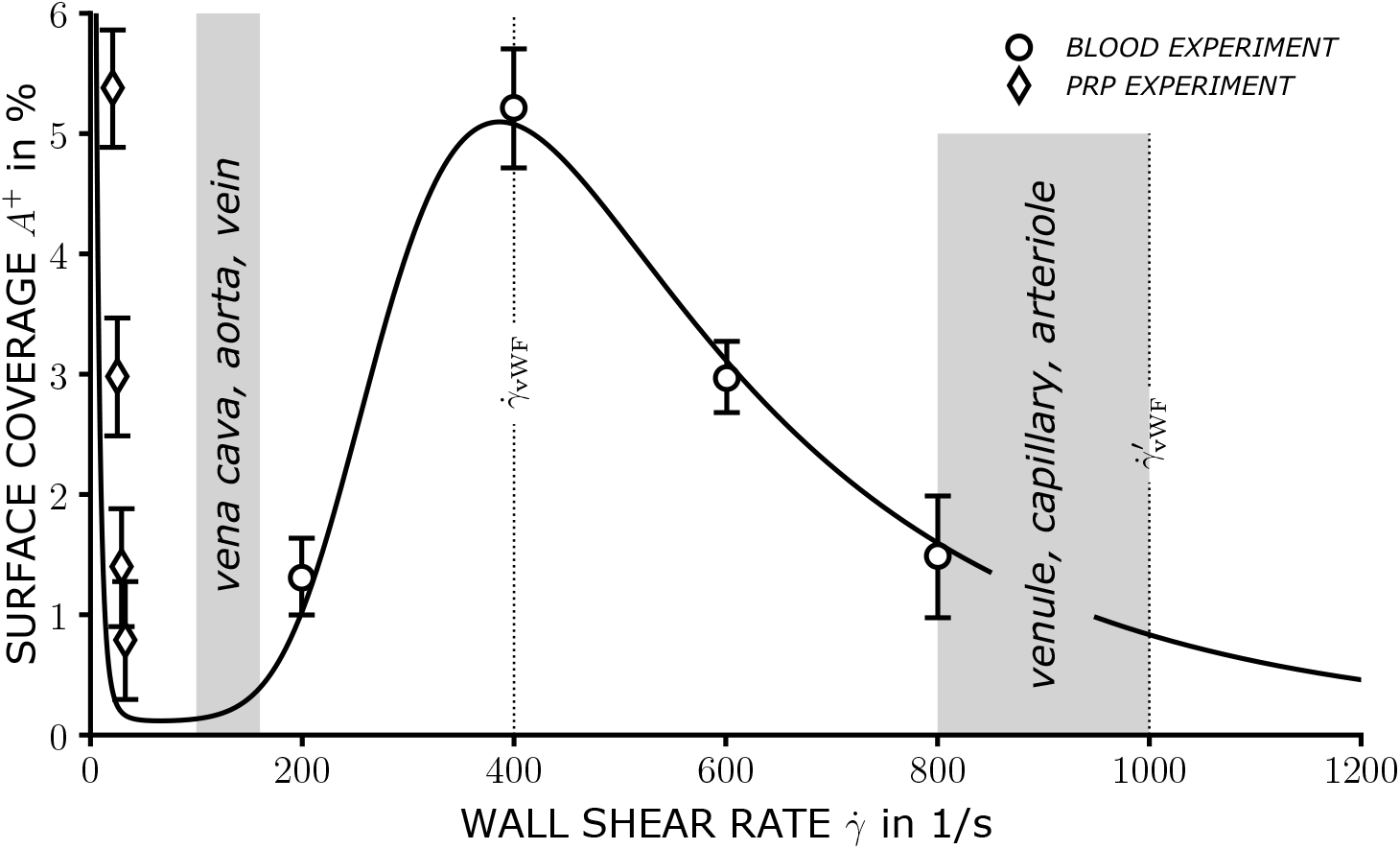
Shear-dependent platelet adhesion and the Murray timescale *τ*. Symbols show platelet surface coverage versus wall shear rate; the solid line is the adhesion model [60]. The high-shear, low-adhesion branch is characterized by 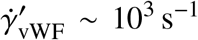, giving 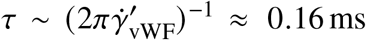 (Supp. S1, Eqs. ES1–ES2). Shaded regions indicate representative vessel-class ranges.

**Extended Data Figure 2:**
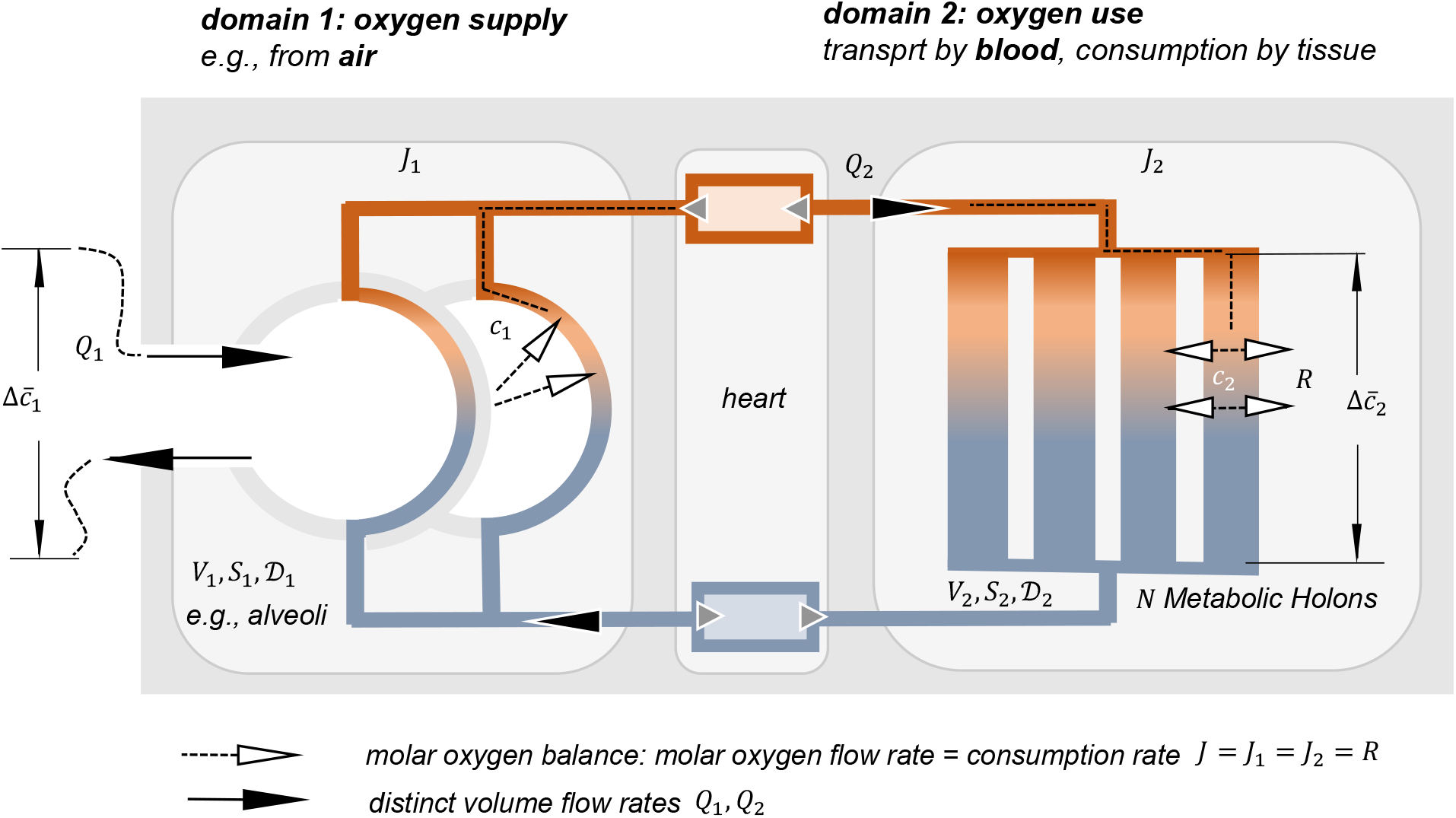
Respiratory symmorphosis through organism-scale transport optimization. Two coupled domains—oxygen supply (domain 1) and oxygen use (domain 2)—are characterized by volume flow rates *Q*_*i*_, volumes *V*_*i*_, diffusion coefficients *D*_*i*_, and exchange surfaces *S*_*i*_. Circulation connects respiratory uptake to the *N* Metabolic Holons, where oxygen is consumed. At steady state, the molar oxygen flow is conserved, *J* = *J*_1_ = *J*_2_ = *R*. Constrained optimization yields contact-time matching (A); a leading-order transport approximation and oxygen conservation then give the further matching relations B and C: **A**, contact times *t*_1_ = *t*_2_ = *ε*_RS_ = *B t*_MH_ = *V*_blood_/(*V*_stroke_ *f*_0_) (Eq. ES8); **B**, exchange surfaces 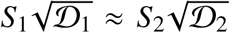 (Eq. ES12) and **C**, oxygen-content matching 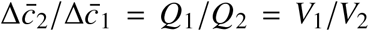 (Eq. ES13). Together, these matching relations express respiratory symmorphosis (Supp. S2).

**Extended Data Figure 3:**
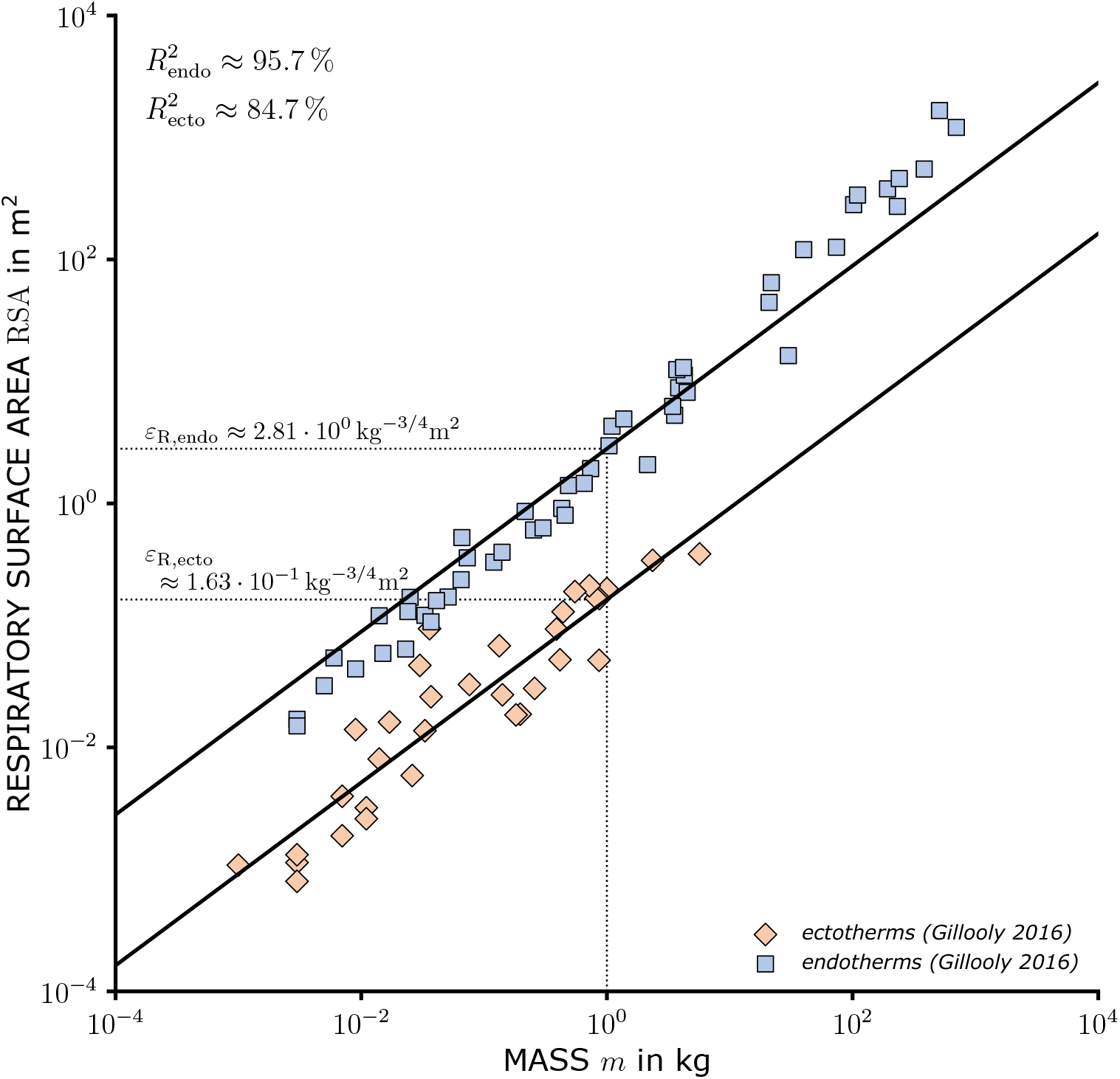
Respiratory surface area (RSA) across body mass in endotherms and ectotherms. Symbols show RSA data from Gillooly *et al*. [41]. Solid lines are fixed-slope regressions of the MH relation RSA = *ε*_R_*m*^3/4^ (Eq. ES27); only the group-specific prefactors are fitted.

**Extended Data Figure 4:**
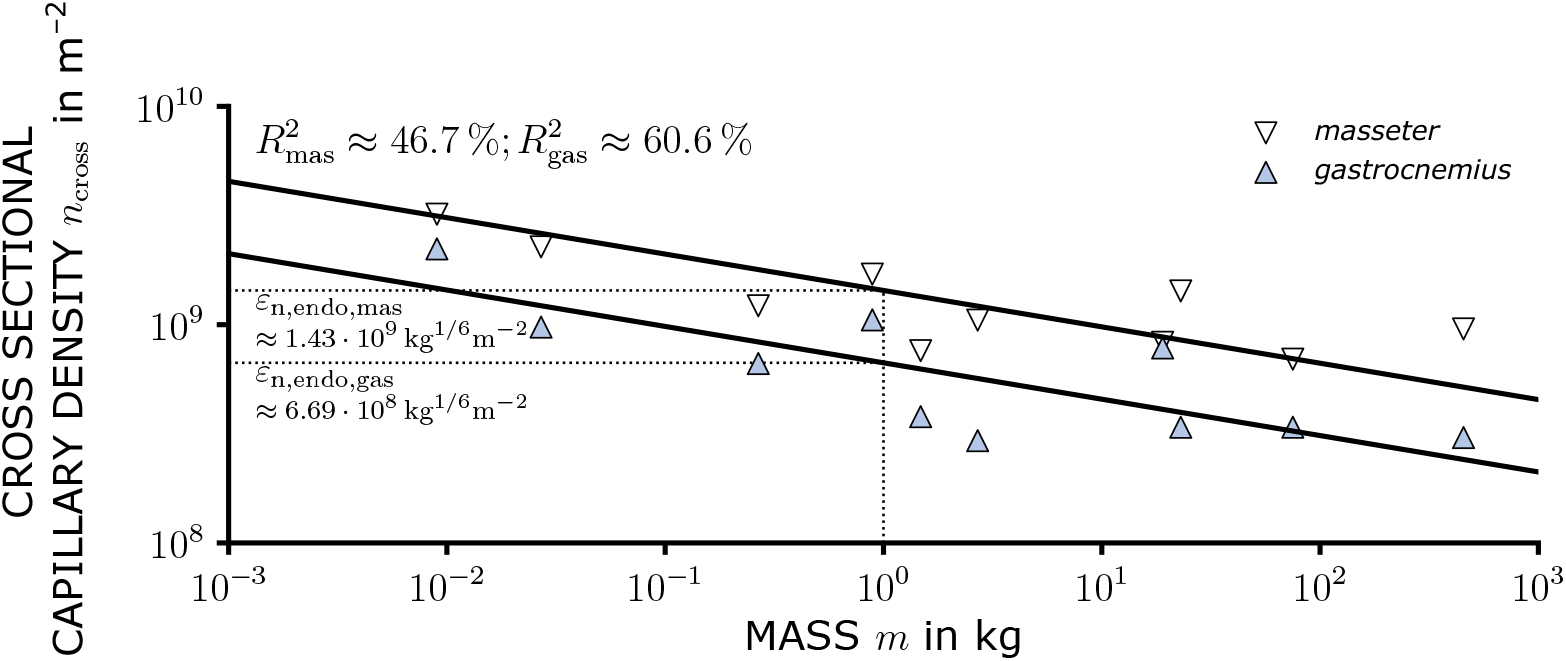
Cross-sectional capillary density across mammals. Symbols show masseter and gastrocnemius data from Schmidt-Nielsen and Pennycuik [25]. Solid fixed-slope regressions test the MH prediction *n*_cross,vitro_ ∝ *m*^−1/6^ (Supp. S3, Eq. ES22); only tissue-specific prefactors are fitted.

**Extended Data Figure 5:**
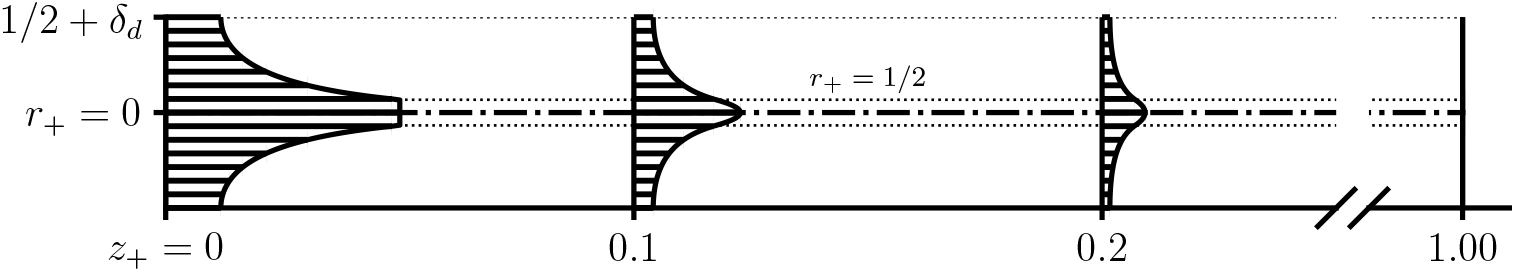
Analytically calculated oxygen-concentration field in a representative MH. The 3D convection–diffusion–reaction solution shows axial oxygen depletion and radial diffusion into reacting tissue (Supp. S8, Eqs. ES36–ES47). It underlies the effective coefficients *D* = Ψ*D* and *k* = Θ*κ* (Supp. S9, Eqs. ES48–ES49). Axial and radial dimensions are enlarged for visualization.

**Extended Data Figure 6:**
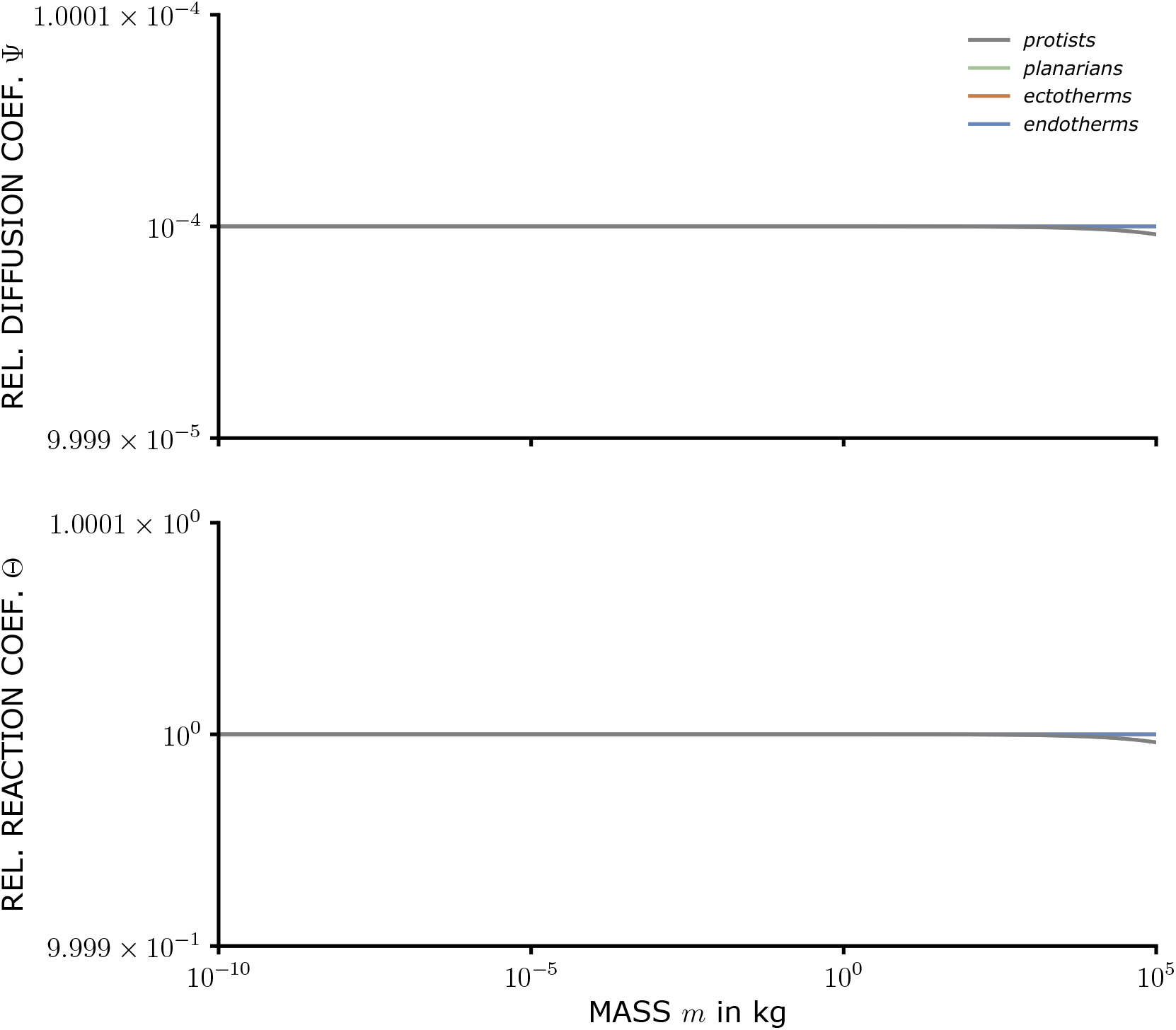
Relative diffusion and reaction coefficients from the 3D CDR solution. The coefficients Ψ = *D*/*D* and Θ = *k*/*κ* follow from the fixed-point CDR solution (Supp. S9, Eqs. ES48 and ES49). Across the physiological MH range, the converged solution remains close to Ψ = 10^−4^ and Θ = 1.

**Extended Data Figure 7:**
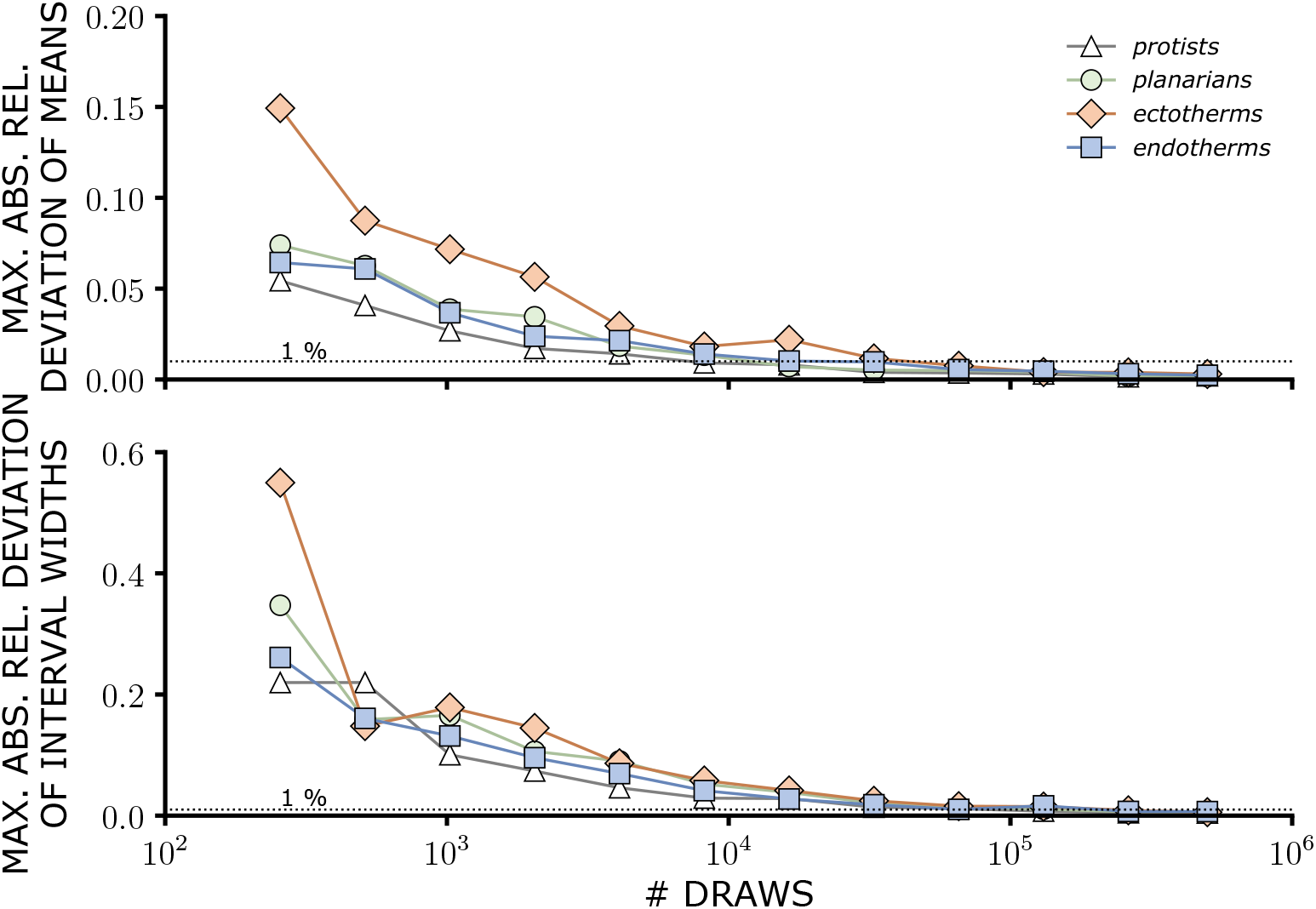
Convergence of the Monte Carlo uncertainty analysis for metabolic rate. Each draw evaluates the general MH model (Eqs. 1–8). Shown are the maximum relative changes in predicted means and 95 % interval widths between successive sample-size doublings, maximized over all considered masses.

**Extended Data Table 1:**
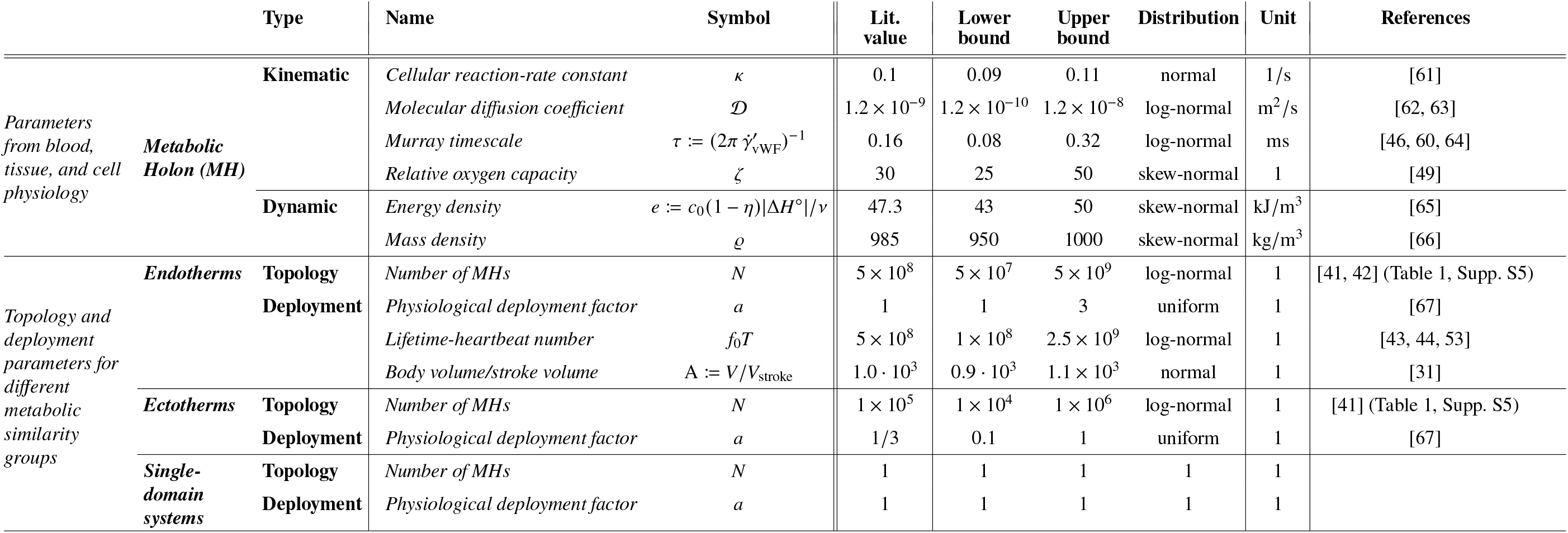
Model parameters of the MH theory. No parameter is fitted to the metabolic-rate or heart-rate observations used for comparison. The core metabolic-rate prediction depends on six independently specified physiological quantities, *κ, D, τ, ζ, e*, and *ϱ*, together with the independently constrained structural quantity *N*. The factor *a* describes physiological deployment away from the basal MH state and is not a parameter of the mesoscale MH optimization; A and the lifetime-heartbeat closure enter only the heart-rate and lifespan extensions, respectively. Numerical predictions use the relative diffusion and reaction coefficients Ψ = 10^−4^ and Θ = 1, so that *D* = 10^−4^ *D* and *k* = *κ*; their derivation and physical interpretation are given in Supplement S9. The nominal topology values *N*_endo_ = 5×10^8^ and *N*_ecto_ = 10^5^ are representative group values anchored by the RSA-based inference and supported by the complementary evidence in Table 1 and Supplement S5.

## Supplementary Information

*Detailed theory, derivations, validation, and methods*

## S1 Body-size-invariant wall shear, Murray’s law, and physio-logical vascular adaptation

Murray’s *physiological principle of minimum work* combines two competing objectives: minimizing the material or maintenance cost associated with blood volume and minimizing viscous dissipation [16]. For a capillary of length *l*, diameter *d*, cross-sectional area *A* = *πd*^2^/4, and fluid–tissue interface *S* = *πdl*, these objectives are represented by the blood mass *ϱ Al* and the dissipated power *P*_D_. To obtain a dimensionally homogeneous objective function, blood mass is weighted by the Pareto parameter *ε*_MH_, which has the dimension of mass-specific power:

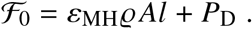

For pressure-driven flow, *U* denotes the cross-sectional mean velocity and the apparent wall shear rate is

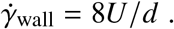

This Poiseuille relation assumes laminar, quasi-stationary flow. Within the MH theory, the corresponding Womersley number scales in the Kleiber asymptote as

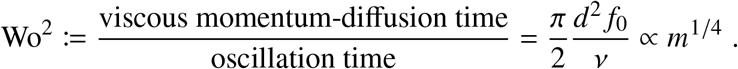

It therefore increases only weakly with body mass; even at the upper mammalian size limit, represented by a large whale, the MH prediction gives Wo^2^ ~ 10^−3^ ≪ 1. Thus, the quasi-stationary Poiseuille description remains valid throughout the physiological mass range considered here. The dissipated power can therefore be written as

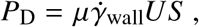

where *µ* denotes the apparent dynamic blood viscosity, i.e., the effective viscosity obtained by representing the local stress–shear-rate relation in Newtonian form although blood itself is non-Newtonian.

In Murray’s optimization, *d* and *U* are the design variables, whereas *l* and the volume flow rate *q* = *U A* are prescribed. The constrained problem is

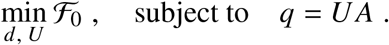

Multiplication of the objective function *ℱ*_0_ by 4/(*ε*_MH_ *ϱ πl*) gives the equivalent objective function

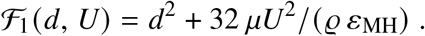

Geometrically, ℱ_1_(*d, U*) defines the objective landscape, whereas the constraint

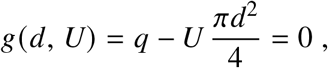

defines the admissible path through this landscape. Introducing a Lagrange multiplier *λ* incorporates the prescribed-flow constraint into the objective and recasts the constrained optimization

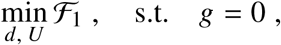

as a stationarity problem in the augmented variables. The constrained optimum is therefore not characterized by ∇ℱ_1_ = 0, but by the condition that no admissible direction along the constraint can further decrease the objective. Equivalently, the gradient of the objective must be normal to the constraint manifold,

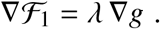

Thus, the Lagrange multiplier identifies the point where the constraint path reaches its minimum on the objective landscape.

With the Lagrangian

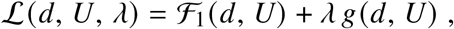

the constrained optimum follows from the stationarity condition ∇ℒ = 0, i.e.,

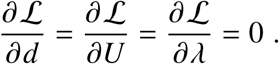

Solving these conditions yields Murray’s law

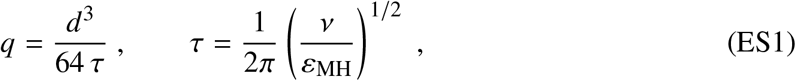

where *v* = *µ*/*ϱ* is the kinematic viscosity.

### Relation to network-based scaling theories

The WBE model, as formalized by Savage *et al*., rests on eight structural and physiological assumptions [13]. Of particular relevance here is the assumption of *“size-invariant terminal units”* [11, 13]: capillary flow rate and dimensions are taken to be independent of body size. This assumption is essential rather than auxiliary because it sets the terminal scale across organisms [13]. By contrast, the mesoscale MH optimization introduces only one additional optimization postulate beyond Murray’s established material–dissipation trade-off: optimal oxygen utilization.

The geometric consequences of “size-invariant terminal units” have been the subject of substantial debate. In particular, their compatibility with space filling and the resulting metabolic scaling was critically questioned by Kozłowski and Konarzewski and subsequently defended by Brown, West, and Enquist [10, 11]. Independent of this debate, Dawson reported systematic variation of mammalian capillary dimensions with body mass, supported by available measurements and inconsistent with strict size invariance [15].

Murray’s optimization provides a different starting point because capillary diameter *d* and flow *q* are intrinsically coupled. Equation ES1 gives *q* ∝ *d*^3^ and, at vascular bifurcations, the corresponding cubic relation between parent and daughter vessel radii [16]. Khamassi *et al*. imposed this cubic relation in the optimization of vascular bifurcations and obtained branching geometries within the range of experimental observations [46]. Thus, flow and vascular geometry cannot be specified independently at the mesoscale.

Where WBE takes terminal capillary characteristics as size-invariant boundary conditions, the MH formulation determines flow and oxygen-supply geometry jointly from convection, diffusion, reaction, and Murray optimization. Capillary dimensions therefore emerge from the mesoscale transport problem and vary with the volume supplied, consistent with reported mammalian capillary scaling [15].

### Microscale determination of the Murray timescale

Mathematically, *ε*_MH_ weights the two objectives and parameterizes the Pareto-optimal geometries. Physically, together with *v*, it defines a Kolmogorov-type timescale familiar from turbulence theory,

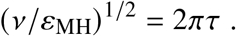

The optimization therefore determines the cubic flow–diameter relation but does not identify the physiological process selecting the numerical value of *τ*.

An independent wall-shear-rate scale is provided by the shear-sensitive interaction of von Willebrand factor (vWF) and platelets. Fluid shear regulates vWF conformation and function [45]. In the microfluidic flow-chamber experiments and adhesion model of Khamassi [60], platelet surface coverage (Extended Data Fig. 1) exhibits a pronounced adhesion range around 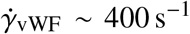, where hydrodynamic transport and receptor–ligand binding are favorably matched.

At higher wall shear rates, platelet surface coverage decreases again. The range approximately 800–1200 s^−1^ defines a high-shear, low-adhesion branch of the same vWF-mediated response [60]. We denote its characteristic shear rate by

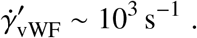

Thus, 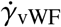 characterizes the adhesion-active part of the response, whereas 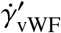 characterizes its high-shear, low-adhesion branch.

Murray’s law relates this independently observed wall-shear-rate scale directly to the Pareto parameter. For fully developed flow in a circular vessel,

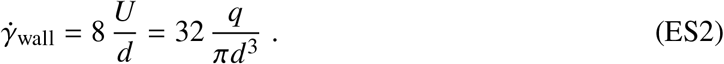

Insertion of Murray’s law, *q* = *d*^3^/(64*τ*), gives

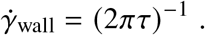

Thus, once *τ* is selected by a group-specific physiological shear scale, Murray optimization implies a body-size-invariant wall-shear state: changes in capillary diameter are accompanied by *q* ∝ *d*^3^, leaving *U*/*d* invariant. Identifying the selected Murray wall-shear rate with the characteristic high-shear, low-adhesion scale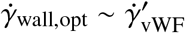, yields

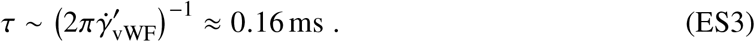

The factor 2*π* has no independent physiological meaning; it enters only through the definition of *τ* chosen to simplify Murray’s relation. The shear-dependent vWF–platelet response therefore provides an independently observed physiological wall-shear-rate scale that closes the otherwise undetermined Pareto parameter.

### Shear-sensitive vascular adaptation

The vWF-associated shear scale specifies a physiological vascular operating state but need not itself be the molecular controller of vessel growth. Independent evidence provides the feedback mechanism by which such a shear state can be maintained. Vascular endothelial cells sense wall shear and transduce persistent changes in the mechanical stimulus into cellular responses that alter vascular geometry [47, 68]. Consistently, wall shear stress acts as a command variable for vascular geometry in the optimization and adaptation framework of Khamassi *et al*. [46].

This provides a feedback interpretation of Murray’s optimum. When wall shear deviates from the selected operating state, shear-sensitive endothelial responses can modify vessel diameter; the resulting change in geometry in turn changes wall shear. The feedback can therefore drive the vessel toward

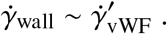

Murray’s relation may thus be interpreted as the stationary state of a physiological feedback loop: the independently observed shear scale specifies the operating point, endothelial mechanosensing detects departures from it, and vascular remodeling adjusts geometry toward that state. The vWF response provides the physiological shear scale, whereas endothelial mechanotransduction provides the adaptive feedback by which it can be maintained.

### Generality of the wall-shear timescale beyond vascular systems

The success of the MH theory in predicting the metabolic rate of planarians is striking, cf. Figure 1. It suggests that *τ* should be interpreted more generally as a physiologically selected convective timescale associated with wall shear rather than exclusively as a parameter of vertebrate blood vessels. In vascular MHs, its numerical value is constrained by the vWF-associated wall-shear-rate scale and therefore reflects the shear-responsive properties of blood and the vessel wall. In simpler tubular transport systems without a vascular network, the corresponding physiological mechanism may differ.

Any internal flow bounded by living tissue generates a characteristic wall-shear rate 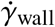. Its physiological operating range need not be arbitrary: processes at the fluid–tissue interface, including solute transport, adhesion, mechanosensing, and tissue remodeling, can provide corresponding biological constraints. Shear-sensitive growth or adaptation may therefore select a preferred operating range also in simpler tubular organisms such as planarians, although the specific physiological process selecting *τ* in such systems remains unknown.

The absence of an independently established nonvascular physiological scale for *τ* does not strongly affect the present metabolic predictions because they depend only weakly on its precise value. In the Kleiber asymptote, the theory gives

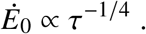

Even substantial variation in the characteristic wall-shear timescale therefore produces a comparatively small change in predicted metabolic rate. The physiological realization of *τ* in nonvascular organisms may differ from the vascular vWF-based constraint while remaining compatible with the same leading-order MH transport structure. Its nonvascular physiological determination therefore remains an open and experimentally testable question.

## S2 Organism-level optimization predicts respiratory symmorphosis

Symmorphosis describes the quantitative matching of structural capacity to functional demand according to an *“economy of design”* [69]. In analogy to Murray’s minimum-work principle [16], we formulate this idea as a constrained optimization of organismal oxygen supply and use (Extended Data Fig. 2). The optimized state is interpreted as a possible outcome of evolutionary selection and adaptation, rather than as an explicit model of evolutionary dynamics.

Domain 1 supplies oxygen from air or water: *V*_1_, *Q*_1_, *S*_1_, and *D*_1_ denote its effective respiratory exchange volume, ventilatory flow, exchange surface, and oxygen diffusivity. Domain 2 represents distributed oxygen use, connected to the respiratory system by circulation: *V*_2_ = *V*_blood_ and *Q*_2_ denote circulating blood volume and flow rate, while *S*_2_ is the aggregate blood–tissue exchange surface of the repeated MHs and *D*_2_ the corresponding oxygen diffusivity. The organismal oxygen flow is common to both domains, *J*_1_ = *J*_2_ = *J*, whereas *v, q*, and the single-MH oxygen flow refer to one mesoscale unit. This organism-level optimization yields matching relations, not the MH metabolic asymptotes; Relation B below also supplies the RSA constraint on *N* used in Supplement S5.

### S2.1 Mesoscale mass transfer

Physiological oxygen exchange is governed at the mesoscale, where molecular diffusion couples the flowing respiratory or circulatory medium to the exchange surface. On this scale, the corresponding air, water, and blood flows are laminar and quasi-stationary.

For the vascular case, Supplement S1 shows from the MH asymptotes that the Womersley number, defined by Wo^2^ := (*π*/2)*d*^2^ *f*_0_/*v*, scales as Wo ∝ *m*^1/8^ and remains small even at the upper mammalian size limit (Wo^2^ ≈ 0.0006 for a large whale). Thus, unsteady inertia is negligible compared with viscous momentum diffusion, supporting the quasi-steady approximation for vascular flow.

We first formulate the molecular transport problem independently of its specific physiological realization. Let *S* denote an exchange surface and 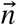 its unit normal. The molecular molar flow through this surface follows directly from Fick’s law, 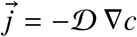, as

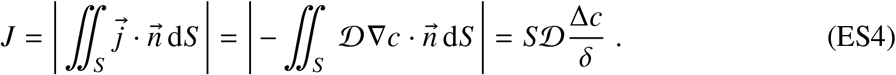

The last equality defines the effective concentration-boundary-layer thickness *δ* for the driving concentration difference Δ*c*. With a characteristic exchange length *L*, the Sherwood number is therefore

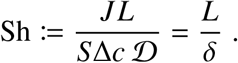

### Mass transfer from Sherwood relations

The Sherwood number represents a dimensionless molar oxygen flow rate, whereas the Péclet number represents a dimensionless volume flow rate. We now establish the relation between these two quantities.

The local concentration field depends on how the fluid is renewed and convected along the exchange surface. In respiratory exchange, the hydrodynamic mechanisms may range from penetration-type renewal to nearly uniform plug-flow convection and developing laminar velocity boundary layers. The central question for the present mesoscale theory is whether these distinct transport mechanisms lead to a common leading-order mass-transfer law.

Within the quasi-stationary laminar regime considered here, all three limiting descriptions yield

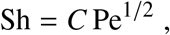

where the dimensionless coefficient *C* accounts for mechanism-specific details, including the Schmidt-number dependence. This common square-root relation provides the basis for the respiratory symmorphosis and matching relations developed below. We now derive it for the three limiting cases.

1. **(i) Transient penetration transport**. Let *y* denote the coordinate in the surface-normal direction. A fluid element exposed to the exchange surface for a finite time obeys the one-dimensional diffusion equation

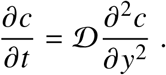 For a sudden change of concentration at the surface of a semi-infinite fluid from *c*_∞_ to *c*_wall_, the classical solution is

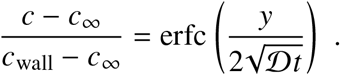

Dimensional analysis already shows that the only length that can be formed from *D* and *t* is 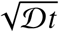. For the error-function profile, the integral penetration depth is defined by

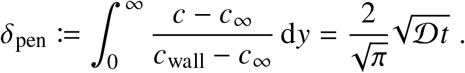

The instantaneous wall flux follows from Fick’s law as

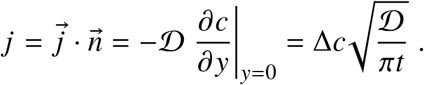

Averaging this flux over the contact time gives

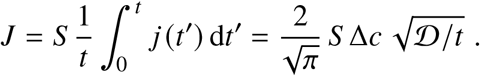

With the Péclet number Pe := *U L*/*D* and the contact or residence time *t* = *L*/*U*, this becomes

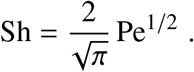
2. **(ii) Plug-flow convection**. Consider now a fluid convected with constant velocity *U* along the exchange surface. With *x* denoting the streamwise coordinate, convection along the surface and molecular diffusion normal to it obey

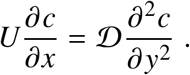 The coordinate transformation *t* = *x*/*U* gives *∂*/*∂x* = (1/*U*) *∂*/*∂t* and transforms this equation exactly into

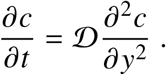

Thus, plug-flow convection and transient penetration transport are mathematically identical under the transformation *t* = *x*/*U*. Consequently, they have the same square-root transfer law,

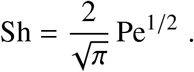
3. **(iii) Laminar Blasius concentration boundary layer**. The plug-flow approximation neglects the no-slip condition at the wall and hence the velocity decrease towards the surface. A laminar boundary layer resolves this velocity field and therefore provides a refined description of continuously flowing fluid along the exchange surface. The classical surface-averaged mass-transfer result is

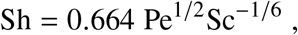

where the Schmidt number Sc := *v*/*D* is the ratio of kinematic viscosity to molecular diffusivity for oxygen in the respective fluid (air, water, or blood plasma). This Sherwood relation follows from the corresponding Nusselt relation by replacing thermal diffusivity with molecular diffusivity and the Nusselt number with the Sherwood number. This analogy is justified by the identical mathematical structure of the corresponding boundary-layer transport equations under equivalent boundary conditions [70, 71].

The three descriptions therefore differ in their local physics but not in their leading mathematical dependence on convective exposure. Penetration transport and plug-flow convection are exactly equivalent under the transformation *t* = *x*/*U*, whereas the Blasius solution resolves the additional velocity-boundary-layer structure. In all three cases, however, the surface-averaged mass transfer reduces to the same square-root law

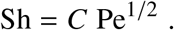

The mechanism-specific information is contained only in the dimensionless prefactor *C*: for penetration transport and plug flow, 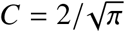; for the laminar Blasius boundary layer, *C* = 0.664 Sc^−1/6^.

Equivalently, the effective transfer thickness is

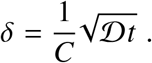

Equivalently, within the coarse-grained, order-of-magnitude MH framework, the effective transfer thickness is expressed in terms of the contact or residence time *t* = *V*/*Q* as 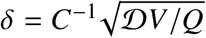.

Thus, irrespective of which of these three limiting descriptions is used, the leading mesoscale dependence is 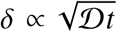 or, equivalently, Sh ∝ Pe^1/2^. It is this shared structure—rather than the particular value of *C*—that generates the respiratory symmorphosis and matching relations derived below. For mammalian airways, geometry-specific physiological mass-transfer models likewise recover the square-root dependence Sh ∝ Pe^1/2^ [72], providing an independent physiological consistency check.

#### Application to respiratory exchange

The common square-root transfer law can now be applied to the oxygen-transport domains considered here: alveolar air or water in gills on the oxygen-supply side, and blood in capillaries on the oxygen-use side. Although the local transport mechanisms may differ, each domain enters the organism-level model through the same reduced relation. Restoring the domain index gives 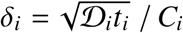, with *t*_*i*_ = *V*_*i*_/*Q*_*i*_ and *i* = 1, 2. Inserting this into Equation ES4 gives

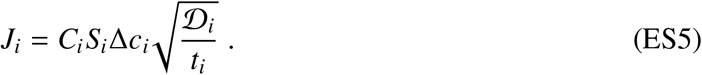

Introducing the volume-specific exchange surface *s*_*i*_ := *S*_*i*_/*V*_*i*_ and using *t*_*i*_ = *V*_*i*_/*Q*_*i*_ gives

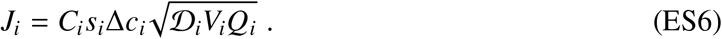

Thus, the local transport physics is condensed into the domain-specific quantities *C*_*i*_, *D*_*i*_, *s*_*i*_, and Δ*c*_*i*_, while organism-level design and operation enter only through the exchange volume *V*_*i*_ and volume flow rate *Q*_*i*_.

### S2.2 Coupling oxygen supply and use: organism-level model-based optimization

At the organismal scale, steady oxygen conservation requires a common molar oxygen flow through both exchange domains, *J*_1_ = *J*_2_ = *J* (Extended Data Fig. 2). For a prescribed organismal oxygen flow *J*, Equation ES6 yields a product constraint *V*_*i*_*Q*_*i*_ = const in each domain: reducing structural exchange volume requires greater operational volume flow, and vice versa. The common molar oxygen flow *J* couples the two domains (Extended Data Fig. 2). The explicit constraints enter the optimization below.

#### Model-based constrained optimization at the organismal macroscale

The core idea of the MH framework is to represent natural selection through model-based constrained optimization. The competing optimization pressures follow Murray’s principle: minimizing structural resource use and operational effort while fulfilling the required function, here oxygen uptake and transport as demanded by the organism’s metabolic needs.

For prescribed organismal oxygen flow *J*, we treat the domain-specific transport parameters *C*_*i*_, *s*_*i*_, Δ*c*_*i*_, and *D*_*i*_ as fixed during the constrained optimization. Equation ES6 then restricts the admissible states of each functional system to

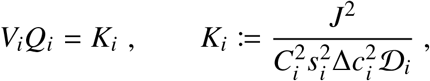

with

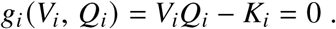

Following the economy-of-design principle underlying symmorphosis and Murray’s *“physiological principle of minimum work”* [16, 69], the competing structural and operational demands are represented by

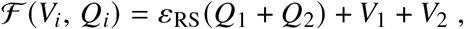

subject to *g*_*i*_ = 0. The first term penalizes operational volume flow, whereas the second penalizes structural exchange volume. The Pareto parameter *ε*_RS_ has dimensions of time and balances these competing demands. The optimum is derived for a prescribed organismal oxygen flow, while physiological parameter variability is treated separately by evaluating the resulting model over an uncertainty ensemble. This separation between optimization and subsequent robustness assessment parallels design-under-uncertainty approaches in engineering, in which performance is assessed over uncertain loads and parameters rather than only at a single nominal point [54]. The Lagrangian is

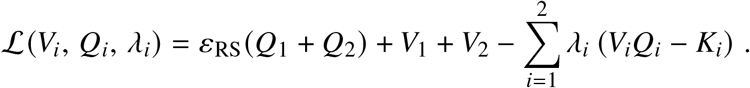

Stationarity gives

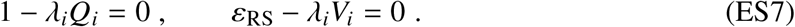

#### Three matching relations of respiratory symmorphosis

The constrained optimization yields three matching relations between organism-level oxygen supply and use. Relation A follows directly from stationarity (Eq. ES7), Relation B additionally uses a leading-order approximation for local mass transfer, and Relation C follows from Relation A and oxygen conservation.

A. **Contact-time matching**. The stationarity conditions (Eq. ES7) give

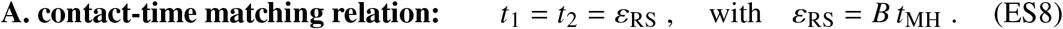 Equivalently, with *t* = *V*/*Q* the same relation gives

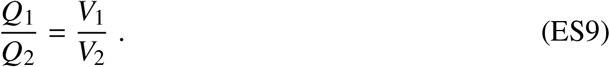

Thus, minimizing structural volume and operational flow, as shaped by natural selection, matches the characteristic contact times of oxygen supply and oxygen use, independently of the detailed geometry of either functional system. The relation of this common contact time to the MH turnover time, *ε*_RS_ = *B t*_MH_, where *B* := *V*_2_/*V* = *V*_blood_/*V* is the relative blood volume, establishes the link between respiratory optimization at the *organismal macroscale*, metabolism at the *MH mesoscale*, and ultimately cellular metabolism and blood rheology at the *microscale*, as derived below.
B. **Respiratory-surface matching**. Substitution of Relation A into Equation ES5 gives

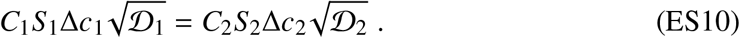 The remaining factors *C*_*i*_Δ*c*_*i*_ describe the local transfer mechanism and the corresponding lateral concentration driving force. Here, Δ*c*_*i*_ denotes the local concentration difference across the exchange surface, distinct from the integral inlet–outlet change 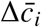 along the main flow direction introduced below (Extended Data Fig. 2). In the coarse-grained two-domain representation, the two local driving differences are referenced to the characteristic tissue–blood concentration scale *c*_0_: Δ*c*_1_ ~ *c*_1_ − *c*_0_ on the oxygen-supply side and Δ*c*_2_ ~ *c*_0_ − *c*_2_ on the oxygen-use side. At high oxygen utilization, *c*_2_ ≪ *c*_0_; the leading-order symmetric case *c*_1_ − *c*_0_ ≈ *c*_0_ − *c*_2_ then gives *c*_1_ ~ 2*c*_0_ and Δ*c*_1_ ~ Δ*c*_2_ ~ *c*_0_. At the same leading-order resolution, we take the dimensionless transfer prefactors to be of comparable magnitude, *C*_1_ ≈ *C*_2_. Thus,

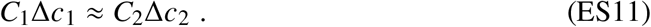

This is a coarse-grained leading-order approximation, not a consequence of the organism-level optimization. Its organism-level consequence is examined using an independently sourced anatomical input in the physiological comparisons that follow. Therefore, we obtain

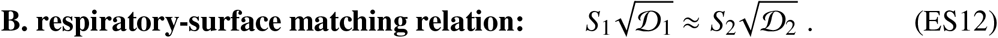

This surface-matching relation links the two exchange domains shown in Extended Data Fig. 2.
C. **Oxygen-content matching**. Relation A already establishes the relation between volume flow and exchange volume. Combining it with the integral organism-level oxygen balance,

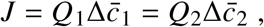

where 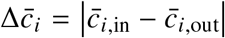 denotes the measurable inlet–outlet oxygen-content change along the flow direction, yields

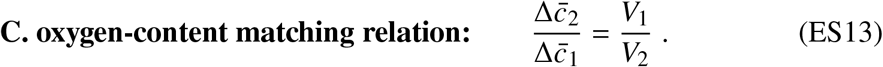

This is a consequence of Relation A and oxygen conservation, not an additional optimization assumption. The integral changes 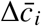 must be distinguished from the local lateral differences Δ*c*_*i*_ used in the mass-transfer relations.

### S2.3 Plausibility tests for a 70-kg human

The organism-level optimization developed above yields three matching relations connecting oxygen supply and oxygen use. To assess their physiological plausibility, we consider a representative 70-kg human at rest. The purpose is not a comprehensive validation of respiratory physiology, but to examine whether the same model-based constrained optimization approach used at the MH mesoscale also yields realistic magnitudes at the organismal macroscale.

We examine the three matching relations through four quantitative comparisons with physiological observations. In each case, an observed quantity on the oxygen-supply side predicts its counterpart on the oxygen-use side: exchange volume and volume flow (Relation A), exchange surface (Relation B), and oxygen-content change (Relation C).

The tests differ in their evidential strength. Exchange surfaces and functional volumes are difficult to identify uniquely with anatomical compartments, whereas volume flows and integral oxygen-content changes are routinely characterized in human physiology. We therefore distinguish agreement in order of magnitude from agreement with more directly accessible physiological quantities.

1. **Test of Relation A: exchange volume**. The functional residual capacity of the human lung is approximately 3 L [73], whereas the total circulating blood volume is approximately 5 L [74]. Taking a representative functional respiratory air volume of 3–5 L, Relation A relates the two structural volumes through their corresponding volume flows. From Equation ES8,

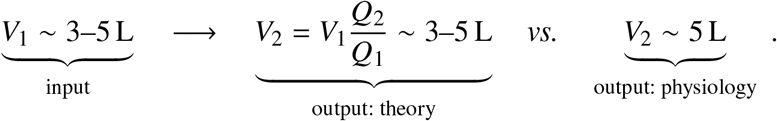

The predicted volume is consistent in magnitude with the observed total circulating blood volume. This agreement is noteworthy because the matching relation follows from the structural– operational trade-off rather than from prescribing equal air and blood volumes. The comparison nevertheless requires a careful interpretation of the functional volumes. The pulmonary air volume varies during respiration, whereas blood is distributed among arteries, capillaries, veins, and cardiac chambers. In particular, the total circulating blood volume is not identical to the volume contained within exchange capillaries. A more precise test would therefore require identifying the effective transport volumes represented by *V*_1_ and *V*_2_ with their anatomical counterparts. At the level considered here, Relation A yields a plausible organism-level volume scale.
2. **Test of Relation A: volume flow**. Resting alveolar ventilation is approximately 4.2–5 L min^−1^ [75], whereas resting cardiac output is approximately 5 L min^−1^ [76]. These quantities represent the operational volume flows through the oxygen-supply and oxygen-use systems, respectively. From Equation ES9,

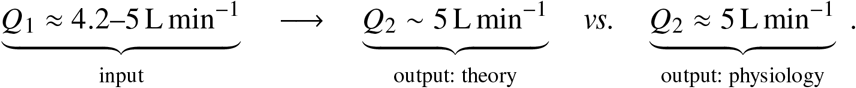

The predicted circulatory volume flow agrees in magnitude with the observed resting cardiac output. Importantly, the approximate equality of the respiratory and circulatory flows is not imposed as an independent matching relation, but emerges from the model under the adopted physiological parameters. Unlike anatomical surface or exchange volume, both volume flows are directly accessible physiological quantities. Their agreement therefore provides a particularly transparent plausibility check of organism-level matching. The comparison applies to the representative resting state; how this matching changes with exercise, altered respiratory conditions, or physiological regulation remains a question for further investigation.
3. **Test of Relation B: respiratory-surface matching**. Classical morphometric studies of the human lung provide an alveolar exchange surface of approximately 70 m^2^ [77]. We use this anatomical reference value independently of the RSA regression of Gillooly *et al*. [41], which serves to constrain the MH topology *N* elsewhere in the theory. With 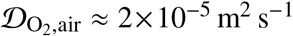 and 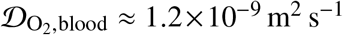 [62, 63], Relation B predicts

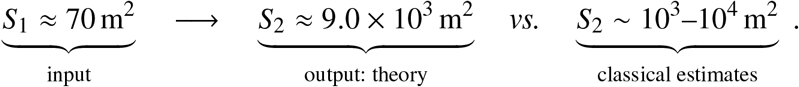

The predicted surface is of the same order of magnitude as classical estimates of the total human endothelial surface, including 4000–7000 m^2^ [78] and 3000–6000 m^2^ [79]. These estimates are indirect, however, and a subsequent reassessment of the underlying anatomical calculations yielded substantially smaller values of 270–720 m^2^ [80]. The comparison is therefore encouraging but not conclusive. In particular, the total anatomical endothelial surface and the effective oxygen-exchange surface *S*_2_ need not be identical. Establishing their precise correspondence, including the role of vascular geometry and the distribution of exchange surfaces, remains an open question. Nevertheless, Relation B produces a physically meaningful macroscopic surface estimate from an independently measured respiratory surface and known molecular diffusivities, without fitting the predicted surface.
4. **Test of Relation C: oxygen-content change**. The local lateral concentration differences Δ*c*_*i*_ entering Equation ES10 are not directly available as whole-organ measurements. The measurable integral changes 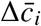 instead test the convective consequence of the same organism-level optimum. At rest, the arteriovenous oxygen-content difference is typically about 5 mL oxygen per 100 mL blood, corresponding to approximately 2.2 mol m^−3^ [81]. A representative value of approximately 2 mol m^−3^ is therefore appropriate for an order-of-magnitude comparison. From Equation ES13,

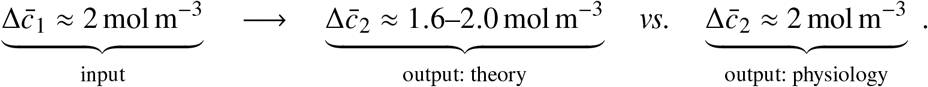

The predicted oxygen-content change is consistent with the physiological arteriovenous difference. This result is also compatible with the integral Fick oxygen balance, which links oxygen consumption, cardiac output, and arteriovenous oxygen extraction. The comparison is therefore a consistency test of the oxygen-content matching relation rather than an independent validation of the local concentration field. In particular, the observed integral changes 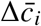 must not be identified with the lateral driving differences Δ*c*_*i*_. A direct test of the local approximation in Equation ES11 would require spatially resolved oxygen-concentration measurements.

The four comparisons assess the three organism-level relations through exchange volume, volume flow, respiratory surface, and oxygen-content change. Despite differences in measurement accessibility and anatomical interpretation, the model yields plausible magnitudes for all four quantities without fitting the predicted values to their observed counterparts.

The surface comparison is subject to substantial uncertainty due to differing anatomical estimates, while the identification of functional exchange volumes requires further investigation. The flow and integral oxygen-content comparisons agree with established physiological observations, although the latter is also constrained by oxygen conservation.

The result supports the broader methodological idea of representing selection and adaptation through model-based constrained optimization: competing pressures to minimize structural resource use and operational effort, subject to fulfilling a prescribed biological function, can lead to matching relations consistent with observed physiological organization. The organism-level analysis complements the MH mesoscale framework and illustrates the potential of this optimization approach for other physiological systems.

### S2.4 Discussion and physical interpretation of respiratory symmorphosis

**Physical meaning of the Pareto parameters—coupling across scales**

The Pareto parameters establish a hierarchical connection across three biological scales. At the mesoscale, Murray’s Pareto parameter *ε*_MH_ governs the optimization of vascular transport within the MH and is constrained by microscale processes, specifically molecular interactions associated with von Willebrand factor (vWF)-mediated platelet adhesion under wall shear. At the macroscale, the respiratory Pareto parameter *ε*_RS_ governs organism-level respiratory optimization, involving respiratory and blood volumes, residence times, and volume flow rates. Its physiological value is determined by the mesoscale MH turnover time *t*_MH_ = *v*/*q* together with organism-level similarity. Thus, microscale physiology constrains mesoscale optimization, which in turn constrains macroscale respiratory organization.

The relative blood volume *B* := *V*_blood_/*V* is defined in the main text, where its order-of-magnitude body-mass invariance within metabolic similarity groups is discussed. Here we use this result to determine the physiological meaning of the organism-level Pareto parameter.

Using Relation A, *Q*_2_ = *Q*_0_ = *N q* = *f*_0_*V*/A, and *V* = *Nv*, we obtain

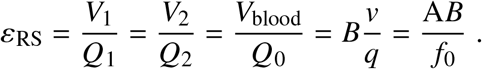

The dimensionless product A*B* = *V*_blood_/*V*_stroke_ is likewise body-mass invariant within a metabolic similarity group under complete similarity. It represents the number of stroke volumes equivalent to the total blood volume. Hence,

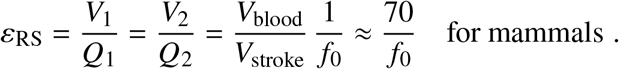

Thus, the organism-level Pareto time is fixed by the MH turnover time and the group-specific relative blood volume. In the Kleiber asymptote, *ε*_RS_ ∝ *v*/*q* ∝ *m*^1/4^. Respiratory contact time and MH turnover time therefore share the same body-mass dependence, differing only by the dimensionless constant *B*.

#### Organism-level matching and local functional reserve

The distinction between the two exchange surfaces is essential: *S*_1_ is the respiratory uptake surface (e.g., alveoli), whereas *S*_2_ is the aggregate systemic blood–tissue exchange surface, represented by *S*_2_ ≈ *N S*/2 in Supplement S5, not the pulmonary capillary surface. Respiratory symmorphosis matches these integrated supply and use capacities; it does not require full utilization of every local exchange unit. In mammalian lungs, blood may approach oxygen saturation within roughly the first third of the pulmonary capillary transit path, leaving substantial local reserve. Such a mesoscale reserve is compatible with organism-level matching of total oxygen supply to use.

The local mechanisms also need not be identical: cyclic gas renewal, plug-flow convection, and continuous laminar boundary-layer transport differ in detail but share the leading square-root transfer dependence established above. This common structure permits one organism-level optimization across air or water uptake and blood-mediated tissue supply without equating the geometries of alveoli, gills, and capillaries.

#### Scope and empirical consequences

Relation A matches contact times and, equivalently, specific volume flows; Relation B matches exchange capacities; Relation C follows from Relation A and the integral oxygen balance. The RSA relation *S*_1_ = RSA and *S*_2_ ≈ *N S*/2 provides the group-specific inference of *N* in Supplement S5. The separate 70-kg comparisons of volume, flow, and inlet–outlet oxygen-content changes test the other implications at order-of-magnitude resolution. Respiratory symmorphosis thus complements, rather than replaces, the mesoscale MH optimization that determines metabolic scaling.

## S3 Kleiber asymptotes and vessel-wall mechanics of the general MH theory

For *v* ≫ *v*_diff_, the following asymptotes are derived from the system of Equations 1–7 by neglecting the *δ*/*d* term within the brackets of Equations 6–7. These relations describe the asymptotic MH geometry underlying the Kleiber regime. Unless stated otherwise, all quantities are MH predictions for *in vivo* conditions.

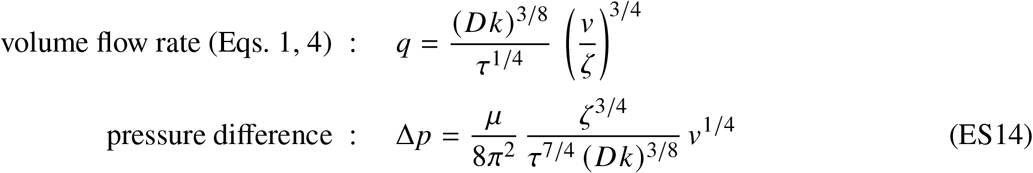

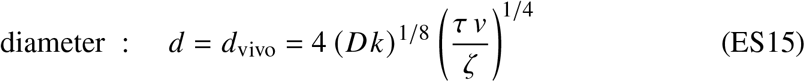

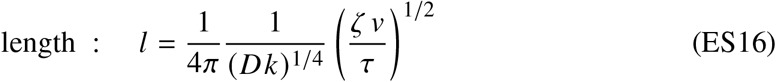

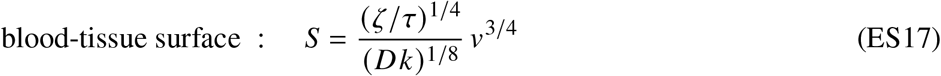

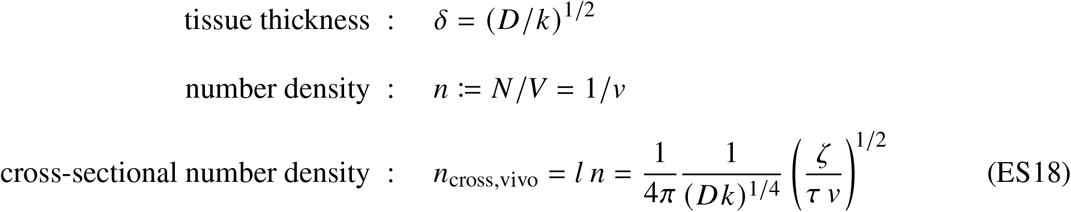

The corresponding dimensionless geometric parameters are:

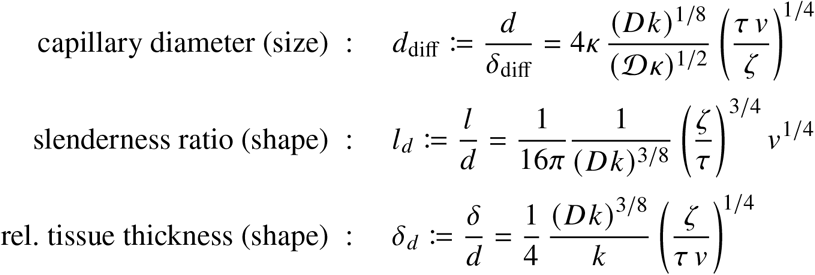

The measures *q, d, l*, and *δ* are solutions of the nonlinear system of Equations 1–7. Quantities related to tissue topology follow directly from MH theory. The pressure scale and the transformation from the predicted *in vivo* geometry to unloaded *in vitro* capillary dimensions are derived below.

### Pressure scaling

The blood pressure drop Δ*p* across the capillary length times the cross-sectional area *A* equals the wall shear stress times the lateral surface of the capillary *S*: 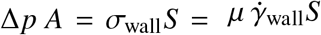. Substitution of 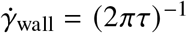, Murray’s law *q* = *d*^3^/(64*τ*), and the Kleiber asymptotic solution recovers Equation ES14. With *v* = *m*/(*ϱN*), the pressure prediction becomes

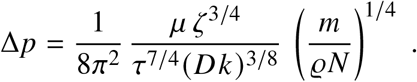

Thus, for *N* ∝ *m*^0^ and body-mass-invariant physiological parameters,

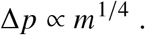

### Vessel-wall mechanics and *in vivo*–*in vitro* conversion

The MH theory predicts the perfused *in vivo* capillary geometry, whereas morphometric measurements may refer to unloaded or weakly pressurized *in vitro* vessels. Their relation is determined here from the nonlinear mechanics of the vessel wall.

Consider a capillary with reference diameter *d*_0_ and collagen-layer thickness *s*_0_. Under a representative transmural pressure *p*, membrane equilibrium gives the circumferential stress

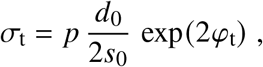

where *φ*_t_ := ln (*d*/*d*_0_)is the logarithmic circumferential strain. The exponential factor 2 accounts for the reduction in wall thickness resulting from incompressibility of the hydrated tissue.

The nonlinear mechanical response of the vessel wall is represented by Fung’s exponential constitutive relation *σ*_t_ = *σ*_0_ exp(*β φ*_t_), where *σ*_0_ = 15–40 kPa is the characteristic reference stress and *β* = 5/2–9/2 is the experimentally observed dimensionless strain-stiffening parameter [82, 83].

We use the representative value *β* = 7/2, the midpoint of this experimental range, throughout the following estimates.

Combining membrane equilibrium with the constitutive relation gives

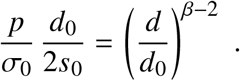

For *β* > 2,

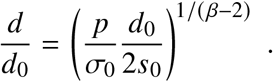

The wall mechanics requires a representative transmural pressure *p*, whereas Δ*p* in Equation ES14 denotes the hydrodynamic pressure drop along the capillary. For the allometric wall-inflation argument, we assume that the body-mass-dependent increment of the representative transmural pressure scales with the hydrodynamic pressure scale Δ*p*, up to a body-mass-invariant proportionality factor. Such a factor leaves the allometric exponents unchanged. For the nominal finite-pressure estimate below, it is taken as unity. We therefore use *p* = *p*_crit_ + Δ*p*, where *p*_crit_ := *σ*_0_ 2*s*_0_/*d*_0_ ~ 1 kPa sets the intrinsic pressure scale associated with the reference vessel wall, consistent with the kPa-scale transmural pressures at which substantial mechanical loading and stress failure of pulmonary capillaries are observed [84].

Identifying *d*_0_ = *d*_vitro_ and *d* = *d*_vivo_ gives the inflation factor

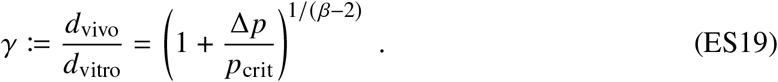

### Finite physiological inflation

Equation ES19 contains both the finite physiological inflation required for absolute morphometric comparisons and the asymptotic body-mass dependence required for allometric scaling. Under physiological conditions, the representative pressure increment is of the order of the intrinsic vessel-wall pressure scale, Δ*p* ~ *p*_crit_. At the nominal value Δ*p* = *p*_crit_, Equation ES19 gives

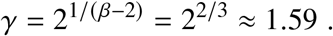

This finite-pressure value is used for the absolute morphometric estimate of *N* in Supplement S5.

### Large-pressure limit and allometric wall inflation

The asymptotic body-mass exponent follows from the large-pressure limit Δ*p* ≫ *p*_crit_, for which the additive unity in Equation ES19 becomes negligible. Combining Equation ES19 with the MH pressure prediction gives

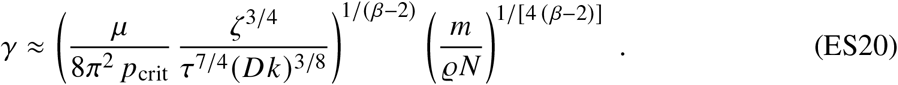

With the representative value fixed above, Equation ES20 then becomes

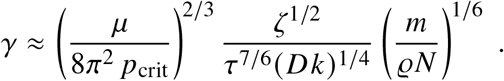

Hence, *γ* ∝ *N*^−1/6^*m*^1/6^, and for *N* ∝ *m*^0^ within a metabolic similarity group,

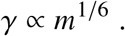

### Allometric scaling of *in vitro* capillary diameter

The MH asymptote predicts *d*_vivo_ ∝ *N*^−1/4^*m*^1/4^. Using *d*_vitro_ = *d*_vivo_/*γ*, together with Equation ES20 gives *d*_vitro_ ∝ *N*^−1/12^*m*^1/12^. For *N* ∝ *m*^0^, this reduces to

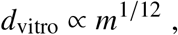

in agreement with the capillary-diameter exponent reported by Dawson [15].

### Allometric scaling of *in vitro* cross-sectional capillary density

The MH prediction refers to the perfused *in vivo* geometry.

In the Kleiber asymptote, Equations ES15 and ES18 imply *n*_cross_ = 4/(*πd*^2^). Hence, with *γ* = *d*_vivo_/*d*_vitro_,

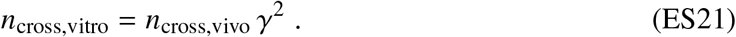

From Equation ES18,

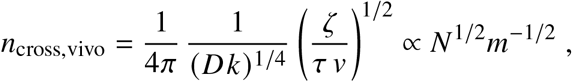

whereas Equation ES20 gives *γ*^2^ ∝ *N*^−1/3^*m*^1/3^. Consequently, the in vitro cross-sectional density scales as *n*_cross,vitro_ ∝ *N*^1/6^*m*^−1/6^.

The full asymptotic expression follows by combining Equations ES18, ES20, and ES21:

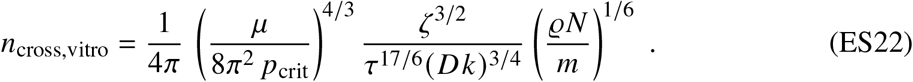

Thus, the within-group similarity prediction *N* ∝ *m*^0^ implies

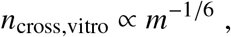

consistent with the capillary-density scaling reported by Schmidt-Nielsen and Pennycuik [25]. Conversely, the observed −1/6 exponent is consistent with body-mass-invariant *N* when the MH *in vivo* prediction is combined with the independently specified vessel-wall mechanics.

## S4 Independent derivation of the MH asymptotes by means of dimensional analysis

### Role of dimensional analysis

The general MH theory is based on model-based *mesoscale optimization* of oxygen transport and yields a general continuous solution spanning the reaction-limited and convective–diffusive regimes, with the corresponding linear and Kleiber asymptotes for different metabolic similarity groups. Here we derive these same asymptotic relations independently by dimensional analysis combined with complete similarity, without solving the MH equations or specifying vascular geometry. The convergence of the two independent derivations provides a stringent theoretical consistency test of the MH theory. Dimensional analysis and complete similarity determine the admissible asymptotic structure, whereas the full MH formulation provides the underlying transport–reaction mechanism, the transition between regimes, and the dimensionless coefficients required for absolute prediction.

We apply the dimensional argument at two levels. First, at the organismal scale, no assumption is made about vascular architecture or about how topology enters the asymptotic coefficient. The dimensionless topology *N* therefore remains an argument of an undetermined dimensionless function. Second, the same analysis is applied to a single MH. Repetition then determines how *N* enters the organismal result. The remaining dependence on oxygen capacity *ζ* cannot be determined by dimensional analysis and is supplied by the full transport–reaction solution of the MH equations.

### Organism-level dimensional analysis without an architectural assumption

Consider the basal whole-organism volume rate within one metabolic similarity group,

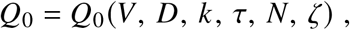

where *V* = *m*/*ϱ* removes the mass scale and leaves a purely kinematic dimensional problem. Buckingham’s Π theorem reduces

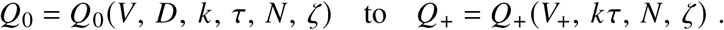

The dimensionless products are

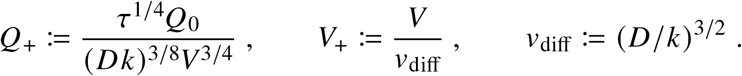

For the Kleiber asymptote, *k τ* is body-mass invariant within a metabolic similarity group and *k τ* ≪ 1. Complete similarity in the limit *k τ* → 0 removes this product from the leading-order relation. For *V*_+_ → ∞, complete similarity with respect to *V*_+_ then gives

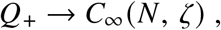

and hence

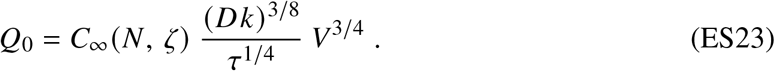

In the opposite reaction-limited branch, convection and its timescale *τ* drop out. In this homogeneous reaction-limited limit, neither vascular architecture nor oxygen-carrying capacity enters the leading-order balance and the leading-order volume rate reduces to *Q*_0_ = *C*_0_ *kV*. The first-order reaction law supplies the remaining physical information and gives *C*_0_ = 1. With Θ = 1, i.e., *k* = *κ*, this gives

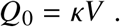

Thus, dimensional analysis combined with the respective asymptotic limits independently recovers both organism-level asymptotes: the linear reaction-limited regime and the 3/4-power Kleiber regime. For the large-organism Kleiber asymptote, dimensional analysis combined with complete similarity therefore independently recovers the 3/4 exponent, but it leaves the dimensionless function *C*_∞_(*N, ζ*) undetermined. In particular, it does not determine how topology or oxygen capacity set the absolute metabolic level.

### MH-level dimensional analysis and repetition

We now apply the same dimensional argument to a single MH, again without specifying its geometry. The relation

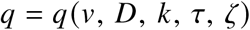

is equivalent to

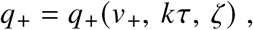

where

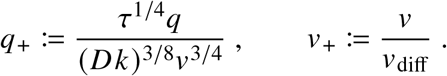

For *v*_+_ → ∞ and *k τ* → 0, complete similarity gives

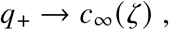

and therefore

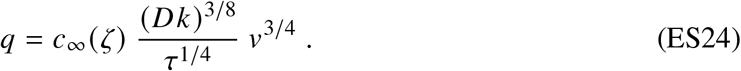

Thus, independently of the explicit MH optimization, dimensional analysis and complete similarity at the level of one functional oxygen-supply domain recover the same mesoscale 3/4-power dependence obtained from the optimized MH equations.

Repetition now provides the kinematic cross-scale mapping,

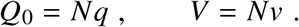

Substitution of *v* = *V*/*N* into Equation ES24 gives

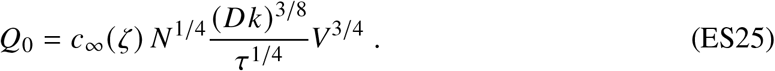

Comparison with the independent organism-level result, Equation ES23, therefore identifies

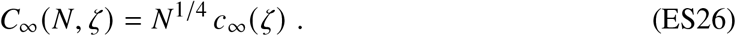

Hence, organism-level dimensional analysis and complete similarity determine the 3/4-power mass dependence without specifying vascular architecture, whereas MH-level dimensional analysis together with repetition resolves the otherwise undetermined topological dependence as *N*^1/4^.

The relations Ė_0_ = *e Q*_0_ in the homogeneous reaction-limited, diffusion-dominated regime and Ė_0_ = *ζ e Q*_0_ in the convective–diffusive Kleiber regime, together with *m* = *ϱV*, convert this kinematic representation into dynamic form.

Dimensional analysis still does not determine the remaining function *c*_∞_(*ζ*). This is the information supplied by the model-based MH optimization. Only the Kleiber asymptote of the general MH theory gives

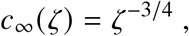

and therefore recovers the complete Kleiber asymptote of the MH theory.

The two routes are thus complementary. Dimensional analysis and complete similarity independently recover the asymptotic mass dependence and, when combined with MH repetition, the topological factor. Mesoscale optimization supplies the physical transport–reaction mechanism, the oxygen-capacity dependence, the absolute coefficient, and the continuous transition between the reaction-limited and Kleiber regimes. Their convergence provides an architecture-independent consistency test of the optimized MH theory.

### Dimensional analysis for heart rate and lifespan

The same dimensional argument applies to basal heart rate and lifespan. For heart rate, the organismal shape factor A = *V*/*V*_stroke_ enters as an additional dimensionless argument,

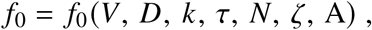

whereas for lifespan the dimensionless empirical closure Ω enters analogously,

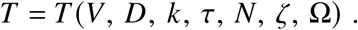

Buckingham’s Π theorem and complete similarity then proceed analogously to the analysis above: A and Ω, being dimensionless, enter only the respective dimensionless asymptotic coefficients, while the resulting mass dependences are *f*_0_ ∝ *m*^−1/4^ and *T* ∝ *m*^1/4^.

## S5 Convergent empirical evidence for the mass-invariance of *N*

The central MH prediction is that the effective number *N* of functional oxygen-supply domains is body-mass invariant within a metabolic similarity group. In vascularized organisms, *N* is represented by the total number of capillary–tissue domains. A direct stereological count across the full body-mass range is unavailable, so the prediction is examined through 11 empirical comparisons spanning structural, geometric, physiological, and organism-level observables (Table 1). Each test probes a different combination of the MH equations; together they cover different organs, species, measurement methods, and six decades of research.

Selected prefactors provide model-based inference (I) of the absolute group-specific magnitude of *N*, scaling exponents provide consistency tests (C) of the prediction *N* ∝ *m*^0^, and downstream observables test predictions (P) after *N* has been specified independently. Such model-based inference is standard in physics: an unobserved quantity is inferred from one set of observables and then constrained by independent consequences of the same model. The inference is therefore overdetermined: conflicts may arise either in the assumed scaling relations or in the absolute magnitude of *N* inferred independently from different observables. The convergence of 11 complementary inference, consistency, and prediction tests in both scaling and magnitude constitutes the empirical evaluation of the topological prediction and strongly constrains any residual dependence of *N* on body mass, without claiming that such dependence can be excluded in principle. Given the convergence of these independent constraints summarized in Table 1, however, a substantial residual dependence of *N* on body mass appears very unlikely. In contrast, the data consistently indicate distinct values of *N* for different metabolic similarity groups.

The tests are analyzed below in the order of Table 1.

### S5.1 Functional oxygen-supply domain number *N* inferred from independent observations

#### No. 1, 2: Respiratory surface area – Gillooly *et al*. (2016)

#### Empirical result

The study by Gillooly *et al*. [41] provides the quantitative basis for inferring the total capillary number *N* in endothermic and ectothermic animals of different body mass. The respiratory surface area (RSA) scales as RSA ∝ *m*^3/4^ with fitted prefactors (Extended Data Fig. 3)

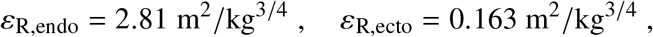

and coefficients of determination 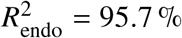 and 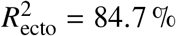.

#### Organism-level symmorphosis inference

The RSA reported by Gillooly *et al*. [41] characterizes the oxygen-supply surface of the respiratory organ. Supplement S2 derives Relation B, the *respiratory-surface matching relation*, which matches this organism-level supply function to the distributed oxygen-use system. For pulmonary RSA, *S*_1_ = RSA and *D*_1_ = *D*_air_. The matched use-side surface is represented by *S*_2_ ≈ *NS*/2, where *S* is the blood–tissue surface of one MH. The factor 1/2 is a coarse-grained model assumption accounting for the effective allocation of MH exchange surface to oxygen use. Thus, *S*_2_ is an aggregate organism-level blood–tissue surface and is not identified with the morphometric pulmonary capillary surface.

The diffusivity correction is essential because the molecular diffusion coefficients of oxygen in the two functional domains—oxygen supply and oxygen use—differ by approximately four orders of magnitude: 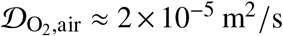 [62, 63] and 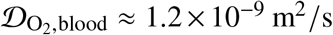 at normal conditions.

#### Application of the *respiratory-surface matching relation* to RSA

Substituting *S*_1_ = RSA and *S*_2_ = *NS*/2 into the respiratory-surface matching relation (Supp. S2, Eq. ES12) gives

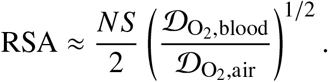

With the lateral capillary surface *S* in the Kleiber asymptote (*v* ≫ *v*_diff_; Supp. S3, Eq. ES17) this yields

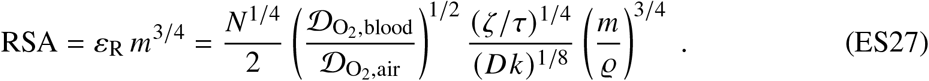

Equation ES27 therefore uses the leading-order relation derived in Supplement S2 in two distinct ways. First, it provides an exponent-level consistency test: the MH-scale relation *S* ∝ *v*^3/4^ follows from the MH equations before any group-specific value of *N* is specified, and with *v* = *V*/*N* the observed scaling RSA ∝ *m*^3/4^ is consistent with *N* ∝ *m*^0^. Second, the measured RSA prefactor supplies the absolute information required to infer the group-specific magnitude of *N*. Thus, RSA tests the predicted mass invariance and anchors the absolute topology without calibration to metabolic-rate data.

#### Inference of *N*

Solving the leading-order Equation ES27 for *N* gives

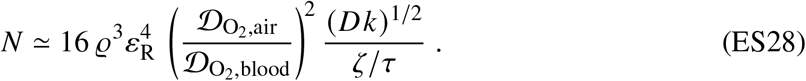

With the physiological parameters of Extended Data Table 1, *ε*_R,endo_ = 2.81 m^2^ kg^−3/4^, and the relative coefficients Ψ = 10^−4^ and Θ = 1 from the CDR solution, Equation ES28 gives *N*_endo_ ≈ 2 × 10^8^. Because the RSA-based inference combines well-constrained data with comparatively robust uncertainty propagation through the MH model, it serves as the anchor for the endotherm topology. It places *N*_endo_ on the 10^8^ scale without calibration to metabolic-rate data.

Estimates from separate observations discussed below provide complementary checks over a broader range, including a whole-organism estimate on the 10^9^ scale. We therefore use *N*_endo_ = 5 × 10^8^, the rounded geometric mean of the RSA-based estimate 2 × 10^8^ and the independent whole-organism estimate 1.2 × 10^9^, as the representative endotherm group value. This value is fixed before the downstream metabolic-rate and heart-rate comparisons and used unchanged as the nominal center of the downstream uncertainty analysis. The relative robustness of the individual inferences is assessed below.

Gillooly *et al*. [41] also reported the respiratory barrier thickness (RBT) for endotherms and ectotherms of different mass. With *ε*_R,ecto_ = 0.163 m^2^ kg^−3/4^ (Extended Data Fig. 3), their central finding RTB_ecto_ ≈ 14 RTB_endo_, independent of body mass, and 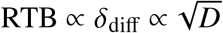, Equation ES28 yields

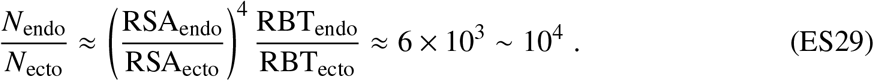

With *N*_endo_ = 5 × 10^8^ this yields *N*_ecto_ = 8 × 10^4^ ~ 10^5^.

The order-of-magnitude ratio, Eq. ES29, is robust: it depends only on the measured RSA prefactors and is insensitive to the precise values of the physiological parameters. Together with the independently constrained group values discussed above, this gives

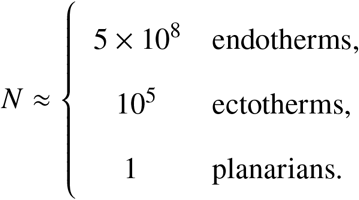

The value *N* = 1 for planarians represents the single-domain limit of the MH framework and sets its natural lower bound. Tests No. 1–2 establish the respiratory-surface magnitude and the robust endotherm–ectotherm topology ratio; together with the independent estimates below, they define the representative group values. These topology values are then fixed and used unchanged in the downstream metabolic-rate and heart-rate comparisons.

### No. 3: Direct capillary estimate and count – Rashevsky (1960)

Data cited by Rashevsky [42] place the total number of capillaries in a dog at approximately *N*_endo_ ≈ 1.2 × 10^9^, based on empirical vascular measurements and branching considerations. This is the only published whole-organism estimate of total capillary number identified in the literature. Together with the RSA-based inference on the 10^8^ scale, it brackets the representative group value *N*_endo_ = 5 × 10^8^ and provides an independent magnitude check (I).

### No. 4: Cross-sectional capillary density – Schmidt-Nielsen & Pennycuik (1961)

#### Empirical result

Schmidt-Nielsen & Pennycuik [25] measured the cross-sectional capillary density *n*_cross_—the number of capillary profiles per unit tissue cross-sectional area—in skeletal muscle of ten mammalian species (bat, mouse, rat, guinea pig, cat, rabbit, dog, sheep, pig, cow) spanning five decades in body mass (Extended Data Fig. 4). Regression of their histological *in vitro* data with the slope fixed at −1/6 gives tissue-dependent prefactors of 6.69 × 10^8^ m^−2^ kg^1/6^ and 1.43 × 10^9^ m^−2^ kg^1/6^ for the two muscle types shown in Extended Data Fig. 4:

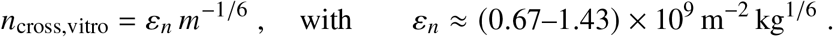

An independent measurement of capillary diameter by Dawson [15] leads to the same capillary-density exponent: *d*_vitro_ ∝ *m*^1/12^ implies 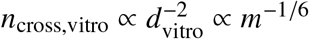.

#### MH prediction

A histological section measures the number of capillary profiles per unit tissue area, *n*_cross_, rather than the volumetric density *n* = *N*/*V*. Within the MH framework, each transport unit of volume *v* and length *l* occupies a cross-sectional area *v*/*l*, so that

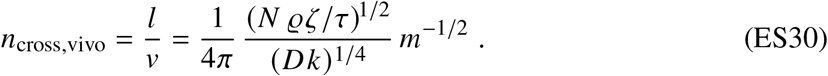

Here, the second equality follows from the Kleiber asymptote derived in Supplement S3.

The conversion from the physiological *in vivo* vessel geometry to the histological *in vitro* cross-sectional density is derived in Supplement S3. There, the MH pressure prediction and nonlinear vessel-wall mechanics determine the inflation factor γ = *d*_vivo_/*d*_vitro_, while the Kleiber asymptote gives

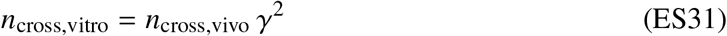

(Supp. S3, Eq. ES21). Combining this relation with the asymptotic wall-inflation scaling yields

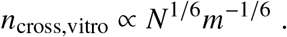

The complete dimensional expression is given in Supplement S3, Equation ES22. Thus, the within-group similarity result *N* ∝ *m*^0^ predicts the observed asymptotic scaling

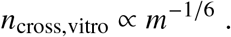

This calibration-free prediction follows from the MH *in vivo* theory combined with independently specified vessel-wall mechanics. The observed −1/6 exponent is therefore consistent with the predicted body-mass invariance of *N* and provides an independent test of the MH framework linking cellular oxygen demand to vascular and tissue-scale organization.

#### Inference of *N*

Equations ES30 and ES31 also permit an independent estimate of the absolute magnitude of *N*. Rearranging gives

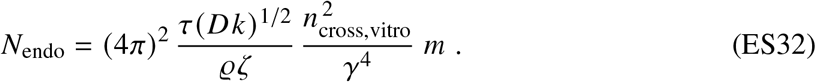

For this magnitude estimate, the finite physiological regime is used, not the large-pressure limit. At the nominal condition Δ*p* = *p*_crit_, the full vessel-wall relation gives γ ≈ 1.59 (Supp. S3). Using this nominal finite-pressure correction and the two tissue-dependent prefactors measured by Schmidt-Nielsen and Pennycuik gives the coarse estimate

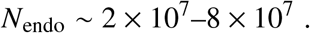

Thus, an independent histological route places the effective MH number in the same broad range as the respiratory-surface estimate *N*_endo_ ≈ 2 × 10^8^ and Rashevsky’s whole-organism estimate *N*_endo_ ≈ 1.2 × 10^9^.

Because *N* depends quadratically on the measured capillary-density prefactor and as γ^−4^ on the inflation correction, this absolute estimate is substantially less precise than the respiratory-surface estimate. Dataset No. 4 nevertheless provides two independent pieces of information: its allometric exponent tests the large-pressure prediction *N* ∝ *m*^0^, whereas its measured prefactor, evaluated with the finite physiological inflation, provides a coarse constraint on the absolute magnitude of *N*.

### S5.2 MH geometry: capillary diameter scaling and absolute values

#### No. 5: *In vitro* capillary diameter scaling – Dawson (2003, 2013)

##### Empirical result

Dawson [15] compiled measurements of capillary diameter across mammals, highlighting the allometric relation

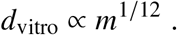

The same exponent is consistent with the cross-sectional capillary-density scaling of Schmidt-Nielsen and Pennycuik (Supp. S5; dataset No. 4).

##### MH prediction and comparison

The MH theory predicts the physiological capillary diameter

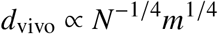

in the Kleiber asymptote (Supp. S3, Eq. ES15). The conversion from the physiological *in vivo* diameter to the histological *in vitro* diameter is derived from nonlinear vessel-wall mechanics in Supplement S3. Its asymptotic result is

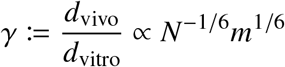

(Supp. S3, Eq. ES20). Hence,

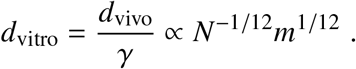

For the within-group similarity result *N* ∝ *m*^0^, this reduces to

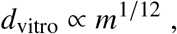

in agreement with the capillary-diameter exponent reported by Dawson [15]. Dataset No. 5 therefore provides an independent exponent-level consistency test of the predicted body-mass invariance of *N* within a metabolic similarity group.

### No. 6: Absolute microvascular geometry – capillary diameter and Krogh radius

#### MH prediction

Once the group-specific topology *N* has been constrained independently, organism size fixes the representative MH volume, *v* = *m*/(ϱ*N*). For prescribed *v, D, k*, τ, and ζ, the closed MH Equations 1–7 determine the MH geometry (*d, l*, δ) and operating state *q* in physical units. Because no parameter is fitted to the microvascular dimensions in this comparison and the group-specific topology *N* is constrained independently, it constitutes a calibration-free prediction of microvascular geometry in physical units.

For a representative 70-kg human, the general MH equations predict an effective capillary diameter of *d* ≈ 11 *µ*m and a tissue transport thickness of δ ≈ 1.0 *µ*m. These dimensions are downstream predictions of the general MH theory in physical units.

#### Krogh radius

The MH tissue thickness δ and the Krogh radius are geometrically distinct quantities. In the MH, δ denotes the effective radial tissue thickness measured outward from the capillary wall (Fig. 2), whereas the Krogh radius represents the characteristic spacing of the tissue-supply domain. It is defined as

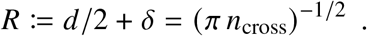

The MH prediction above therefore gives an effective Krogh radius of *R* ≈ 6.7 *µ*m.

#### Human morphometric comparison

Krogh’s classical tissue-cylinder geometry provides the spatial substrate of the Metabolic Holon [18], but anatomy alone does not define the functional unit. Classical morphometric studies of human skeletal muscle report capillary densities of several hundred capillaries per square millimetre [58, 85], corresponding to a characteristic equivalent Krogh radius of approximately *R*_meas_ ≈ 30 *µ*m.

Morphometry of human quadriceps muscle gives a shrinkage-corrected capillary lumen diameter of approximately *d*_meas_ ≈ 5.3 *µ*m [59].

The resulting absolute-scale comparison is therefore

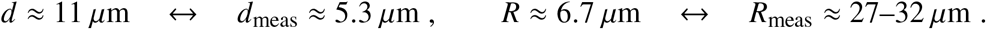

#### Interpretation

Comparison No. 6 provides a cross-scale test that differs fundamentally from the preceding exponent-level comparisons. Starting from organism size, independently constrained topology, and independently specified physical parameters, the MH theory predicts characteristic dimensions at the microscopic scale without fitting a geometric length. The comparison therefore tests whether the coarse-grained theory carries the correct physical scale from the organism level down to the microvasculature.

The two morphometric quantities have different experimental meanings. The measured capillary lumen diameter *d*_meas_ is a direct local histological quantity, whereas *R*_meas_ is an equivalent Krogh radius inferred from cross-sectional capillary density and therefore represents a characteristic tissue-supply spacing. The latter necessarily incorporates local variation in capillary arrangement, perfusion, fibre geometry, metabolic demand, and tissue organization.

This distinction reflects the stated resolution of the MH framework. For a given organism volume *V*, quantities such as *n* and *n*_cross_ describe coarse-grained organism-level organization, while *d, l*, δ, and *R* characterize the corresponding representative MH. Local variations among organs, tissues, or individual muscles are not resolved. Moreover, the MH framework is an order-of-magnitude theory of physical similarity within a metabolic similarity group: it predicts characteristic physical scales rather than tissue-specific histological dimensions.

Accordingly, agreement is to be assessed at the level of characteristic magnitude rather than pointwise histological equality. For human skeletal muscle, the predicted capillary diameter differs from the measured value by a factor of approximately 2.1, whereas the predicted effective Krogh radius is approximately 4.0–4.8 times smaller than the morphometric estimate. Both deviations are smaller than one full order of magnitude and therefore lie within the intended resolution of the theory. Comparison No. 6 should thus be regarded as a test of whether the MH framework predicts the correct microscopic scale, rather than the detailed local geometry of a particular tissue.

### S5.3 Metabolic rate, heart rate, and lifespan

Tests No. 7–10 are downstream prediction tests once the relevant topology has been fixed. Test No. 11 is a partially constrained downstream consequence: the theory predicts *T* ∝ *m*^1/4^, while the empirical lifetime-heartbeat number sets its absolute scale. The predicted metabolic rate (Eq. 8), heart rate (Eq. 10), and lifespan (Eq. 11) are compared with the data of Hemmingsen [22], Thommen *et al*. [17], Seymour & Blaylock [43], Witting [44], and others in Figures 1–3 of the main text. The coefficients of determination are *R*^2^ = 97.7 % (metabolic rate; pooled across all metabolic similarity groups), 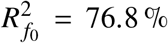 (heart rate), and 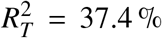 (lifespan). The lower value for lifespan is consistent with the influence of extrinsic mortality factors that are not represented in the metabolic model. The downstream predictions use *N*_endo_ = 5 × 10^8^ and *N*_ecto_ = 10^5^ as the nominal group values, with topology uncertainty propagated according to Extended Data Table 1; the relative diffusion and reaction coefficients are held at the representative values Ψ = 10^−4^ and Θ = 1.

### S5.4 Conclusion

Table 1 separates the epistemic role of each empirical comparison. Respiratory surface area provides the anchor inference: with Ψ = 10^−4^ and Θ = 1, it tests the mass exponent and gives *N*_endo_ ≈ 2×10^8^. The measured respiratory-barrier ratio then constrains the endotherm–ectotherm topology ratio to ~ 10^4^. The remaining routes provide complementary checks based on separate observations and with different uncertainty propagation. Cross-sectional capillary density gives both an independent exponent test and the coarser magnitude estimate *N*_endo_ ~ 2 × 10^7^–8 × 10^7^, whereas Rashevsky’s morphometric estimate gives *N*_endo_ ≈ 10^9^. Taken together, the RSA-based anchor inference and these complementary checks support the representative group values *N*_endo_ = 5×10^8^ and *N*_ecto_ = 10^5^ used unchanged for downstream predictions. Absolute microvascular geometry provides a direct prediction–measurement test of the MH geometry, currently at order-of-magnitude resolution; metabolic rate and heart rate test downstream predictions, while lifespan tests the predicted scaling after its empirical absolute-scale closure. Across all 11 comparisons, no observation contradicts the asymptotic prediction *N* → const ∝ *m*^0^.

#### Direct tests

A body-wide stereological determination of the total number of functional MH capillary domains across a broad mass range would provide a direct test of *N*. Test No. 1 provides an independent reference estimate of *N*_endo_ ≈ 2 × 10^8^ from respiratory surface scaling. Rashevsky gives *N*_endo_ ≈ 1.2×10^9^ for one species, whereas Dataset No. 4 provides the less precise independent estimate *N*_endo_ ~ 2 × 10^7^–8 × 10^7^. The latter is more sensitive to model-input uncertainty, most notably through the γ^−4^ dependence of the inflation correction. Together, these independent magnitude estimates and exponent tests define a quantitative target for direct stereological measurements rather than an adjustable model correction.

## S6 Uncertainty quantification of the MH theory

Deviations between literature data and the presented model responses stem from two different sources of uncertainty: *epistemic uncertainty*, i.e., uncertainty due to incomplete knowledge, and *aleatoric uncertainty*, which is truly random and irreducible, e.g., measurement noise [54].

The focus of this work lies on the model – not on the data. The model is not calibrated to the target observables. Its parameter uncertainty instead reflects uncertainty in independently measured physiological and physicochemical inputs. Thus, the reported uncertainty corresponds to the epistemic part of the uncertainty, which is rooted in incomplete knowledge of the model parameters, together with the physiological deployment variability introduced with the parameter *a*.

Statistical analyses of metabolic-rate observations address a different inference problem: they ask which allometric exponent is supported by the data and show that the exponent is not uniquely determined by the observations alone [6, 12]. Here, by contrast, the mass dependence is derived from the physical model. Probability distributions represent our degree of belief in independently specified model inputs, and the Monte Carlo calculation propagates this input uncertainty to the predicted observables; neither the exponent nor a multiplicative normalization is inferred from the target metabolic-rate data.

This treatment separates physical optimization from robustness assessment. The optimization determines the model state from the governing constraints, whereas uncertainty propagation evaluates the resulting predictions over an ensemble of admissible physiological parameter values. In the terminology of design under uncertainty, performance is therefore assessed beyond a single nominal parameter point without introducing an additional fitting or optimization criterion [54].

To quantify this uncertainty, we adopt a Bayesian interpretation of parameter uncertainty, representing our *degree-of-belief* over the uncertain model parameters by probability distributions.

### Parameter distributions

To obtain meaningful distributions, we derived upper and lower bounds for the parameter values based on literature data and conservative scientific judgment (Extended Data Table 1). We assumed the parameter values to fall between those bounds with high certainty, i.e., 95 % of the probability mass of the distributions should be distributed between those bounds. Additionally, the nominal values listed in Extended Data Table 1 should reflect the expected value of the distributions.

Relating the parameter bounds to the nominal values allows a categorization of the intervals into three different groups:

1. **symmetric intervals on the linear scale** Here, the *difference* between the nominal value and the bounds is equal.
2. **symmetric intervals on the log-scale** Here, the *ratio* between the nominal value and the bounds is equal.
3. **asymmetric intervals** To reflect these properties, the chosen distributional families for the parameter set are the (i) normal, (ii) log-normal, and (iii) skew-normal. An exception is the activity factor *a*, for which a (iv) uniform distribution over the specified interval was selected.

The normal distribution *N* (*µ*, σ^2^) is fully parameterized by its expected value *µ* and variance σ^2^, where σ denotes the standard deviation. As mentioned above, the expected value is equated with the nominal value of the parameter. The standard deviation can be derived from the 95 % bounds [lb, ub] together with the quantiles Φ^−1^(·) of the standard normal distribution *N* (0, 1). Let *X* ~ *N* (*µ*, σ^2^), then *Z* := (*X* − *µ*)/σ ~ *N* (0, 1) and the following hold

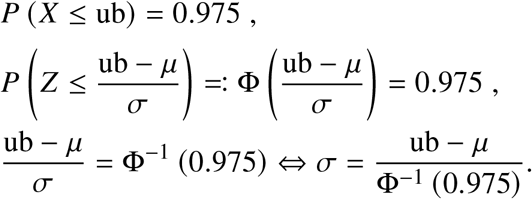

The parametrization of the log-normal distribution, although being defined by two parameters as well, is less straightforward. The location is often specified using the median *m*. Let *X* ~ *N* (*µ*^∗^, (σ^∗^)^2^) and *Y* := exp(*X*). Then *Y* follows a log-normal distribution with median *m* = exp(*µ*^∗^) and expectation *µ* = exp(*µ*^∗^ + (σ^∗^)^2^/2). This yields

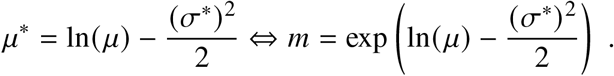

The scale is then often specified using the standard deviation σ^∗^ of the normally distributed counterpart *X*. Due to the log-normal distribution being asymmetrical on the linear scale, both bounds have to be taken into account for the derivation:

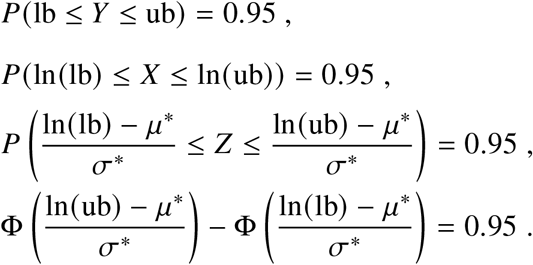

Since *µ*^∗^ = *µ*^∗^(σ^∗^), this requires numerical solving. Additionally, the log-normal distribution is symmetric around its median on the log-scale. Hence, a more natural way is to choose σ^∗^ such that 95 % of the probability mass falls between [*m*/*k, mk*] instead of [*µ*/*k, µk*]:

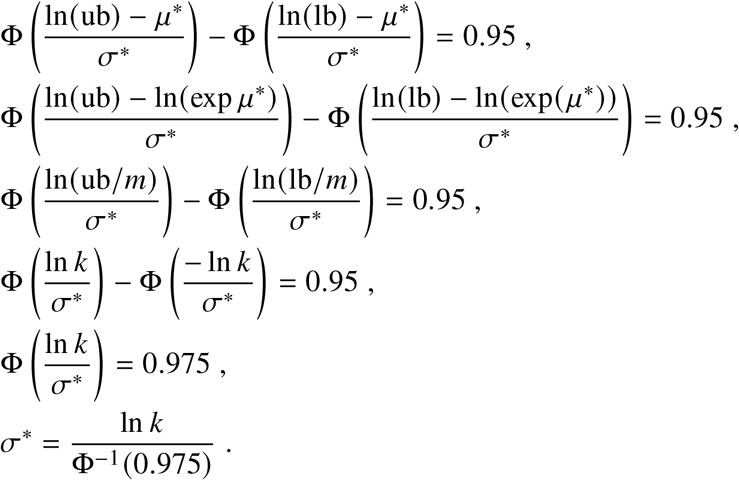

Note that in this way, we preserve the nominal value as the expectation but achieve slightly less probability mass in the initial interval [*µ*/*k, µk*].

The skew-normal distribution SN (ξ, ω, α) requires an additional shape parameter α alongside its location ξ and scale ω. Specifying the expected value *µ* as the nominal value and enforcing 95 % probability mass between [lb, ub] does however only yield two equations for the three unknowns, leaving the problem under-determined. Thus, we impose an additional symmetry constraint to center the expected value inside the interval in the following way:

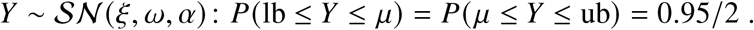

The expected value can be expressed as

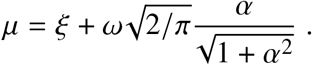

The resulting equation system is then solved numerically.

### Uncertainty propagation

To deduce the resulting degree-of-belief over the response variables, the parameter uncertainties are propagated via Monte Carlo simulations. For these numerical predictions, the relative diffusion and reaction coefficients are held at their representative values Ψ = 10^−4^ and Θ = 1; hence *D* = 10^−4^D and *k* = *κ*, while uncertainty in the effective coefficients enters through the sampled molecular diffusion coefficient D and cellular reaction-rate constant *κ*. For the conditional distribution of the metabolic rate Ė (*m*), this requires repeated solving of the system of Equations 1–7 for a fixed mass *m*, since an explicit expression cannot be derived. To reduce computation time, we parallelized the simulation across the mass grid: each processor core was assigned a fixed mass. Then, for 5 × 10^5^ iterations, samples from the parameter distributions were drawn and fixed in the equation system, which was subsequently solved numerically. The start values for the numerical solver were obtained by linear interpolation of the results from the spreadsheet method introduced in Supplement S10. A fraction ≪ 0.1 % of parameter combinations did not converge, so the realized sample size *n* is correspondingly below the target. The number of iterations was chosen from convergence of the mean and the 95 % interval widths; the convergence analysis is shown in Extended Data Fig. 7.

For each doubling of the sample size, a snapshot of the simulation was stored. The mean and interval width (difference between 97.5 % and 2.5 % quantile) are compared to the previous snapshot by the absolute relative difference and the maximum over all masses is analyzed. The respective maximum relative standard errors for the final sample were 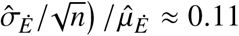% for the mean and ≈ 0.27 % for the interval widths, while the latter was approximated by bootstrapping with 10^3^ repetitions. An exemplary kernel-density estimation for the metabolic rate Ė based on the Monte Carlo simulation is given in Fig. 1b.

The Monte Carlo simulation of the distributions of basal heart rate *f*_0_ and lifespan *T* conditional on body mass is straightforward, because explicit expressions for both quantities are available (Eqs. 10 and 11). Hence, an evaluation at different masses is not necessary and it suffices to perform a Monte Carlo simulation for the coefficients. Due to the equivalence of quantiles under monotone transformations, the 95 % interval for the response variables follows directly from the intervals of the coefficients. The sample size is again chosen to be 5 × 10^5^ for consistency and because similar results for the convergence can be observed.

## S7 Molar mass balances of oxygen – integral form

The oxygen balances developed here and in Supplement S8 form the axiomatic conservation basis of the MH theory. This explicit treatment in both integral and spatially resolved differential form is therefore intentional: the theory ultimately links cellular oxygen consumption to organismal metabolism through conservation of oxygen as it is transported and consumed. Historically, Lavoisier’s work established the central role of oxygen in combustion and respiration and helped place quantitative mass conservation at the foundation of modern chemistry.

In the present theory, the corresponding conservation statement is applied directly to the molar amount of oxygen within a material volume *V* (*t*). The Reynolds transport theorem then transforms this material-volume balance into an equivalent balance over a control volume *V* fixed in space. Mathematically, the theorem is the multidimensional extension of Leibniz’s rule for differentiating an integral with time-dependent integration boundaries.

The molar amount of oxygen within a material volume *V* (*t*) is

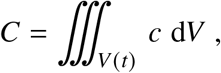

where *c* denotes the local molar oxygen concentration available for transport and, within the tissue, cellular consumption. Conservation of mass gives

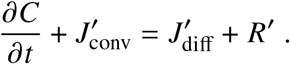

### Convection

The left-hand side contains the local accumulation ∂*C*/∂*t* and the convective molar flow 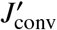. At steady state, the accumulation vanishes and

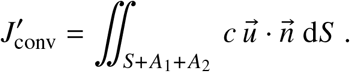

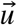 is the blood velocity at 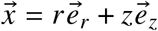, where *r* and *z* are the radial and axial coordinates. *S* = π *dl* is the blood-tissue interface with normal vector 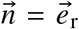 for the capillary volume *v*_cap_ and 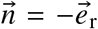 for the tissue volume *v*_tiss_. *A*_1_ = π *d*^2^/4 is the cross-sectional area at the inlet with normal vector 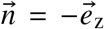 and *A*_2_ is the cross-sectional area at the outlet with normal vector 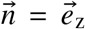. The cross-sectional area of the capillary is constant *A* = *A*_1_ = *A*_2_. The blood volume flow rate through the MH is

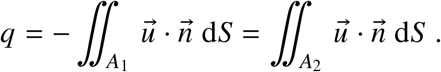

Since there is no convective flow across the blood-tissue interface *S*, the convective molar flow rate of oxygen molecules is

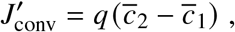

where the flow-weighted cross-sectional oxygen concentration, the mixed-cup concentration, is

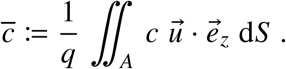

The main text defines the positive oxygen-delivery rate as

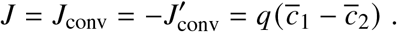

because 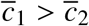.

### Diffusion

The diffusive molar flow rate for the capillary is

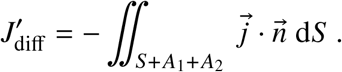

The normal diffusive flux 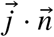 across the inlet and outlet areas *A*_1,2_ is negligible. Fick’s law gives 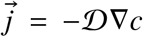, where *D* is the molecular diffusion coefficient. The diffusive molar flow into the capillary volume is therefore

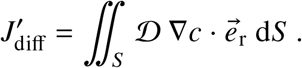

The corresponding diffusive molar flow into the tissue is

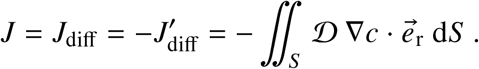

This follows because the normal vector is 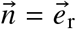 for the capillary and 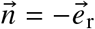 for the tissue.

### Reaction

Cellular respiration is an oxygen sink confined to the tissue. With the volume-specific signed source term *r* = −*κ c*, the signed integral reaction term is

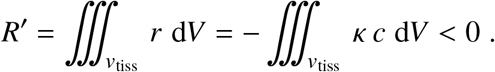

The MH equations use instead the positive oxygen-consumption magnitude *R* := −*R*^′^ > 0. Thus, reaction is physically a sink even though *R* is written as a positive rate in the main-text transport balances. Within the capillary there is no reaction.

### Mass balances

At steady state, the tissue and capillary balances are

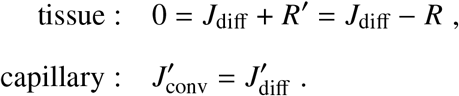

The main text uses this integral formulation as a zero-dimensional, or lumped-parameter, model.

## S8 Molar mass balances of oxygen – differential form

### S8.1 Formulation of the boundary value problem (BVP)

#### Convection-diffusion-reaction (CDR) model

Throughout the CDR formulation, *c* and *c*^′^ denote the local oxygen concentration fields above the residual baseline *c*_res_ ≪ *c*_0_ζ and thus available for transport and, within the tissue, cellular consumption. At the capillary inlet, *c*^′^ = ζ*c*_0_.

The differential oxygen balances follow from the integral balances by Gauss’s theorem:

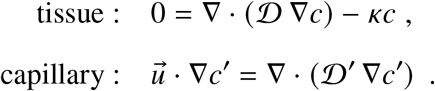

These equations form the 3D convection–diffusion–reaction (CDR) model of the MH. Where necessary, a prime distinguishes the concentration within the capillary from that in the tissue; this notation is otherwise omitted for simplicity.

There is no radial component of the blood velocity field within the capillary, 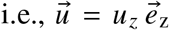. Using the cross-sectionally averaged blood velocity

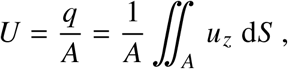

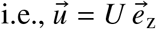, and the homogeneous molecular diffusion coefficient *D* = *D*^′^ = const, the balances become

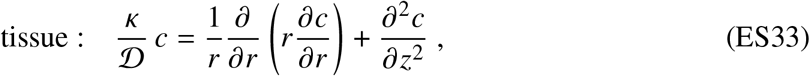

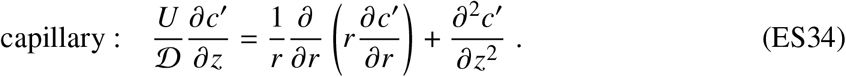

To solve the boundary-value problem, we nondimensionalize the coordinates and geometry. The typical lateral length is the capillary diameter *d* and the typical axial length is the capillary length *l*. Therefore, the transformation is defined by

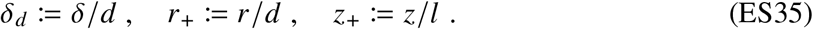

Inserting Equation ES35 into Equations ES33 and ES34 gives

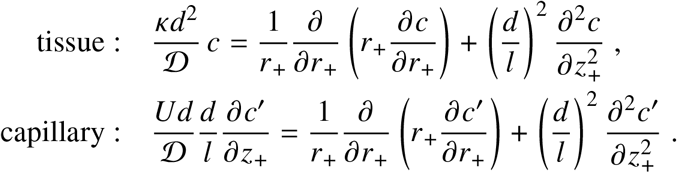

Three dimensionless products characterize this system: (i) the slenderness ratio *l*_*d*_ := *l*/*d*, (ii) the dimensionless capillary diameter *d*_diff_ := *d*/δ_diff_, with δ_diff_ := (*D*/*k*)^1/2^, and (iii) the reduced Péclet number P, derived from the conventional Péclet number Pe := *Ud*/D. The reduced Péclet number controls mass transfer from blood to tissue and is constrained by Murray’s law *U* = *d*/(16πτ). This yields

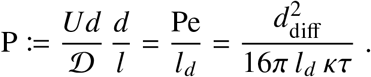

For the physiological geometries considered here, P < 1. Using the Kleiber asymptotes in Supplement S3, the reduced Péclet number becomes

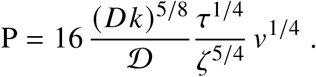

It depends only weakly on body mass (*v* = *m*/(*N* ϱ)).

For the vascular MHs considered here, and in particular in the MH large-domain regime *v* ≫ *v*_diff_, the MH is slender, with *l*/*d* ≫ 1. The CDR system then reduces to

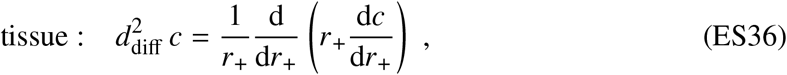

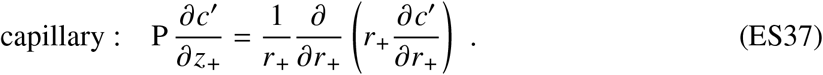

Equation ES33 reduces to the ordinary differential Equation ES36 in *r*_+_ at each axial position *z*_+_; under the cross-sectionally averaged-velocity approximation, the capillary concentration follows from the Graetz-type problem in Equation ES37.

### Boundary conditions

#### Blood-tissue interface

At the blood-tissue interface *r*_+_ = 1/2, both the normal component of the molar flux vector and the oxygen concentration must be continuous:

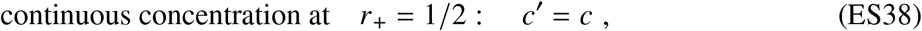

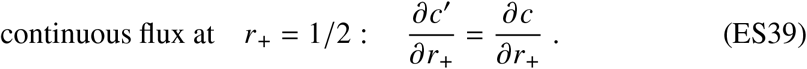

#### Lateral boundary of the MH

At the lateral MH boundary *r*_+_ = 1/2 + δ_*d*_, symmetry imposes the Neumann no-flux condition

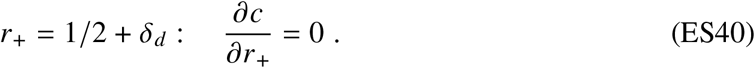

#### Axial boundary of the tissue

For a slender convective–diffusive transport unit (*l* ≫ *d*), the influence of the Neumann boundary conditions ∂*c*/∂*z*_+_ = 0 at *z*_+_ = 0 and *z*_+_ = 1 is confined to an axial region of order δ_*d*_ ≪ *l*_*d*_. These end effects are therefore negligible for slender MHs and are disregarded in what follows; the remaining lateral condition is given by Equation ES40.

#### Boundary conditions for the capillary

At the capillary inlet, *c*^′^ = ζ*c*_0_, and axial symmetry requires

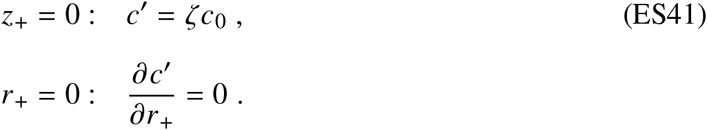

### S8.2 Solution of the boundary value problem (BVP)

We first solve the tissue problem and then the capillary problem. Their coupling produces an eigenvalue problem with discrete roots λ_0_ < λ_1_ < λ_2_ < ….

#### Oxygen concentration field within the tissue

The tissue ODE is the modified Bessel equation of order zero. Its general solution is a linear combination of *I*_0_ and *K*_0_:

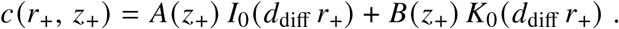

The no-flux condition at *r*_+_ = 1/2 + δ_*d*_ gives

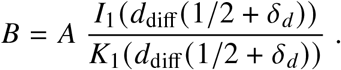

With the radial structure function

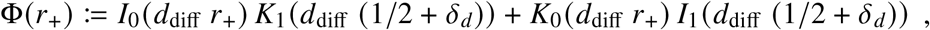

the oxygen concentration field of the tissue is

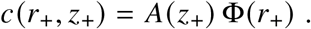

### S8.3 Oxygen concentration field of the capillary

The Graetz problem is solved by separation of variables, with a radial Bessel function *J*_0_ and axial exponential decay:

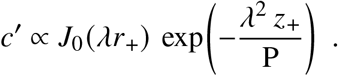

The boundary conditions admit a discrete set of eigenvalues λ_*n*_ (*n* = 0, 1, 2, …). Linear superposition of the corresponding modes gives

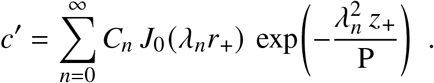

At the fluid–tissue interface *r*_+_ = 1/2, Equations ES38 and ES39 give

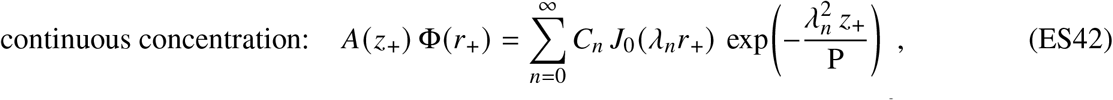

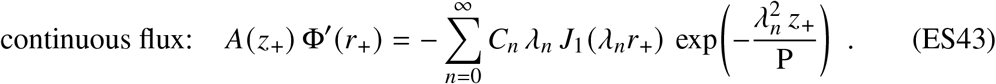

### Dispersion relation determining the eigenvalues λ_***n***_

[Dispersion relation determining the eigenvalues ***λ***_***n***_] Every mode given by an eigenvalue *n* = 0, 1, 2, … has to fulfill the so-called *dispersion relation* derived by the two conditions ES42 and ES43 at the blood-tissue interface:

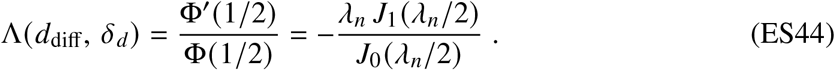

The eigenvalues λ_0_ < λ_1_ < λ_2_ … are the discrete roots of the given Equation ES44.

### Expansion of the inlet condition in a Bessel series

The inlet boundary condition (Eq. ES41) *c*^′^ = ζ*c*_0_ for *z*_+_ = 0 and 0 ≤ *r*_+_ ≤ 1/2 can be expanded into a Bessel series

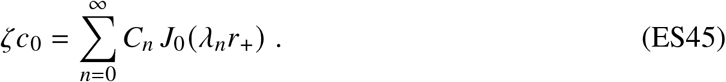

The coefficients *C*_*n*_ of this series are derived using the orthogonality property

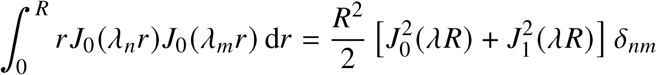

of the Bessel functions. Here, δ_*nm*_ is the Kronecker delta and *R* is the radius.

Therefore, multiplying by *r*_+_*J*_0_(λ_*m*_*r*_+_) and integrating over 0 ≤ *r*_+_ ≤ 1/2 yields the coefficients of the Bessel series

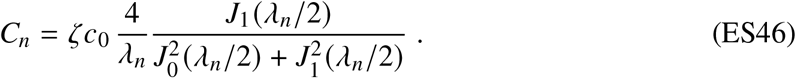

### Axial exponential decay

For known λ_*n*_ and *C*_*n*_, the axial decay of the oxygen concentration is given by the function A(*z*_+_). It is derived from the concentration continuity across the blood-tissue interface:

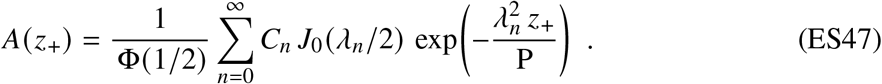

### Complete solution of the CDR problem

Equations ES45, ES46, and ES47 determine the modal amplitudes and axial decay. For clarity, the 3D representation of the oxygen concentration field in both the tissue and capillary is shown here in dimensionless form:

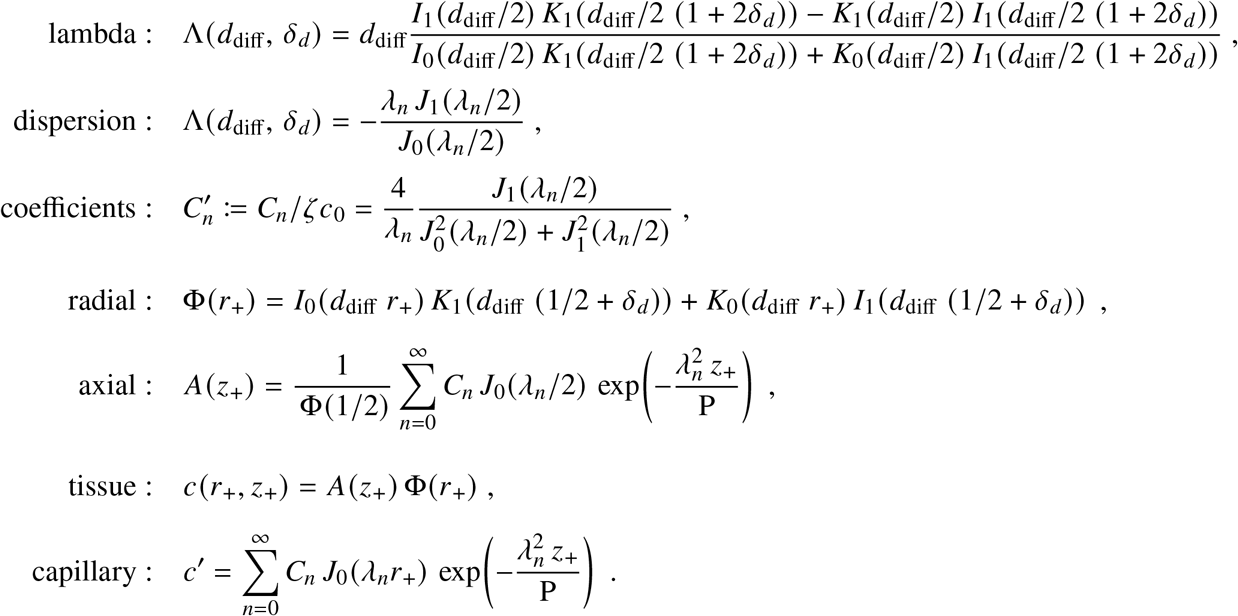

This system is the analytical CDR solution for one MH. A representative dimensional concentration field reconstructed from this solution is shown in Extended Data Fig. 5, emphasizing axial depletion and the radial tissue gradient. The solution is used to determine the effective Nernst diffusion coefficient *D* = ΨD and the effective reaction rate constant *k* = Θ*k* in the following section.

## S9 Effective diffusion and reaction from the three-dimensional CDR solution

The spatially resolved CDR solution derived in Supplement S8 is mapped here onto the 0D MH description. We derive the effective diffusion coefficient *D* and reaction rate constant *k*, establish the exact coupling between the relative diffusion and reaction coefficients Ψ and Θ and the associated MH Damköhler number, and then interpret their numerical values and their influence on MH geometry.

### Effective Nernst diffusion coefficient *D*

Oxygen transport from blood to tissue is represented in the 0D MH balance by an effective Nernst diffusion coefficient *D*, related to the molecular diffusion coefficient by Ψ := *D*/*D*. The coefficient *D* is defined by requiring the Nernst form *J*_diff_ = *c*_0_*DS*/δ to equal the molecular diffusive flux obtained by integrating Fick’s law over the fluid–tissue interface,

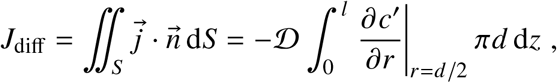

where 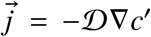 is the molecular diffusive oxygen-flux density and 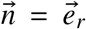 denotes the unit normal directed from blood into tissue. Thus,

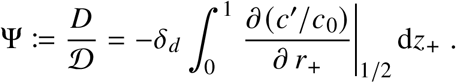

Substituting the capillary concentration field *c*^′^(*r*_+_, *z*_+_) from the CDR solution yields

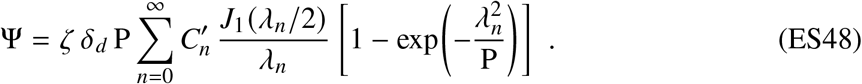

The reduced Péclet number in Equation ES48 is

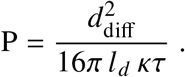

### Effective reaction rate constant *k*

The distributed first-order oxygen consumption within the tissue is represented in the 0D MH balance by an effective reaction rate constant *k*, related to the cellular oxygen-consumption rate constant by Θ := *k*/*k*. Here *R* > 0 denotes the positive oxygen-consumption magnitude used in the main MH equations, i.e., *R* = −*R*^′^ relative to the signed sink term of Supplement S8. The coefficient is defined by requiring *R* = *c*_0_*k v*_tiss_ to equal the volume integral of the positive local consumption magnitude,

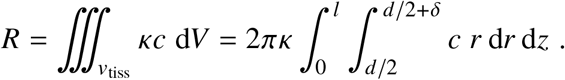

Using the positive consumption magnitude gives

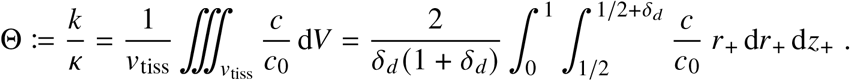

Here 0 ≤ Θ ≤ ζ, where Θ = ζ corresponds to a spatially homogeneous concentration field equal to the inlet concentration ζ*c*_0_ throughout the tissue. Substituting the tissue concentration field *c*(*r*_+_, *z*_+_) yields

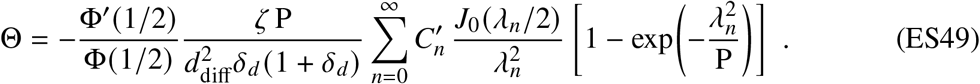

### Coupling of Ψ and Θ and the MH Damköhler number

The two effective coefficients are not independent closures. Because both are integral representations of the same three-dimensional CDR solution, the relative diffusion and reaction coefficients Ψ := *D*/D and Θ := *k*/*k* are constrained jointly. Using the dispersion relation gives exactly

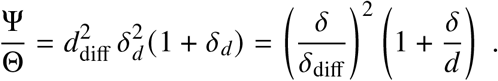

Thus, for a prescribed MH geometry, the ratio Ψ/Θ is fixed by the three-dimensional boundary-value problem. The relative diffusion and reaction coefficients Ψ := *D*/D and Θ := *k*/*k* are derived quantities of the same CDR solution and cannot be prescribed independently.

With *D* = ΨD and *k* = Θ*k*, the same identity becomes

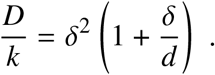

Precisely the same relation follows from the integral 0D MH balance *J*_diff_ = *R*. This equivalence is required by conservation and provides an internal consistency check between the two descriptions. This is because the differential CDR equations follow from the integral balances by Gauss’s theorem; conversely, integration of the three-dimensional solution over the fluid–tissue interface and tissue volume recovers the same integral (0D) balance. The effective coefficients *D* and *k*, and hence Ψ and Θ, therefore preserve exactly the diffusion–reaction balance of the underlying three-dimensional problem.

The same identity can be written as

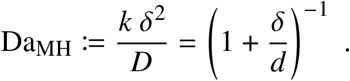

This relation follows from conservation and MH geometry and is not an additional optimization criterion. For the general optimized MH solution corresponding to an organism of given mass within a given metabolic similarity group, Da_MH_ < 1, with its value determined by the geometric ratio δ/*d*. Only in the MH-scale Kleiber asymptote, *v*/*v*_diff_ → ∞ with *v*_diff_ := (*D*/*k*)^3/2^, does δ/*d* → 0, and therefore

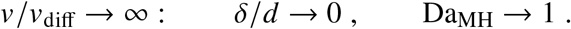

Thus, diffusion and cellular reaction become asymptotically matched in the Kleiber regime. Importantly, Da_MH_ → 1 is not imposed as a criticality condition but follows from the exact diffusion– reaction coupling in this asymptotic limit.

### Asymptotic forms of Ψ **and** Θ

For P ≪ 1, the asymptotic forms of the two relative coefficients are

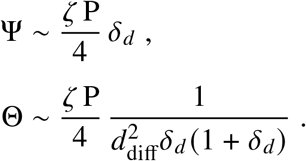

Their product is

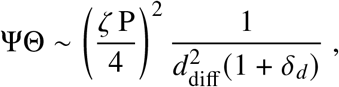

whereas their ratio recovers the exact coupling,

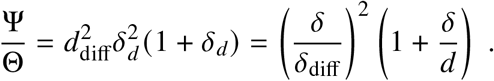

In the Kleiber geometry, δ_*d*_ = δ/*d* → 0, and the exact coupling therefore reduces to

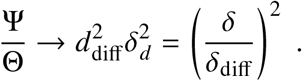

This limit does not by itself determine the separate values of Ψ and Θ. In particular, δ_*d*_ → 0 does not imply Θ → 1: Θ is the tissue-volume average of *c*/*c*_0_ and its numerical value follows from the complete 3D CDR solution. Over the physiological Kleiber domain considered here, the solution remains close to Ψ = 10^−4^ and Θ = 1 (Extended Data Fig. 6).

The absence of a leading explicit size power can nevertheless be seen from the optimized MH asymptotes derived in Supplement S3. Their geometric factors scale as P ∝ *v*^1/4^, δ_*d*_ ∝ *v*^−1/4^, and 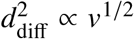. Hence the explicit powers of *v* cancel in both Pδ_*d*_ and 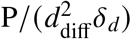. The leading CDR expressions therefore impose no independent power-law dependence of Ψ or Θ on MH volume; any residual variation is contained in the dimensionless CDR solution itself and, for Θ, in the factor (1 + δ_*d*_)^−1^.

### Physical interpretation of Ψ **and** Θ

Neither Equation ES48 nor Equation ES49 contains the topology *N* as an independent argument. Topology enters only when organism mass is mapped onto the MH volume, *v* = *m*/(*N* ϱ), and hence onto the optimized MH geometry. Changing *N* at fixed organism mass therefore changes the absolute scale of one MH, but does not prescribe either relative coefficient independently. Strictly, the dimensionless CDR boundary-value problem depends on both shape variables and dimensionless size variables; Ψ and Θ are therefore not functions of geometric shape alone. What matters for the topology inference is that the coordinated rescaling of the optimized MH introduces no leading explicit power of *v*, as shown by the asymptotic cancellation above.

Extended Data Fig. 6 shows the resulting Ψ(*m*) and Θ(*m*) for the representative group topologies *N* = 1, *N*_ecto_ = 10^5^, and *N*_endo_ = 5 × 10^8^. Their determination is self-consistent: initial values of Ψ and Θ determine the MH geometry, while the corresponding 3D CDR solution yields updated values of the same coefficients. This fixed-point iteration converges over the physiological MH size range, with changes from the initial values indistinguishable at the scale of Extended Data Fig. 6.

The converged solution remains close to Ψ = 10^−4^ and Θ = 1 despite the large variation in MH volume, confirming numerically the leading-order cancellation of explicit MH-scale dependence. Topology enters the CDR evaluation only through *v* = *m*/(*N* ϱ) and is not treated as a tunable parameter.

The relative reaction coefficient Θ has a direct interpretation from its definition,

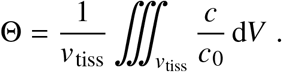

It is the tissue-volume average of the local oxygen concentration relative to the dissolved-oxygen scale *c*_0_. Hence, Θ = 1 means that the volume-averaged tissue concentration remains on the scale of *c*_0_. It does not imply a spatially homogeneous concentration or reaction rate. Because the local first-order reaction rate is proportional to *c*, spatial variation of the concentration directly implies spatial variation of the reaction rate.

The relative diffusion coefficient Ψ has a different physical meaning. The coefficient *D* is defined such that the 0D Nernst relation

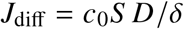

reproduces the diffusive flux integrated over the complete fluid–tissue interface. Thus, Ψ := *D*/D does not measure a change of the molecular diffusivity itself. Rather, it measures how the spatially heterogeneous three-dimensional diffusive transport is represented by the 0D Nernst form using the complete interface *S* and the characteristic distance δ.

The spatial CDR solution explains why this relative coefficient can be small. The strongest wall-normal gradients occur where the capillary oxygen concentration is high, whereas axial oxygen depletion reduces the driving gradient further downstream. Strong local gradients can therefore coexist with a much smaller coefficient after the transport over the complete interface is represented by a single 0D quantity.

The exact coupling provides a complementary geometric interpretation,

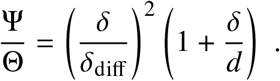

In the physiological MH domain represented in Extended Data Fig. 6, δ ≪ δ_diff_. The strong separation Ψ ≪ Θ therefore reflects the separation between the locally supplied tissue thickness δ and the molecular reaction–diffusion length δ_diff_. Together with Θ = 1, this explains why the CDR solution yields the relative diffusion coefficient Ψ several orders of magnitude below unity.

## S10 Spreadsheet method for solving the system of MH equations

Equations 1–7 form a nonlinear system that is solved by Newton–Raphson iteration or by the tabulation below.

### Asymptotes

The two limits have analytical solutions. For small organisms, *d*/δ ≪ 1, the first term in the brackets of Equations 6 and 7 is negligible. In this limit, *d*, τ, and *q* drop out, yielding the reaction-limited asymptote given in the main text; the MH reduces continuously to the nonvascular diffusion–reaction regime.

For large MHs, *d*/δ ≫ 1, the second term is negligible in each case. The tissue thickness approaches the diffusion limit, 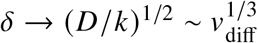.

### Tabulated solution of the MH equations

The nonlinear system can also be solved by tabulating the following columns over *d*/δ.

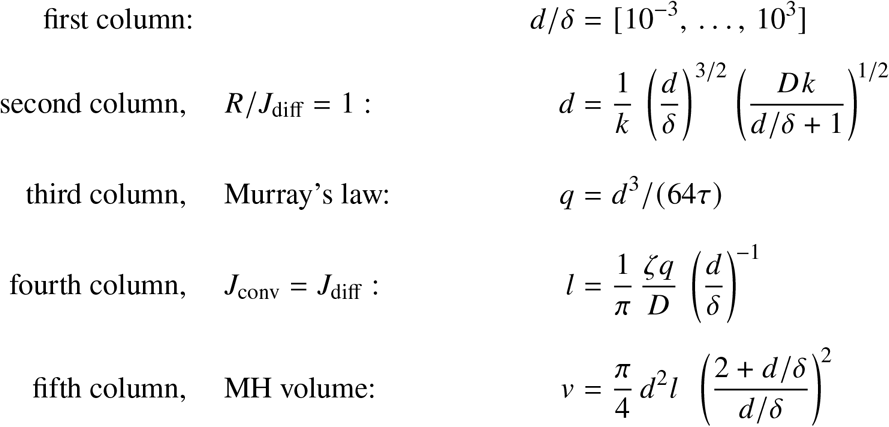

## S11 Scope and organizational implications of the MH frame-work

The reduced MH model retains the dominant transport, reaction, geometric, and topological relations relevant to the predictions tested here. Organ heterogeneity, transit-time distributions, nonlinear binding kinetics, and regulatory feedback introduce additional spatial and temporal variation that is not resolved by this leading-order description. Figure 2 summarizes the cross-scale organization and the information entering the MH mesoscale.

### Organizational consequences of repetition

Repetition has consequences beyond metabolic scaling. Repeated functional units can localize failure and provide a basis for redundancy: where neighboring or connected units compensate for one another, loss of one unit need not compromise the whole system. Repetition may therefore promote resilience while allowing optimized mesoscale functional units to constitute the larger organismal system.

Diversity acts at a different organizational level. Even within a metabolic similarity group, organisms need not be identical: similarity constrains the leading coarse-grained dimensionless organization while leaving room for substantial variation in its biological realization. Such variation provides alternative realizations on which selection can act, whereas repetition builds the organismal whole from an already established mesoscale functional organization. Changes in the effective topology *N* may consequently provide a signature of evolutionary transitions in functional organization. The MH framework, however, does not explicitly model the generation, selection, or diversification of evolutionary variants.

### Architectural complexity

A system built from many repeated MHs may be architecturally complicated while its leading metabolic behavior remains governed by the transport–reaction physics of one optimized mesoscale functional unit. The vascular distribution system remains essential for connecting the MHs and regulating flow among them; its detailed architecture can affect heterogeneity, regulation, robustness, and higher-order transport behavior beyond the leading-order description developed here.

Repetition and diversity therefore serve distinct but complementary roles in biological organization. Repetition builds the organismal whole from optimized mesoscale holons. Diversity, including variation within similarity groups, provides alternative realizations on which selection can act and from which evolutionary innovation can emerge. In this sense, holonic organization provides a possible route by which simple mesoscale organization can coexist with large organismal scale and biological diversity.

## S12 Evolutionary interpretation of the MH framework

The MH theory describes transport, reaction, geometry, and organismal organization through physical laws and constrained optimization. The three competing pressures at the MH mesoscale arise from resource scarcity and functional demand, subject to physicochemical constraints. The optimized MH is interpreted as a possible outcome of selection and adaptation under these pressures, not as a prescribed evolutionary trajectory. Deviations from the predicted optimum may arise when historical, developmental, or additional functional constraints modify the accessible design space (Fig. 4).

Optimization and innovation play fundamentally different roles. Selection and adaptation may favor preferred states within an existing design space, whereas innovation changes its boundaries, allowing previously inaccessible physical solutions. *Optimization moves the point; innovation moves the boundaries—and thereby the space itself*. Organism-level optimization (Supp. S2) remains distinct from mesoscale MH optimization and does not generate the metabolic scaling law.

The framework thus connects a Newtonian description of physical constraints with a Darwinian interpretation of selection, adaptation, and innovation [21]. It does not model mutations, population dynamics, fitness landscapes, or evolutionary histories. Instead, it identifies physically optimized states and provides a design space within which evolutionary hypotheses can be formulated and tested.

